# Spatiotemporal analysis of gene expression in the human dentate gyrus reveals age-associated changes in cellular maturation and neuroinflammation

**DOI:** 10.1101/2023.11.20.567883

**Authors:** Anthony D. Ramnauth, Madhavi Tippani, Heena R. Divecha, Alexis R. Papariello, Ryan A. Miller, Erik D. Nelson, Elizabeth A. Pattie, Joel E. Kleinman, Kristen R. Maynard, Leonardo Collado-Torres, Thomas M. Hyde, Keri Martinowich, Stephanie C. Hicks, Stephanie C. Page

**Affiliations:** Lieber Institute for Brain Development, Johns Hopkins Medical Campus, Baltimore, MD, USA; The Solomon H. Snyder Department of Neuroscience, Johns Hopkins School of Medicine, Baltimore, MD, USA; Cellular and Molecular Medicine Graduate Program, Johns Hopkins School of Medicine, Baltimore, MD, USA; Department of Psychiatry and Behavioral Sciences, Johns Hopkins School of Medicine, Baltimore, MD, USA; Department of Biostatistics, Johns Hopkins Bloomberg School of Public Health, Baltimore, MD, USA; Department of Neurology, Johns Hopkins School of Medicine, Baltimore, MD, USA; Johns Hopkins Kavli Neuroscience Discovery Institute, Baltimore, MD, USA; Department of Biomedical Engineering, Johns Hopkins School of Medicine, Baltimore, MD, USA; Center for Computational Biology, Johns Hopkins University, Baltimore, MD, USA; Malone Center for Engineering in Healthcare, Johns Hopkins University, Baltimore, MD, USA

**Keywords:** hippocampus, neurogenesis, aging, inhibition, extracellular matrix, glia

## Abstract

The dentate gyrus of the hippocampus is important for many cognitive functions, including learning, memory, and mood. Here, we investigated age-associated changes in transcriptome-wide spatial gene expression in the human dentate gyrus across the lifespan. Genes associated with neurogenesis and the extracellular matrix were enriched in infants, while gene markers of inhibitory neurons and cell proliferation showed increases and decreases in post-infancy, respectively. While we did not find evidence for neural proliferation post-infancy, we did identify molecular signatures supporting protracted maturation of granule cells. We also identified a wide-spread hippocampal aging signature and an age-associated increase in genes related to neuroinflammation. Our findings suggest major changes to the putative neurogenic niche after infancy and identify molecular foci of brain aging in glial and neuropil enriched tissue.

## 1 Introduction

The dentate gyrus (DG) of the hippocampus (HPC) is important for many cognitive processes ^1–6^, which mature and decline over development and aging ^7^. For example, the ability to form long term memories does not begin until early childhood ^8,9^, while cognitive decline begins in late adulthood ^10–12^ and accelerates in age-related dementias such as Alzheimer’s disease (AD) ^13,14^. The development and deterioration of these cognitive functions is partially regulated by molecular, cellular and anatomical processes that change across infancy, adolescence, and adulthood in the DG^15,16^. The DG contains distinct spatial domains due to the unique organization of its specialized cell types. Dentate granule cells (GCs) are asymmetrical glutamatergic neurons organized in a densely packed granule cell layer (GCL) whose dendrites comprise the molecular layer (ML). A thin subgranular zone (SGZ) separates the GCL from the cornu ammonis 4 (hilus/CA4), which connects the DG to the remainder of the HPC and relays information from CA3 to CA1 to the subiculum. The GCs in the GCL receive major excitatory inputs from the entorhinal cortex, and are specialized to regulate information flow into the HPC ^1,2,17^. The spatial heterogeneity of tissues results in regions dense with cell bodies of varying cell types in varying states, or regions sparse in cell bodies but enriched with synapses, termed neuropil. Importantly, this spatial heterogeneity can be captured with spatially-resolved transcriptomics.

In most mammalian species that have been studied, a spatially-defined neurogenic niche is retained in the adult DG. In line with this function, populations of proliferating radial glial-like stem cells reside in the SGZ, with processes that extend into the ML. These stem cells differentiate and migrate as neuroblasts into the GCL during their developmental trajectory into mature dentate granule cells ^18^. While adult neurogenesis in the rodent is well-established, the existence, abundance, and function of adult neurogenesis in the human DG remains controversial ^19^. Cell types representing the full developmental trajectory of neurogenesis (radial glial-like stem cells, migrating neuroblasts, immature granule cells) have been identified and profiled using single-cell RNA sequencing (scRNA-seq) in mouse HPC ^20^. Similarly, single-nucleus RNA sequencing (snRNA-seq) in macaque HPC identified all major neural precursor populations, but comparisons to mice suggested differences in neurogenic processes ^21^. While some immunohistochemical studies support the existence of adult neurogenesis in human HPC ^22–30^, recent snRNA-seq studies suggest an absence or paucity of newborn neurons ^31,32^. However, since snRNA-seq studies lack spatial context and do not capture extranuclear transcripts, findings regarding adult neurogenesis at the human SGZ remain inconclusive.

The DG also undergoes dynamic changes with aging, and is particularly vulnerable to the neurotoxic effects of “inflammaging” - an age-related increase in pro-inflammatory markers ^33^. Inflammation is associated with blood-brain barrier (BBB) permeability, and human functional magnetic resonance imaging (fMRI) studies suggest pronounced aging-related increases in BBB permeability at the DG ^34^. In aged rodents, neuroblasts adopt a senescence-associated secretory phenotype that induces auto-inflammation ^35^, while glial cells, including microglia and astrocytes, undergo age-dependent increases in neuroinflammation in the DG, which decreases neurogenesis and exacerbates degeneration ^33,36–44^. Importantly, activated microglia can induce reactive astrocytes ^39^, triggering further inflammation and neurodegeneration. However, microglial molecular profiles which have been characterized in mouse models cannot fully recapitulate the transcriptomic phenotype of aged human microglia ^45^, highlighting the importance of human post-mortem studies. Again, as with neurogenesis, many molecular profiling studies lack spatial resolution in characterizing regional vulnerabilities to inflammaging.

To better address age-based changes to the neurogenic niche and regional vulnerability to “inflammaging,” we deployed SRT in the human DG, which allowed us to simultaneously profile DG sub-domains enriched and depleted in cell bodies. By profiling intact tissue sections, spatially-resolved transcriptomics (SRT) provides gene expression information within the existing architecture, and captures extranuclear transcripts. In this study, we investigated changes in transcriptome-wide spatial gene expression in the human DG across four stages of the lifespan (infant, teen, middle-age and elderly). We performed differential expression analyses across age groups at the level of the composite DG as well as individual DG sub-domains. Here, we show decreased neural proliferation and protracted development of inhibitory neurons and oligodendrocytes after infancy, and accelerated age-related changes associated with neuroinflammation in glia-rich areas. To further explore spatio-molecular heterogeneity in the DG across the human lifespan, we provide full access to the data, code, and an interactive web application (https://libd.shinyapps.io/Lifespan_DG/).

## 2 Results

### 2.1 Experimental design to investigate spatial gene expression in the human dentate gyrus across the lifespan

We dissected tissue blocks from coronal slabs of fresh-frozen postmortem human brains at the level of the anterior HPC from 17 neurotypical donors (ages 0.18 to 76.7 years, **Table S1**). We binned donors into four age groups representing key developmental timepoints: infant (0-2 years), teen (13-18 years), adult (30-50 years), and elderly (70+ years) to investigate spatio-molecular signatures across the lifespan. Each block was scored to isolate the DG, and one cryosection per donor was mounted on a 10x Genomics Visium slide (*N*=17 total capture areas, **Figure 1a**). Following image acquisition and sequencing, we assessed library quality and confirmed DG inclusion on each capture area using standard quality control (QC) methods (**Figure S1, Figure S2**). A capture area from one elderly donor (Br3874) was removed because it did not contain the GCL (**Figure S2**, **Figure S3**), and hence the final dataset included *N*=16 capture areas with a total of 68,685 spots. Data preprocessing included normalization and feature selection, as well as integration with `Harmony` *^46^*to correct for batch effects (**Figure S4**).

**Figure 1:**
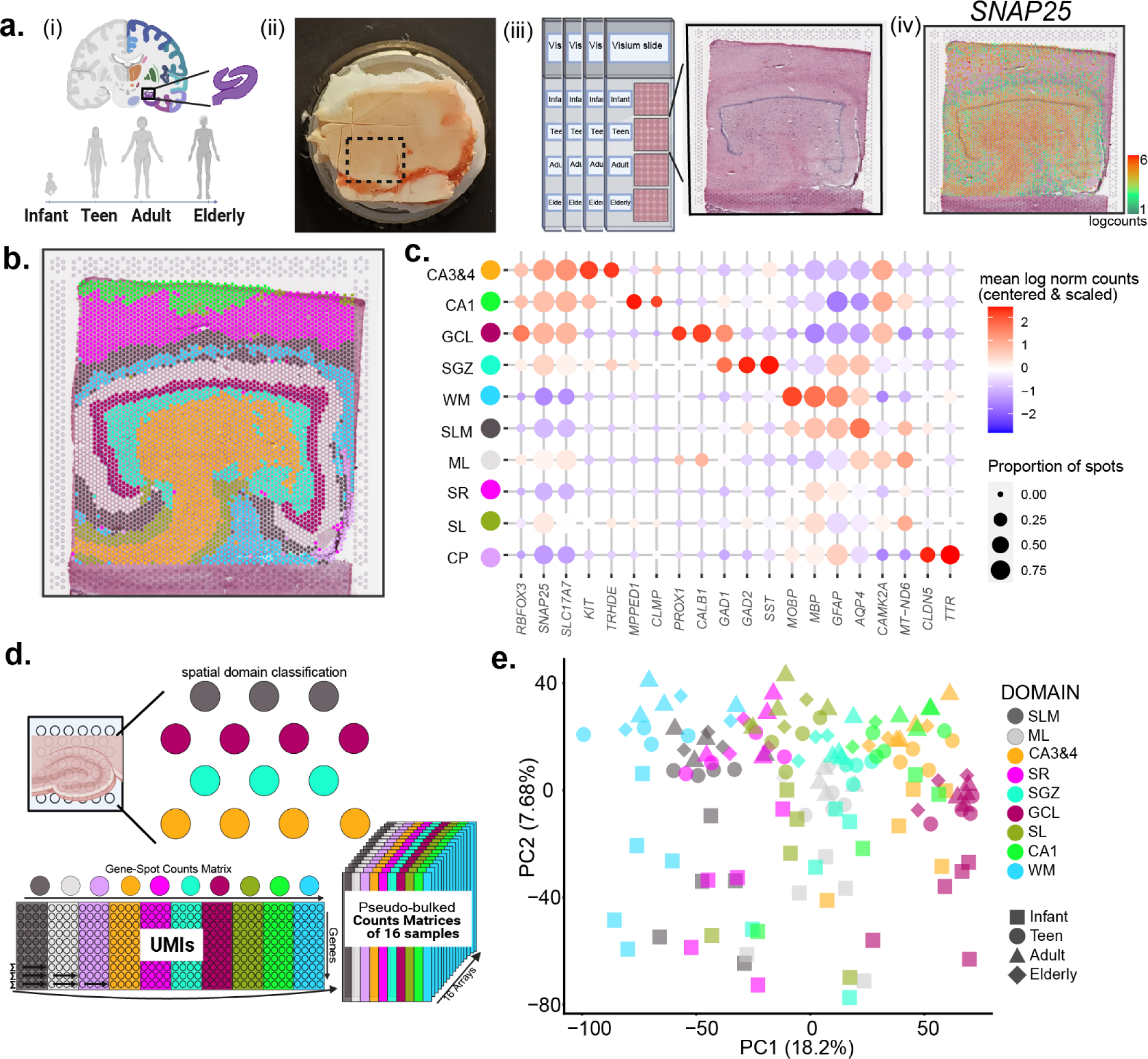
Spatially-resolved transcriptomic profiling and unsupervised spatial domain detection in the human dentate gyrus across the postnatal lifespan. **(a)** (i) Experimental design for acquiring SRT data in the human dentate gyrus (DG) across four age groups: infant (0-2 years), teen (13-18 years), adult (30-50 years), and elderly (70+ years). (ii) Tissue blocks were dissected from frozen brain slabs and scored to isolate the DG, then (iii) cryosectioned at 10μm, mounted onto Visium slides (10x Genomics), and stained with hematoxylin and eosin (H&E). (iv) On-slide cDNA synthesis was followed by library construction and Illumina sequencing, providing gene expression levels (logcounts) at X-Y spatial coordinates as illustrated by representative spot plot for *SNAP25*. (b) DG from donor Br1412 with Visium spots labeled by spatial domains predicted by `BayesSpace` at *k*=10. (c) Dot plot for domain-specific and neuropil-enriched gene markers for the 10 spatial domains. Dots are sized by the proportion of spots with nonzero expression and colored by mean log_2_ normalized counts, centered and scaled. Abbreviations: CA3&4 ≅ cornu ammonis 3 & 4, CA1 ≅ cornu ammonis 1, GCL ≅ granule cell layer, SGZ ≅ subgranular zone, WM ≅ white matter, SLM ≅ stratum lacunosum moleculare, ML ≅ molecular layer, SR ≅ stratum radiatum, SL ≅ stratum lucidum, CP ≅ choroid plexus. (d) Strategy for pseudo-bulking spots across spatial domains for each of the *N*=16 capture areas. (e) Scatter plot of the first two principal components (PCs) after pseudo-bulking spots for each sample and spatial domain (excluding domain 3, which maps choroid plexus). Data points are labeled by color (spatial domain) and shape (age), with percent variance explained in parentheses.

To identify spatial domains, we performed unsupervised spatial clustering using `BayesSpace` as previously described ^47,48^. Using *k*=10 to predict spatial domains (**Figure 1b, Figure S5**), we found canonical domain-specific gene markers expressed (**Figure 1c, Figure S6**). For example, domain 7 is enriched for GCL markers *PROX1* and *CALB1*, and domain 6 is enriched for interneuron markers characteristic of the subgranular zone (SGZ) including *GAD1*, *GAD2*, and *SST*. Domain 4 is enriched for markers of both CA3 and CA4 including *KIT* and *TRHDE,* and thus was classified as ‘CA3&4’ (**Figure 1c, Figure S6**). Domain 2 is enriched for synaptic markers such as *CAMK2A* and mitochondrial gene *MT-ND6*, which is characteristic of the ML, a domain primarily composed of neuropil (**Figure S5**, **Figure S6**). Independently, we assigned spots to canonical HPC sub-regions using H&E histology and known marker gene expression (**Figure S7**). Comparison of predicted spatial domains to manual annotations using principal components (PCA) and Uniform Manifold Approximation and Projection (UMAP) ^49^ visualization revealed less intermingling of spot cluster labels in expression space for predicted domains, suggesting overclustering and/or mixing of cluster labels at borders of spatial domains by manual annotation (**Figure S8a-c, Table S2**). We also evaluated the quality of the predicted spatial domains using cluster purity ^50^, and found higher purity in predicted spatial domains compared to manual annotations (**Figure S8d**). Given the more precise delineation of spatial domains following unsupervised clustering compared to manual annotations, we proceeded with predicted spatial domains for all downstream analyses.

To identify differentially expressed genes (DEGs) across spatial domains, we pseudo-bulked Visium spots by summing the UMI counts of each gene for each capture area across spots within a spatial domain (**Figure 1d**). Using the 160 pseudo-bulked samples (16 capture areas x 10 spatial domains), we performed PCA, and found that spatial domain 3, which mapped to the choroid plexus (CP) (**Figure 1c, Figure S6**), explained a large fraction of variation (**Figure S9a**). Given that CP is a secretory tissue that produces cerebrospinal fluid and is not a domain related to the HPC, we removed CP from further downstream analysis. PCA without CP illustrated that spatial domain, followed by age group, were the top components of variation. Infant samples separated from all other age groups (**Figure 1e, Figure S9b**), highlighting a major developmental distinction associated with early life.

### 2.2 Identification of age-associated molecular signatures in the human dentate gyrus

Using the pseudo-bulked data, we further combined the ML, GCL, SGZ, and CA3&4 spatial domains *in silico* to generate a composite DG, which reduced sparsity and increased UMI coverage of genes for age-based comparisons (**Figure 2a**). We included CA3&4 in the composite DG because, in humans, this region contains mossy cells and interneurons as well as CA3 pyramidal neurons ^15^. Some tissue sections contained other HPC domains, but since they were not equally represented across donors (**Figure S5**, **Figure S10**), they were not included in these composite DG analyses. We performed differential expression (DE) analysis on the pseudo-bulked data to compare the composite DG in each age group to all other age groups (**Table S3**). We identified the most DEGs in the infant group (*n*=3284), followed by the elderly group (*n*=1745). Fewer significant DEGs with smaller fold changes were identified in teen (*n*=462) and adult (*n*=834) groups (**Figure 2b-e**). Interestingly, we noted a dichotomy between adult/elderly and infant/teen with respect to genes associated with proliferation and neuroinflammation. Specifically, some genes associated with neurogenesis were significantly upregulated only in infancy, while genes associated with activated microglia and reactive astrocytes were downregulated in the infant and teen groups (**Figure 2b-c**), but upregulated in the adult and elderly groups (**Figure 2d-e**). This pattern suggests an inverse relationship between proliferative potential and glial inflammatory activity that changes with onset of adulthood. We also noted depletion of some inhibitory neuronal marker genes in the infant group, which were significantly enriched in the adult group; specifically *LAMP5* and *CCK* (**Figure 2b, d**).

**Figure 2.**
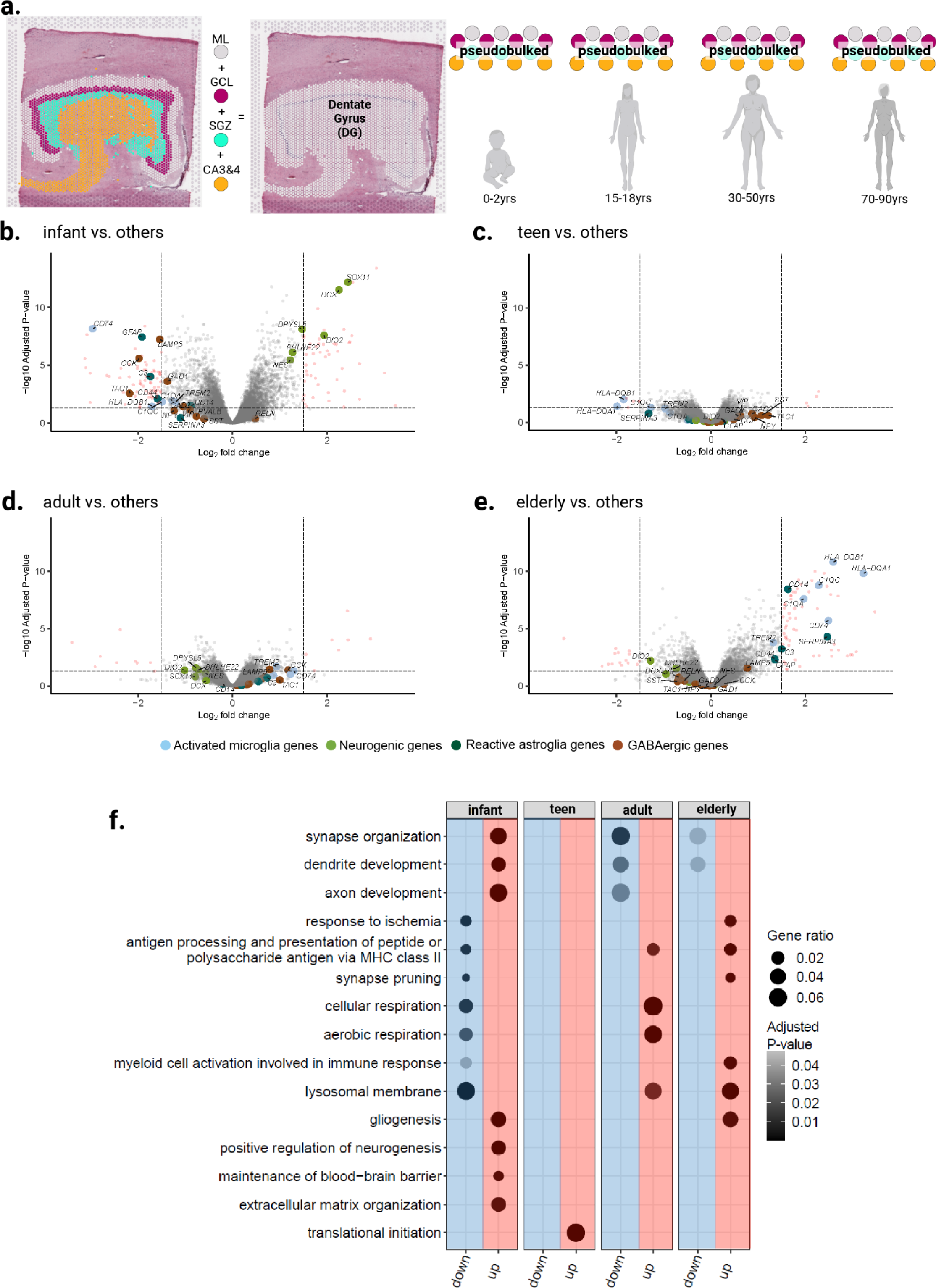
Differential expression (DE) analysis of composite DG identifies genes associated with age group (infant, teen, adult, elderly). (**a**) Pseudo-bulked spatial domains for ML, GCL, SGZ and CA3&4 were collapsed to generate a composite DG molecular profile for each donor. (**b-e**) Volcano plots demonstrate DE comparing one age group to all others for the pseudo-bulked composite DG. Point colors highlight selected genes associated with hallmarks of aging including neurogenesis (green), activated microglia (light blue), reactive astroglia (dark green), and GABAergic genes (brown). The *x*-axis is the log_2_ fold change in expression highlighting genes ≥1.5 logFC or ≤-1.5 logFC and the *y*-axis is the negative log_10_ adjusted *p*-values; red points indicate genes greater than these values. (**f**) Dot plots for major gene ontology terms. Dot plots are faceted by age groups with down- and up-regulation columns colored by blue and red, respectively; dot size represents the fraction of the gene set that was differentially expressed (gene ratio), while black gradient represents adjusted *p*-value.

Gene ontology (GO) analysis further supported increased proliferation of both neurons and glia in the infant group, and enhanced neuroinflammation in adulthood, which persisted in the elderly group (**Figure 2f, Table S4**). With respect to proliferation, we note upregulation of gliogenesis in the elderly group. GO terms for respiration-related processes were increased in the adult, although not the elderly group, suggesting increased oxidative stress. The GO term for blood brain barrier (BBB) maintenance was increased in the infant group, while response to ischemia was decreased in the infant and increased in elderly, suggesting deterioration of vasculature and BBB over age (**Figure 2f**). We also found reversals in GO terms from infancy to advanced age related to development of synapses and dendrites, lysosomal membrane, synapse pruning, and the major histocompatibility class II (MHC-II) immune response (**Figure 2f**). Additionally, the GO term for extracellular matrix (ECM) organization is only enriched in the infant group (**Figure 2f**). To determine spatial heterogeneity of the observed changes in early postnatal development, we next investigated changes across individual DG sub-domains with a focus on infant samples.

### 2.3 Age-associated changes in genes related to inhibitory neurons, neuronal proliferation, and extracellular matrix map to defined areas in the DG

As expected, analysis for pseudobulked DG sub-domains across age groups identified many of the same genes from analysis of the composite DG. However, we obtained improved spatial resolution by mapping DE signals to specific DG sub-domains. Several genes associated with inhibitory interneuron markers were depleted in the infant group across DG sub-domains. While *CCK* was depleted in all DG sub-domains, *GAD1* was depleted specifically in ML and GCL and *LAMP5* in ML, GCL, and SGZ (**Figure 3a-d, Table S5**). To support these findings, we queried existing bulk RNA-seq data from human HPC, which included donors spanning the lifespan ^51^, then compared this bulk HPC RNA-seq data with existing RNA-seq data from human GCL isolated with laser-capture microdissection (LCM) ^51,52^, *CCK* expression increased after infancy in our data as well as the bulk HPC RNA-seq dataset ^51^ (**Figure S11**). Following infancy, *LAMP5*, a gene expressed in a subpopulation of GABAergic inhibitory neurons ^53^, showed unexpected expression in the GCL, which is primarily composed of excitatory neurons (**Figure S12a**). We observed increased *LAMP5* expression over the lifespan in the bulk RNA-seq data ^51^ (**Figure S12b**), and found enrichment of *LAMP5* in the GCL relative to the aggregate HPC in the LCM data ^52^ (**Figure S12c**). We qualitatively validated these changes in *GAD1* and *LAMP5* expression in the GCL using single molecule fluorescent *in situ* hybridization (smFISH) in infant and adult donors (**Figure 3e**) and observed abundant *LAMP5* expression in *SLC17A7*-expressing excitatory neurons in the GCL as well as in *GAD1*-expressing inhibitory neurons in the SGZ, further supporting the existence of *LAMP5* expression in excitatory neurons.

**Figure 3.**
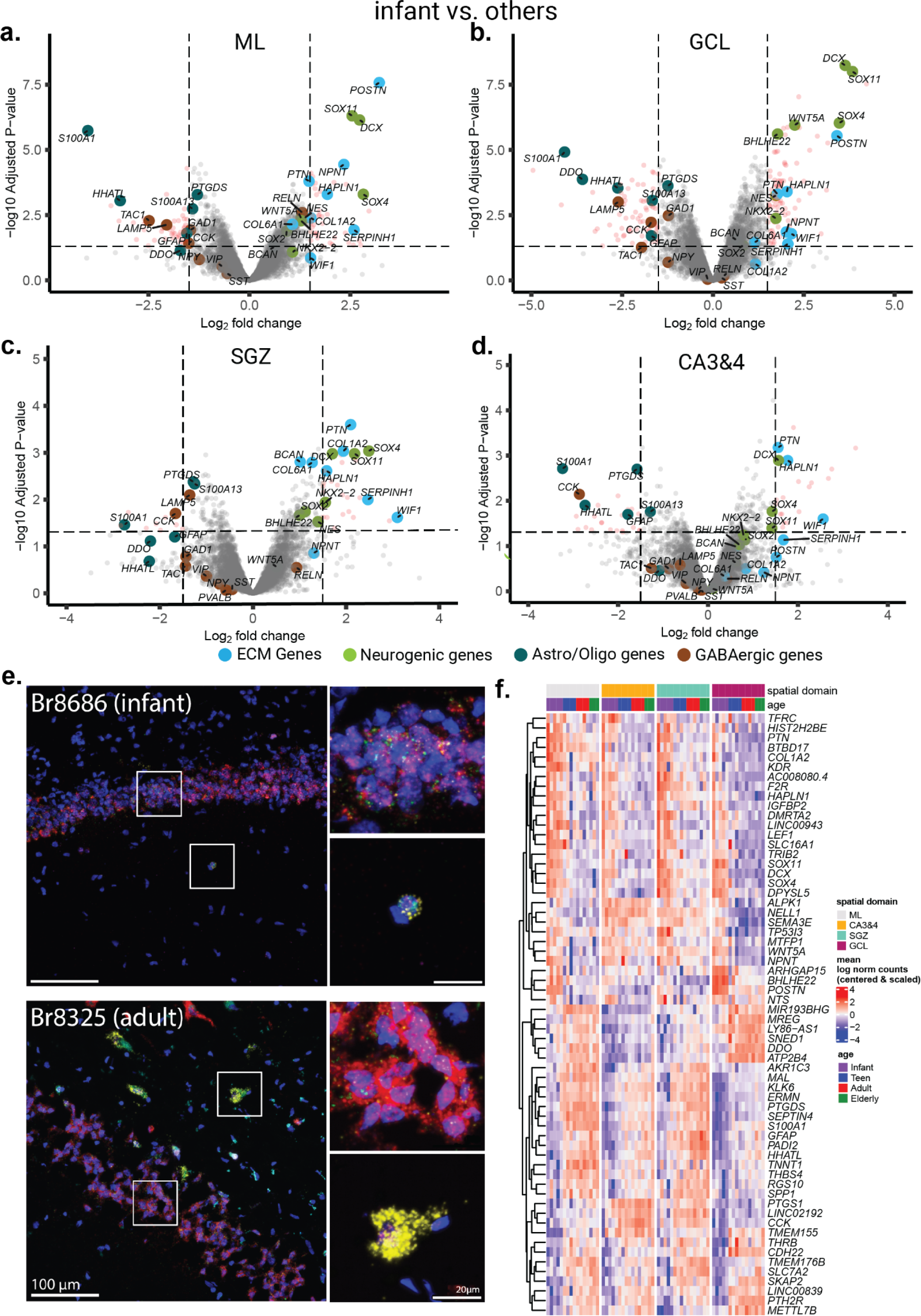
Differential gene expression of infant versus all other age groups across DG sub-domains. (**a-d**) Volcano plots of infant versus non-infant for DG sub-domains: ML (**a**), GCL (**b**), SGZ (**c**) and CA3&4 (**d**). The *x*-axis is the log_2_ fold change in expression and the *y*-axis is the negative log_10_ adjusted *p*-values. (**e**) Representative smFISH images from the infant GCL (top) and adult GCL (bottom) for *PROX1* (green), *GAD1* (yellow), *SLC17A7* (red), *LAMP5* (magenta), and lipofuscin autofluorescence (LIPO, cyan). Scale bar is 100μm, squares denote inset region. Inset scale bar is 20μm. (**f**) Heatmap of the mean log_2_ normalized counts (centered and scaled) for top 5 infant enriched and depleted genes with ≥1.5 logFC or ≤-1.5 logFC from each pseudo-bulked DG sub-domain. Hierarchical clustering was performed across rows. Columns are organized by spatial domains corresponding to DG regions.

Similar to the composite DG analysis, comparison of the infant group versus all other age groups yielded the most DEGs across individual DG sub-domains (**Figure S13a**, **Figure S14, Table S5**). We ruled out that this finding was due to technical variables (**Figure S13b-d**), although we noted that estimated cell counts per spot at the GCL were higher in the infant group, which may contribute to this finding (**Figure S15**). In contrast, the adult and teen age groups had few DEGs in each sub-domain (**Figure S13a, Figure S16, Table S5**). However, because separating the DG into sub-domains decreases the number of spots per age group, this DE analysis may be underpowered to detect age-associated differences.

Given the paucity of DEGs from the other age groups, we asked if we could find gene expression gradients across age for each DG spatial domain by combining the top DEGs unique to infant (infant vs. all others) for all DG sub-domains (**Figure 3f, Figure S17**). This illustrated rapid decreases in expression after infancy of many canonical neurodevelopmental markers, including *SOX4*, *SOX11*, and *IGFBP2*. Expression of *DCX*, a marker of immature granule cells that is frequently used as a proxy of neurogenesis, markedly decreased after infancy (**Figure S18a**), which is in line with findings from several other reports ^31,32,54^. We likewise noted that *DCX* expression rapidly decreased with age in the bulk HPC RNA-seq dataset queried above ^51^ (**Figure S18b**). One of the most significantly depleted genes in infancy was *METTL7B*, which was shown to be upregulated over the lifespan in the excitatory neurons of the human HPC using snRNA-seq ^31^. We found depletion of *METTL7B* expression in the infant ML and SGZ that increased with age (**Figure 3f, Figure S17**). In agreement with upregulation of the GO term for ECM organization in the infant group (**Figure 2f**), we found enrichment of many ECM-related genes in infant ML, SGZ and GCL, including *POSTN*, *PTN*, *WIF1*, *COL1A2*, *BCAN*, *COL6A1*, *HAPLN1*, *SERPINH1*, and *NPNT* (**Figure 3a-d, Table S5**). Expression of *POSTN*, which encodes a cell adhesion molecule (periostin) implicated in rodent hippocampal neurogenesis ^55^, declined with age in the bulk RNA-seq data ^51^ and was enriched in the GCL in the LCM data ^52^ (**Figure 3a-b, Figure S19**).

To further query the extent of neurogenesis across the human lifespan, we obtained previously published molecular profiles generated across multiple species, including mouse, macaques, and human ^20,21,56^, that represented discrete stages of the granule cell maturation trajectory - neural progenitor cells (NPC), neuroblast (NB), immature granule cells (imGC) mature granule cells (GC) - and conducted gene set enrichment analysis in human DG sub-domains across age groups. A gene set for mouse late-stage neuroblasts (NB2) was the earliest stage in this GC maturation trajectory that showed consistent enrichment in the human GCL in all age groups (**Figure S20a**). However, many of the genes included in this set were not restricted to the SGZ, but were also expressed in other DG sub-domains (**Figure S20b, Figure S21**). A gene set for human imGCs, derived with a machine learning algorithm trained on human HPC snRNA-seq data ^57^, showed strongest enrichment in CA3&4, which decreased to non-significant enrichment in the elderly group (**Figure S20a**). In contrast, another gene set for imGCs, which was derived from macaque scRNA-seq data ^21^, showed consistent enrichment in the GCL for all age groups (**Figure S20a,c**). Together, these data show that although genes associated with neurogenesis in other species are expressed in the human HPC throughout the lifespan, they are not spatially restricted to the neurogenic niche.

### 2.4 Nmf-based transfer learning suggests continued presence of immature granule cells in adulthood

To extend our gene set enrichment analyses, we next asked if signatures of neurogenesis could be queried with other computational strategies. Non-negative matrix factorization (nmf) is a dimensionality reduction technique where the resulting latent factors could be used to identify patterns of coordinated gene expression that correlate with biological processes, including cell type and cell state ^58^. Nmf patterns generated from one dataset can predict the presence and distribution of these same latent factors in another dataset via transfer learning ^58^. This strategy was recently used to integrate snRNA-seq and Visium data from adult neurotypical donors in HPC ^59^. Specifically, 100 nmf patterns were identified in snRNA-seq data from which we identified three patterns (nmf26, nmf5, and nmf14) that were enriched in GC populations with a sequential overlap of weight gradients where nmf26 partially overlaps with nmf5, which partially overlaps with nmf14 (**Figure 4a**). Projecting these three patterns onto paired SRT data from the same donors ^59^ revealed that, although all three patterns were restricted to the GCL (**Figure 4b**), nmf26 weights were more sparsely distributed. We thus compared the top 10 marker genes for these three patterns to better understand the biological relevance (**Figure 4c**). Interestingly, top gene markers for nmf26 (**Figure 4c**) overlapped with genes enriched in the infant GCL (**Figure 3b, Table S5**) including *POSTN* (**Figure S19**), *TARID*, *FIGN*, *WIPF3*, *NREP*, and *PTGFR*. The top gene marker for nmf26, *ALDH1A2*, encodes a retinaldehyde dehydrogenase enzyme that synthesizes retinoic acid and regulates granule cell differentiation in rodents ^60^.

**Figure 4:**
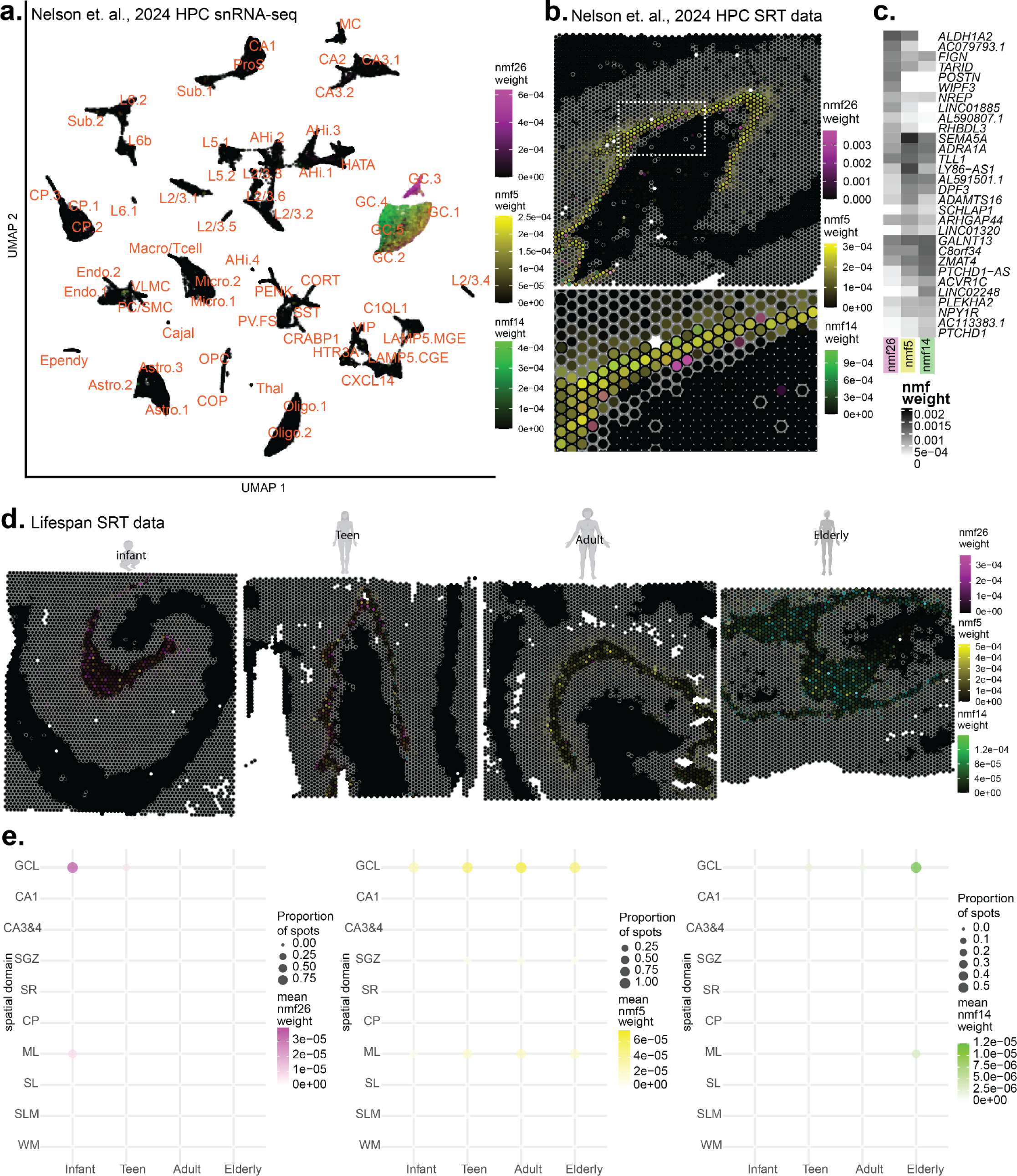
Nmf patterns approximate a maturation trajectory for human dentate granule cells. (**a**) UMAP of human adult HPC snRNA-seq dataset ^59^ with superfine cell class annotations and nuclei are colored by the weight of three nmf patterns associated selectively with the GC superfine cell class. RGB scale converted to represent all three nmf patterns simultaneously. Overlapping gradients of weights from different nmf patterns have a blended color for a given nucleus. **(b)** Representative spot plots from the adult HPC SRT dataset ^59^. Spot outlines are colored black to delineate GCL, CA3&4, and CA1, other spatial domains are outlined in grey. Spots are colored by weight of the three nmf patterns. Inset (below) is zoomed into the GCL. (**c**) Heatmap of gene-level nmf weights of top 10 marker genes for nmf26, nmf5, and nmf14 ^59^. (**d**) Representative spot plots for a given age group from the human DG lifespan SRT dataset. GCL, CA3&4, CA1 spots outlined in black, others outlined in grey. Spots are colored by weight of nmf26, nmf5, nmf14. (**e**) Dot plot of spatial domain versus age group across all donors. Size represents the proportion of spots with non-zero nmf pattern weights, and respective color scale represents the mean values of weights for a given nmf pattern.

Given the above findings, we asked if these nmf patterns were capturing a gene expression profile suggestive of a granule cell maturation trajectory. Using transfer learning, we next projected these three snRNA-seq nmf patterns identified in the adult HPC into the lifespan Visium data collected here. As expected, the three patterns were restricted to the GCL, but each pattern was strongly associated with age (**Figure 4d,e, Figure S22**). Nmf26 was expressed in many spots throughout the infant GCL with high weights, but was present in fewer spots and at decreased weights in the teen, adult and elderly groups. Nmf5 and nmf14 showed increased prevalence and higher weights at the GCL with age, although the nmf5 weights peaked in adulthood while nmf14 was primarily restricted to the elderly group. Projecting these patterns onto a scRNA-seq data from the adult macaque DG ^21^ revealed that nmf26 weights were enriched in a subset of NB and imGCs, while nmf5 and nmf14 weights were enriched in mature GCs (**Figure S23a,c**). Likewise, in a neurogenic trajectory dataset from the mouse ^20^, nmf26 weights were enriched in juvenile and immature GCs, while nmf14 was largely absent (**Figure S23b,d**). Together, this data supports existence of a spatially-organized maturation trajectory where a subpopulation of immature GCs that are abundant in the infant, are captured by nmf26, but rapidly decline over aging. In contrast, nmf5 captures the majority of GCs, while nmf14 captures a more mature expression profile not found in infancy.

As depicted in the UMAP, nmf26 (infant-associated) and nmf14 (non-infant-associated) are enriched in different GC clusters (GC.3 and GC.4, respectively) (**Figure 4a**). To identify genes that distinguish between these two cell types, we used the pseudobulked DE results taken from Nelson et al., 2024 ^59^ which calculated DEGs by comparing one cluster to all others. We then compared GC.3 and GC.4 to identify gene uniquely enriched in either GC.3 or GC.4 (**Figure S24a**). Similar to our findings in the infant GCL (**Figure 3b, Figure S19a**), *POSTN* was enriched in GC.3 (**Figure 4c**). Another gene which showed differential enrichment in GC.3, and depletion in GC.4, was *FST*, which, when knocked out in rodent models, negatively affects adult neurogenesis, learning, and synaptic plasticity ^61^. GO analysis of these DEGs highlighted opposing, functional differences between GC.3 and GC.4 related to development, cellular signaling, RNA translation and ECM (**Figure S24b**). For example, neurogenesis and neuron differentiation GO terms were simultaneously down- and up-regulated in GC.3, while the GO term for generation of neurons was down-regulated, suggesting that GC.3 neurons are undergoing granule cell maturation as opposed to proliferation (**Figure S24b**). GC.3 also exhibited functional enrichment of GO terms related to synaptic plasticity and transmission, suggesting that GC.3 cells may be more primed for plasticity compared to GC.4 (**Figure S24b**). Many terms for developmental and differentiation were down-regulated in GC.4, suggesting that neurons in the GC.4 cluster are more mature than GC.3 (**Figure S24b**). Importantly, while these analyses suggest a data-driven signature of imGCs present in adult HPC, we do not find a signature indicative of neural proliferation. However, GO analysis and DE results in infants suggest continued proliferation over aging of other cell types (**Figure 2)**, namely glial cells, paired with increases in glial activation states.

### 2.5 Neuroinflammatory cell abundance and activity is a robust signature of aging in the DG

While early postnatal development is marked by proliferation and differentiation of neurons, these biological processes shift from neurons to glia following infancy. We noted a decrease in gene signatures associated with neurogenesis and a concurrent increase in gliogenesis-related gene expression over aging (**Figure 2**, **Figure 3**, **Figure 4**). We observed significant depletion of pan-glial gene expression in specific DG subfields in infancy, including genes typically expressed in oligodendrocytes and astrocytes such as *GFAP* in ML, GCL, and CA3&4; *TMEM176* in ML, SGZ, and GCL; *S100A1, S100A13,* and *PTGDS* in all DG sub-domains; and *HHATL* in ML, GCL, and CA3&4 (**Figure 3a-d, Table S5**). We also observed enrichment of GO terms associated with MHC-II peptides in the elderly group in the composite DG (**Figure 2e**-**f**). Since MHC-II peptides are expressed in late-stage activated microglia ^41^, we investigated spatial enrichment of markers for activated microglia ^41,62^. These genes were enriched in WM and neuropil-rich domains including the ML, SL, SLM, and SR, spatial domains that are characterized by abundant neuronal processes and a lack of cell bodies (**Figure 5a**). Many genes involved in the MHC-II immune response also showed age-dependent increases in specific DG sub-domains (**Figure 5b**), including enrichment in the elderly ML (**Figure 5c**). While not significantly enriched in any individual sub-domain, *CD74,* a reactive microglia marker, was enriched in the composite DG of the elderly group (**Figure 2d**), and its expression increased with age in all DG sub-domains (**Figure 5b, Figure S25a**). We confirmed this finding by querying bulk HPC RNA-seq data ^51^ for *CD74* expression with age (**Figure S25b**). Likewise, the microglial marker *TREM2* was enriched in the composite DG of the adult and elderly groups (**Figure 2d-e**), although not in any individual DG sub-domain. Microglial marker genes were generally upregulated, including *C1QB* and *IL4R ^45^* in the ML (**Figure 5c**) and in CA3&4 (**Figure 5f**) of the elderly group. While not significantly enriched in any individual DG sub-domain, *HAMP*, which encodes hepcidin and is associated with iron sequestering in microglia and neurons, was enriched in composite elderly DG (**Table S3**).

**Figure 5.**
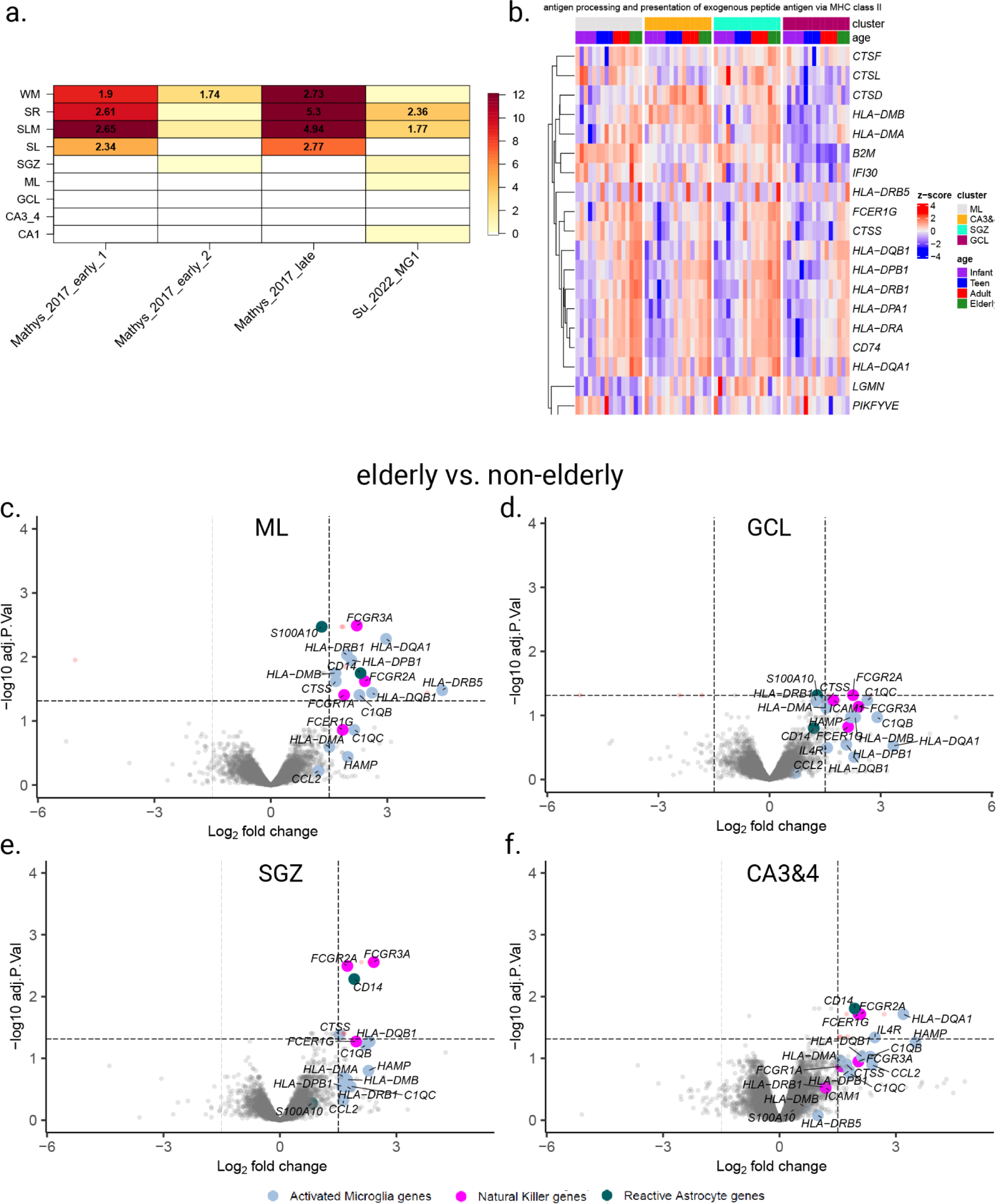
Markers of inflammation are enriched in DG sub-domains in the elderly group. (**a**) Enrichment analyses using Fisher’s exact tests for predefined gene sets for activated microglial markers. Nomenclature adopted from each publication: microglial clusters for early_activated_1, early_activated_2, late_activated ^41^, and human microglial cluster 1 (MG1) ^62^. Color indicates negative log_10_ *p*-values while numbers within significant heatmap cells indicate odds ratios for the enrichments. (**b**) Heatmap of the mean log_2_ normalized counts (centered and scaled) of pseudo-bulked data limited to DG sub-domains with genes, which satisfy gene ontology accession GO:0002504. Hierarchical clustering was performed across all rows. Columns are organized by spatial domains corresponding to DG regions and by age group. (**c-f**) Volcano plots of elderly vs. non-elderly for individual DG sub-domains: ML (**c**), GCL (**d**), SGZ (**e**) and CA3&4 (**f**). The *x*-axis is the log_2_ fold change in expression and the *y*-axis is the negative log_10_ adjusted *p*-values.

Interestingly, we observed a variety of immune-related gene expression in elderly samples, including genes expressed in activated human microglia and other immune-related cell types. For example, *FCGR3A*, which is enriched in elderly ML, SGZ, and CA3&4 (**Figure 5c, e-f**), regulates survival and proliferation of natural killer (NK) cells ^63,64^. To investigate changes in estimated cell proportions, we applied `cell2location` ^65^, a method for spot-level deconvolution of cell types ^66^, to publicly-available snRNA-seq data from human HPC, which we subset to include only the DG ^31^ (**Methods**, **Figure S26**, **Figure S27**, **Figure S28**). We found an increasing proportion of microglia and T-cells in the SGZ with age (**Figure S29a, Figure S29c**).

Activated microglia can induce reactive astrocytes ^39^, and we detected enrichment of genes that are markers of reactive astrocytes in the composite elderly DG, including *GFAP*, *C3*, *SERPINA3*, *EMP1*, *CD109*, *CD44*, *SERPING1*, and *FKBP5* (**Figure 2e, Table S3**). Additional genes associated with reactive astrocytes show spatial specificity, including *S100A10* in the ML and *CD14* in the elderly ML, SGZ, and CA4&3 (**Figure 5c-f, Table S5**). We also investigated changes in estimated cell proportions for oligodendrocytes and subtypes of astrocytes across aging. `Cell2location` estimated an increase in proportion for oligodendrocytes in ML, SGZ, and CA3&4 with age (**Figure S30b**), and for both subtypes of astrocytes (Astro_1, *GFAP*^+^ and Astro_2,*GFAP*^-^) in the ML and SGZ with age (**Figure S30c-d**). Taken together, these results suggest enrichment of activated microglia and other neuroinflammatory markers in specific HPC domains, and spatially restricted vulnerability to age-related gene expression changes within sub-domains of the DG (**Figure 5a**).

### 2.6 A minimal gene set tracks wide-spread hippocampal tissue changes with aging

Given strong association of many specific gene markers with age-related processes, including proliferation and inflammation (**Figure 2**, **Figure 4**, **Figure 5**), we asked if a minimal set of genes could track the spatial domains where gene expression is most dynamic over aging. A previous study compared bulk and single nucleus transcriptomic profiles from microdissected areas of various brain regions from mice at various ages, paired with SRT profiles, and identified that WM-rich spatial domains were most vulnerable to gene expression changes over aging ^67^. We adapted this data-driven workflow ^67^, and used DEGs derived from comparing older age groups to the infant age group within each spatial domain to compute a common aging score (CAS). Each CAS is a single value that represents the difference in transcriptional programs between biological conditions; in our case, infant versus each other age.

To obtain the set of DEGs for computing the CAS score, we aimed to leverage all spots within the tissue, and thus included all spatial domains in our samples. To control for possible differences in cell types across spatial domains, we performed DE within each of the 9 pseudo-bulked spatial domains (DG: ML, GCL, SGZ, CA3&4, nonDG: CA1, SLM, SL, SR, WM, excluding CP) for each non-infant age group compared to the infant group. Genes were selected based on thresholds of adjusted *p*-values (smaller than 0.05 and a log_2_ fold change of at least 1.5 in at least two of the differential expression tests), and were shared with at least 3 spatial domains. This produced a minimal, signed (+/-), gene set of 46 (+) and 13 (-) genes associated with human aging relative to the infant group (**Table S6**). Some of these DEGs included the neurogenesis-related markers *DCX*, *SOX11*, and *SOX4,* which decrease with age. Other DEGs increased with age, including oligodendrogenesis markers *MAL* and *OPALIN*, and markers associated with MHC class II and inflammatory signaling (**Figure 6a, Figure S31**). GO analysis of these aging DEGs suggested downregulation of neural proliferation paired with upregulation of inflammatory activity and myelination (**Figure 6b**).

**Figure 6.**
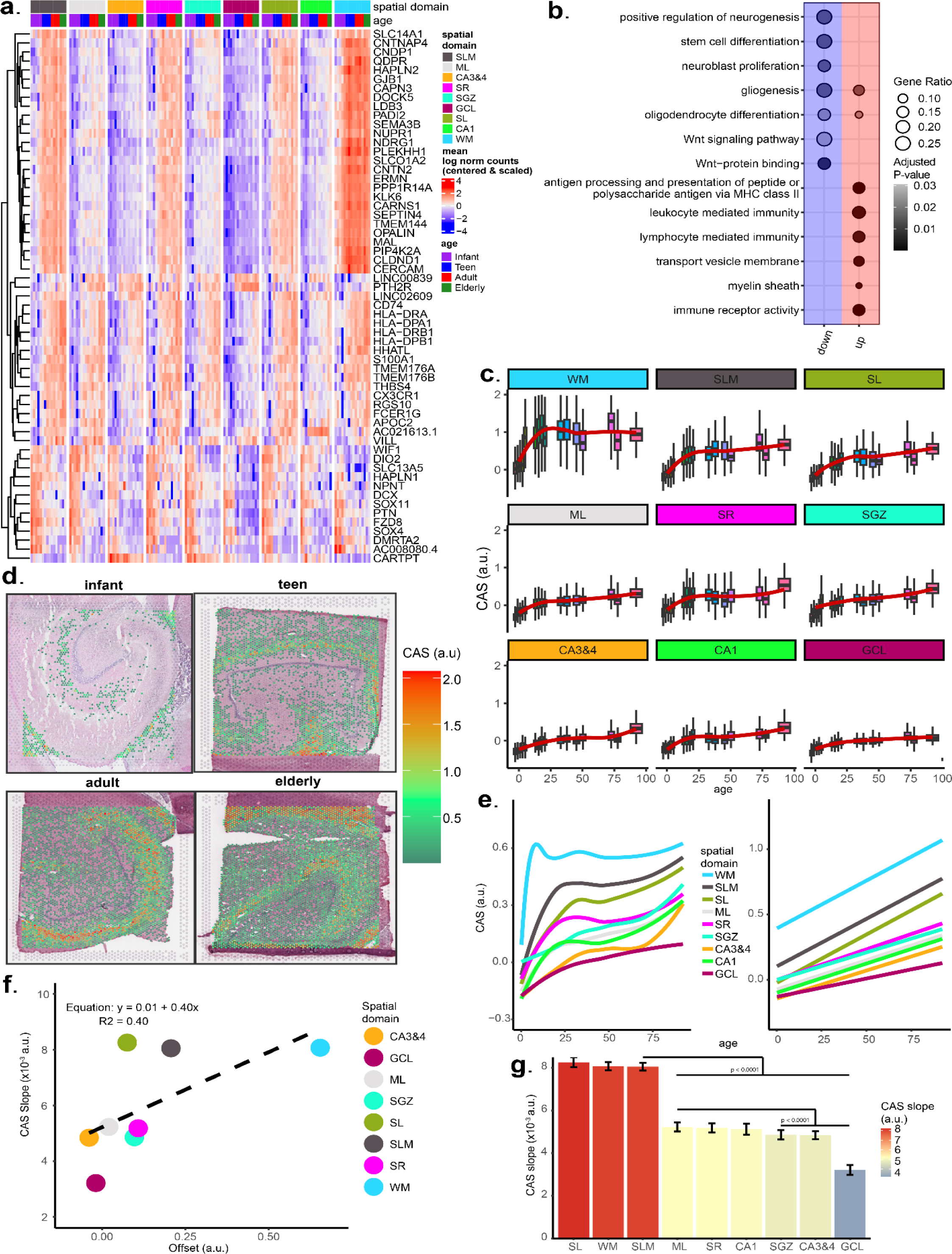
Wide-spread HPC aging signature identifies regions of local tissue that change more with age. (**a**) Heatmap of the mean log_2_ normalized counts (centered and scaled) for the wide-spread aging signature gene set. Hierarchical clustering was performed across rows. Columns are organized by spatial domains corresponding to HPC spatial domains. Labeled genes are those that show strong increasing age gradients across many spatial domains. (**b**) Dot plot for major gene ontology terms from aging signature gene set. Dot plots are faceted by their down- and up-regulation columns colored by blue and red, respectively; dot size represents the fraction of gene set that was differentially expressed (Gene ratio), while black gradient represents adjusted p-value. (**c**) Boxplots of CAS (in arbitrary units) of individual Visium spots versus age, faceted by HPC spatial domain and fitted with local regression line. (**d**) Data visualization of the Visium spots for four representative samples from each age group (infant (Br8533), teen (Br1412), adult (Br3942), elderly (Br5242)). Color of spots are for CAS in arbitrary units. (**e**) CAS trajectories of all HPC spatial domains vs. age approximated via local regression (left) and linear regression (right). (**f**) Offset of linear fit and slope comparison from linear modeling of CAS across all Visium spots. (**g**) CAS slope of linear approximations in **e**, colored by slope, for each spatial domain. Mean ± 95% confidence intervals. Adjusted *p*-values derived from two-sided Tukey’s HSD test.

Using this widespread aging gene set as input, we employed the `Vision` package ^68^ to compute CAS for each gene expression spot in all capture areas in our study (**Figure 6c-d**, **Figure S32**). We then ordered CAS values for each spot by age for each spatial domain. The trajectory of these values rapidly increased from infancy to teen for all spatial domains, but the increase and variability were larger in glial- and neuropil-enriched spots (**Figure 6c**), reflecting the changes in expression of the genes that comprise the human DG aging gene signature derived from these data (**Figure 6a**). We observe the largest changes in CAS in WM spots, which also have more CAS heterogeneity (**Figure 6c-d**). This implies localized changes over aging within WM. It is important to note that our human DG DEGs and CAS results did not take into account changes in cell density with age, which have decreasing trends for the GCL and increasing trends for WM (**Figure S15**). In the mouse CAS study ^67^, the youngest age group was 3 months old, which approximately correlates to humans in their mid-twenties ^69,70^. As a result, although we do see similarities in spatial assignment of mouse-derived CAS values (**Figure S34a**), our human-derived values were more sensitive to spots enriched for oligodendrocytes (**Figure 6c-d**, **Figure S32, Figure S33, Figure S34b**). Comparing our human DG CAS values to those derived from the whole mouse brain ^67^ highlights the importance of including younger ages in developmental comparisons of age-related gene expression changes.

To assess if CAS was solely driven by cell type changes over development and aging, we investigated the relationship between CAS and the predicted cell type proportions in each spot. Using the outputs from `cell2location` (**Figure S27**, **Figure S28**), we applied PCA to extract the top components of variation for cell type proportions for all spots^71^. Differences in the predicted cell type proportions were not sufficient to explain the majority of variation in CAS values (**Figure S33**), although PCs 1-4 did explain 38% of the variation between the spot-level CAS values and predicted cell type proportions per spot. We note that the relationship was particularly strong for oligodendrocytes, but we do not detect major differences in the proportion of spots from WM with age (**Figure S10**), and thus this relationship may be driven by increased oligodendrogenesis post-infancy ^72^ rather than differences in the amount of WM present in each sample.

CAS trajectories can be approximated with linear modeling ^67^, an approach that allowed us to use the slope, or rate of change across age, to infer a “CAS velocity” for each spatial domain (**Figure 6e**). We then used the slopes to approximate the spatial domains that exhibit a higher rate of aging-associated changes as captured by our DEGs. Similar to SRT data from mice ^67^, the CAS baseline (i.e. CAS assigned to the infant spots) did not strongly predict a spatial domain’s CAS velocity across age (**Figure 6f**). Comparing CAS velocities for each spatial domain, we found that glial-enriched spatial domains had the largest CAS velocities, followed by neuropil-enriched spatial domains (**Figure 6g**). This is in agreement with the mouse study ^67^ in which age-associated changes were enriched in glial-rich regions. Of note, the ML, which is the most neuropil-enriched domain in the DG, had a significantly larger CAS velocity than any other DG domain. This highlights the importance of retaining cytoplasmic transcripts, which are captured in SRT approaches, but not snRNA-seq (**Figure 6g**). The differences in CAS velocities between spatial domains suggests that the human HPC exhibits differential aging within its sub-domains, and that glial-enriched and neuropil-enriched domains exhibit a greater degree of aging-related changes in gene expression. This is consistent with our results suggesting increased microglia abundance and microglia-related gene expression signatures over aging.

## 3 Discussion

Transcriptomic profiling studies in the human HPC remain relatively limited. Previous studies have used bulk or laser-capture microdissection RNA-seq ^51,52,73^, which has limited spatial accuracy and cell type resolution. More recent studies have begun to use snRNA-seq ^31,32,56,62,74^, which performs at cellular resolution, but does not retain spatial information, and cannot capture transcripts from extra-nuclear compartments. SRT can provide transcriptome-wide resolution with spatial fidelity and retain cytosolic transcripts, but it was only recently deployed in human HPC, and in a limited number of donors ^54^. Here, we used the 10x Genomics Visium Spatial Gene Expression platform to identify molecular signatures across human DG sub-domains that are specifically associated with individual lifespan stages (infant, teen, adult and elderly).

In the infant DG, we observed an increased abundance of genes encoding extracellular matrix (ECM)-related molecules that have also been implicated in HPC function during development. *PTN* (pleiotrophin) and its receptor *PTPRZ1* are associated with increased DG neurogenesis in a mouse model of senescence following exposure to environmental enrichment ^75,76^. *HALPN1* (hyaluronan and proteoglycan link protein 1) is important for maturation of perineuronal nets in the mouse dorsal CA1 that surround PV+ interneurons, which affects neuronal allocation and memory precision ^77^. Periostin (*POSTN*) promotes neural stem cell proliferation as well as neuronal and astroglial *in vitro*, while periostin injection into the lateral ventricles of neonatal rats after hypoxic-ischemic brain injury stimulates proliferation and differentiation in the SGZ ^55^. Interestingly, *Postn* is not expressed after birth in the mouse DG ^78^, suggesting a human-specific role postnatally. Our results provide important insight about age-associated changes in ECM composition, density, and localization across the DG spatial topography. Identification of ECM-related genes that are enriched in the human DG during infancy, may provide important insight for therapeutic strategies to improve cellular rejuvenation and proliferation.

Our data does not support continued neural proliferation at the GCL following infancy. However, our spatial enrichment analysis (**Figure S20**) and results from our nmf-based transfer learning analyses (**Figure 4**, **Figure S22**, **Figure S23, Figure S24**)^59^ support the existence of a subpopulation of GCs expressing immature markers through adulthood ^79^. Our data cannot determine whether this population persists due to protracted maturation, if the expression of immature markers is indicative of a reversion to a more immature cellular state, or if this immature cell state represents an alternative endpoint in granule cell maturation. In support of prolonged maturation in the human DG, maturation trajectories of GCs in macaques are up to six times longer than those in rodents ^80^. Further supporting the ability of GCs to revert in the maturation state, chronic fluoxetine administration in mice induces reversion of GCs to immature states ^81^. While we cannot definitively determine the origin of cells that are enriched for imGC pattern nmf26, many genes that define this pattern are enriched in the infant GCL. Some of these nmf26 marker genes, including *POSTN* and *ALDH1A2*, exhibit species-specific expression patterns. *ALDH1A2* and other genes that encode enzymes that synthesize retinoic acid (RA) ^82^ are enriched in the human GCL, but are not highly expressed in the adult mouse DG ^60^. RA promotes synaptic plasticity in DG neurons ^83^, and it also affects DG neurogenesis in a biphasic manner - depletion reduces neural differentiation and cell survival ^84,85^ while excess amounts reduce neural proliferation ^86^. Given these established roles for *ALDH1A2* and *POSTN* in GC plasticity and development, our data showing specificity of their expression in the human GCL renders these genes as prime candidates for follow up studies. Manipulating expression of these genes in mice postnatally could help determine whether modulating ECM and RA signaling can stimulate proliferation or modulate imGC fate in the DG.

We find increased expression of markers for GABAergic neurons, including *GAD1* and *LAMP5*, in the GCL, a domain primarily composed of excitatory neurons. Although initially surprising, *DCX* and *GAD1* co-expression was previously demonstrated in the adult GCL using smFISH and immunohistochemistry ^31,54^. However, *LAMP5*^+^ GABAergic neurons in the GCL have not been reported. Although LAMP5 protein is localized in axon terminals of GABAergic neurons ^87^, our smFISH data shows *LAMP5* transcripts localized in cell bodies. Interestingly, *LAMP5*^+^ excitatory neurons are found in human samples in cortical layer 2/3 and in the basolateral amygdala, and both inhibitory and excitatory *LAMP5*^+^ cells are more abundant in humans than mice ^88,89^. Additional research is needed to determine if the increase in *LAMP5* over age is due to increased *LAMP5* transcription in expressor cells, recruitment of previously non-expressing cells to the *LAMP5*+ population, or proliferation of a *LAMP5^+^*population.

We observed the most DEGs in the infant group, followed by the elderly group, suggesting that gene expression changes undergo major shifts over aging. Many genes that were enriched in the elderly group were associated with neuroinflammation. Because our data leverages the inclusion of cytoplasmic, non-nuclear transcripts, we are able to provide evidence that neuropil-rich regions are preferentially vulnerable to increases in inflammation-related gene expression (**Figure 5**, **Figure 6**). Since microglia show a robust cellular signature of aging in the DG, there is strong interest in evaluating these cells as therapeutic targets for age-associated neurodegeneration and cognitive decline. Our data contributes to existing evidence that microglia accumulate throughout the DG with aging. However, we cannot distinguish if inflammation-associated gene expression is limited to microglia, or if other immune cells, such as NK cells, or BBB disruption contribute to these signatures. We observe evidence of disruption to the BBB (**Figure 2d**, **Figure 5**) and of increased oxidative stress (**Figure 2e**), which could drive inflammation and contribute to decreases in neuronal proliferation ^42,90–92^. However, there is conflicting evidence regarding the presence and origin of invasive immune cells in the HPC^35,93–95^, and although NK cells, T-cells and microglia share some overlapping transcriptional signatures, the extent of overlap in humans is unclear. However, microglial markers also share some overlap with neurons. For example, *CD74* is expressed in activated microglia, but expression of CD74 protein has also been detected in neurons ^96–98^ and it may be involved in AD pathology ^96^. These findings highlight the importance of analyzing both neuronal and non-neuronal cellular populations across species in aging.

Our study has several limitations, including the relatively small sample size (5 infant donors, 4 teen donors, 4 adult donors, and 3 elderly donors, and no childhood donors), which likely limit statistical power to identify age-associated DEGs across groups. Given their relatively large diameter (55 μm diameter), Visium spots can contain multiple cell types, and cell density across spots can be heterogeneous. Thus, age-related statistics, including CAS, may be biased by changes in cell composition or abundance, and hence do not represent a truly cell-independent signature of aging. Our data lacked nuclei segmentation for some samples, precluding inclusion of nuclei density as a covariate in CAS calculations. Despite these limitations, transcriptome-wide profiles of the human HPC with spatial resolution are lacking. Our study, with the inclusion of donors across the lifespan and inclusion of extranuclear transcripts with spatial domain-level specificity, is an important resource which provides critical insight into species-specific cellular mechanisms of HPC development and age-related changes in hippocampal function.

## 4 Methods

### 4.1 Tissue samples

Postmortem human brain tissue from male and female neurotypical donors (*N*=17) of European and/or African ancestry spanning ages 0.18 to 76.7 years were obtained by brain donations collected through the following locations and protocols at the time of autopsy: the Office of the Chief Medical Examiner of the State of Maryland, under the Maryland Department of Health’s IRB protocol #12–24, the Departments of Pathology at Western Michigan University Homer Stryker MD School of Medicine and the University of North Dakota School of Medicine and Health Sciences, and the County of Santa Clara Medical Examiner-Coroner Office, all under WCG IRB protocol #20111080. One additional sample was consented through the National Institute of Mental Health Intramural Research Program (NIH protocol #90-M-0142), and was acquired by LIBD via material transfer agreement. All donors were obtained with informed consent from the legal next of kin. All donors here were negative for illicit drugs of abuse at time of death. Demographics for the 17 donors are listed in **Table S1**. Clinical characterization, diagnoses, macro- and microscopic neuropathological examinations, were performed on all samples using a standardized paradigm; subjects with evidence of macro- or microscopic neuropathology were excluded. Details of tissue acquisition, handling, processing, dissection, clinical characterizations, diagnoses, neuropathological examinations, RNA extraction and quality control (QC) measures have been described previously ^99^. Briefly, the anterior half of the HPC containing dentate gyrus was microdissected using a hand-held dental drill as previously described ^51^, staying within the anterior half of the HPC as guided by visual inspection of the HPC itself, anterior to the appearance of the lateral geniculate nucleus and progressive diminution of the putamen. Tissue blocks were then stored at -80°C.

### 4.2 Tissue processing and quality control

Frozen brain blocks were embedded on the posterior end in OCT (TissueTek Sakura) and cryosectioned at −10°C (Thermo Cryostar). Brain blocks were cryosectioned and stained with hematoxylin and eosin (H&E) to verify the presence of anatomical landmarks of the dentate gyrus such as the GCL. Sections were placed on chilled Visium Tissue Optimization Slides (catalog no. 3000394, 10x Genomics) and Visium Spatial Gene Expression Slides (catalog no. 2000233, 10x Genomics), and adhesion of tissue to slide was facilitated by warming the back of the slide. Tissue sections were then fixed in chilled methanol, and stained according to the Methanol Fixation, H&E Staining & Imaging for Visium Spatial Protocol (catalog no. CG000160 Rev C, 10x Genomics) or Visium Spatial Tissue Optimization User Guide (catalog no. CG000238 Rev C, 10x Genomics). Visium Tissue Optimization Slides were used to choose the optimal permeabilization time. For gene expression experiments, tissue was permeabilized for 18 min, which was selected as the optimal time based on tissue optimization time-course experiments. Brightfield histology images of H&E stained sections were taken on a Leica Aperio CS2 slide scanner equipped with a color camera and a 20x/0.75 NA objective with a 2x optical magnification changer for 40x scanning, or on an Olympus VS200 slide scanner equipped with a 10x/0.4NA objective. For tissue optimization experiments, fluorescent images were taken with a Cytation C10 Confocal Imaging Reader (Agilent) equipped with TRITC filter (ex 556/em 600) and 10x objective at approximately 400ms exposure time. Sample Br3874 contained no GCL as verified by H&E staining and incorrect spatial domain assignment by `BayesSpace` and was removed from downstream analyses (**Figure S3**).

### 4.3 Visium data generation

Libraries were prepared according to the Visium Spatial Gene Expression User Guide (CG000239 Rev C, 10x Genomics). For two slides, Visium Spatial Gene Expression Slides were shipped to 10x Genomics after tissue mounting for H&E staining, and imaging, cDNA synthesis, and library preparation. Libraries were quality controlled with Agilent Bioanalyzer High Sensitivity dsDNA chips and sequenced on a NovaSeq 6000 System (Illumina) at a sequencing depth of a minimum of 60,000 reads per Visium spot. Sequencing was performed using the following read protocol: read 1: 28 cycles; i7 index read: 10 cycles; i5 index read: 10 cycles; and read 2: 90 cycles.

### 4.4 Visium raw data processing

The manual alignment of raw histology images were processed by sample using 10x Genomics `Loupe` browser [v.6.0.0]. Raw sequencing data files (FASTQ files) for the sequenced libraries were processed using 10x Genomics `SpaceRanger` software [v. 1.3.1], which uses human genome reference transcriptome version GRCh38 2020-A (July 7, 2020) provided by 10x Genomics for genome alignment ^100^. The preprocessed Visium data for each sample, integrated with the output from `VistoSeg` ^101^ (see Methods Section 4.5, Visium-H&E image segmentation and processing), were stored in a R/Bioconductor S4 class using the `SpatialExperiment` v.1.6.1 R/Bioconductor package ^102^.

### 4.5 Visium-H&E image segmentation and processing

Nuclei segmentation was performed using `VistoSeg`, a MATLAB-based software package ^101^ for samples for which sufficiently high resolution images were acquired (**Figure S15**). Briefly, Gaussian smoothing and contrast adjustment were performed to enhance the nuclei in the image. The image is converted L*a*b color space. The a*b color space is extracted from the L*a*b-converted image and is given to a K-means clustering function, along with the number of colors (*k*) the user visually identifies in the image; here *k*=5. The function outputs a binary mask for each of the (*k*) distinguishable color gradients in the image. The segmentation is further refined by extracting the intensity of the pixels from the binary mask of nuclei and applying an intensity threshold to separate the darker nuclei regions at center from the lighter regions at the borders. `Loupe` Browser v.6.0.0 produces a JSON file for each full-resolution capture area tiff from `VistoSeg`, encoding properties of the image e,g., spot diameter in pixels. `SpaceRanger` provides a .csv file for each full-resolution capture area tiff that includes information for each spot with an identification barcode and pixel coordinates for the center of the spots. The `VistoSeg` package integrates these files with the segmented imaging data to provide a final table with nuclei count per Visium spot for each capture area.

### 4.6 Spot-level data processing

All Visium data analyses were performed using a `SpatialExperiment` (*spe*) S4 class storing the object constructed with the `SpatialExperiment` v.1.6.1 R/Bioconductor package ^102^. The *spe* class extends the `SingleCellExperiment` class used for scRNA-seq data for spatial context, with observations at the level of Visium spots rather than cells. Objects in the *spe* class hold additional spatial information e.g., `colData` has information on whether Visium spots overlap with the tissue from imaging data, `spatialCoords` has the *x*- and *y*-coordinates of the Visium capture areas, and `imgData` holds the imaging files and information pertaining to the images (such as pixel resolution). To the *spe* class, we added information including the sum of segmented cells per spot computed from the `VistoSeg` package, the sum of UMIs per spot, the sum of genes expressed per spot, donor age, donor sex, RNA integrity numbers (RIN), race, and post-mortem interval (PMI) in hours.

Spot level analysis was performed as previously described ^48,103,104^. Briefly, spot level quality control (QC) were evaluated using the `perCellQCMetrics()` function from the `scuttle` package v.1.6.2 R Bioconductor package ^49^ and low quality spots having low UMI counts, low gene counts, or high percent of mitochondrial genetic expression were dropped using the `isOutlier()` function of the same package (**Figure S1**, **Figure S2**). The `scran` v.1.24.0 R Bioconductor package ^105^ functions `quickCluster()` (blocking for each brain donor) and `computeSumFactors()`, then `logNormCounts()` function from the `scuttle` package were used to compute the log-transformed and normalized gene expression counts at the spot level. The `scran` package function `modelGeneVar()` was used to model the gene mean expression and variance (blocking for each brain donor), and `getTopHVGs()` was used to identify the top 10% HVGs. The top 10% HVGs were used to compute 50 principal components (PCs) with the `runPCA()` function from `scater` v.1.24.0 R Bioconductor package and `runUMAP()`, from the same package, was used for Uniform Manifold Approximation and Projection (UMAP) dimensionality reduction ^49^. Primarily for the purposes of unsupervised spatial clustering, corrections for potential batch effects and high dataset variabilities were performed by employing an transcriptomic data integration algorithm that projects Visium spots into a shared dimensionally reduced PC embedding (`Harmony` embeddings), which encourages spots to group by spot type rather than by dataset-specific conditions (**Figure S4**) The above mentioned algorithm was implemented by employing the `HarmonyMatrix()` function from the `Harmony` v.0.1.0 R package ^46^ on a matrix containing reduced dimension coordinates for Visium spots in PC space constructed with the `reducedDim()` function from the `SingleCellExperiment` v.1.18.0 R Bioconductor package ^106^. This results in a QCed and batch-corrected *spe* object.

### 4.7 Spatial domain detection

To enable inspection of the *spe*, we generated an interactive shiny v.1.7.5 R package web application at https://libd.shinyapps.io/Lifespan_DG/ using `spatialLIBD` v.1.12.0 R Bioconductor package ^107^. A blinded experimenter (A.R.P.) manually assigned spots to anatomical domains following consideration of marker gene expression and histological staining (**Figure S7**). Simultaneously, generation of unsupervised spatial domains were performed on the *spe* object by a separate experimenter (A.D.R.) using the `spatialCluster()` function from the `BayesSpace` v.1.6.0 R Bioconductor package ^47^ as previously described ^48^ (**Figure 1c, Figure S5**). Briefly, the number of clusters are determined *a priori* by biological/anatomical knowledge and fixed prior to spatial clustering. For each Visium spot, a low dimensional representation of the gene expression vector is obtained from the `Harmony` embeddings. Bayesian priors are determined by an initial non-spatial clustering with `Mclust` *^108^* and compared with a Markov random field given by the Potts model, which encodes information on all the spots and their neighboring spots; this allows for smoothing of initial clusters by encouraging neighboring spots to be grouped in the same cluster. The resulting Bayesian model is a fixed precision matrix model, where iterative Gibbs sampling is used for updating most of the parameters in the Metropolis–Hastings algorithm; Markov chain Monte Carlo (MCMC) method is used to update latent clusters iteratively produced by the Potts model. The mode (average) of the chain for each cluster label of a spot is assigned as the final cluster label of that spot. The number of repetitions was set empirically via trial by trial basis. We chose *k*=10 as the number of clusters and ran `BayesSpace` at 50,000 iterations. At *k* > 10 we saw less smoothing of spatial domains and bifurcation of the GCL into two clusters with mixing of the ML. Additionally, we noted that some of the capture areas contained thalamic regions, which are enriched for inhibitory cell markers and partially included in spatial domain 6 (**Figure S5**, **Figure S6**). Since this could interfere with differential expression results pertaining to the SGZ, we set a threshold of logcount < 1 for expression of the pan-thalamic marker *TCF7L2*, which removed virtually all thalamus-containing Visium spots (**Figure S5b**). A total of 65,782 spots were included in pseudobulk and differential expression analysis. To assess cluster neighborhood purity for each Visium spot, we used the `neighborPurity()` function from the `bluster` v.1.10.0 Bioconductor package, which uses a hypersphere-based approach to compute the “purity” of each cluster based on the number of contaminating spots from different clusters in its neighborhood (**Figure S8**).

### 4.8 Spatial domain-level processing

For gene modeling and functional enrichment analyses, the spots were pseudo-bulked by the generated spatial domain and donor, as previously described ^103^. Briefly, taking the QCed and batch-corrected *spe* object, we summed the raw gene-expression counts, for a given gene, across all spots in a given donor and a given spatial domain, and repeated this procedure for each gene with the `aggregateAcrossCells()` function from the `scuttle` package. We filtered for genes that have statistically sufficiently large counts in the pseudo-bulked spatial domains with the `filterByExpr()` function and calculated log normalized counts with the `calcNormFactors()` function, both functions from the `edgeR` v.3.38.4 R Bioconductor package ^109^. We also filtered for pseudo-bulked low Visium spot count by setting a threshold for >50 spots. Principal component analysis of the pseudo-bulked spots revealed that `BayesSpace` domain 3 had variation in many of the principal components that separated it from the other clusters, thus minimizing the variance between HPC spatial domains (**Figure S9**). Examination of cluster 3 gene markers suggested that this cluster was choroid plexus (CP) (**Figure 1c, Figure S6)**. To prevent masking of variance within the HPC proper, `BayesSpace` domain 3 was removed from downstream analyses.

### 4.9 Spatial domain-level gene modeling of age groups

Using the pseudo-bulked spatial domain-level data, spatial domains of interest were isolated or combined and the `pseudoBulkDGE()` function, from the `edgeR` v.3.38.4 R Bioconductor package, was used following the limma-voom method with `eBayes` for differential modeling comparing one age group to all of the other age groups. We computed Student’s *t*-test statistics, log_2_FC, and adjusted *p*-values.

For comparing differential expression across the pseudo-bulked spatial domain-level data, gene modeling was performed using the `spatialLIBD` package. Briefly, the `BayesSpace` spatial domain labels were set as the `registration_variable`. The `registration_model()` function was used to define the statistical model for computing the block correlation, with age and sex as covariates. The `registration_block_cor()` function was used to compute the block correlation using the donor sample IDs as the blocking factor. Then the functions `registration_stats_enrichment()`, `registration_stats_pairwise()`, `registration_stats_anova()` were used, employing the limma-voom method with `eBayes`, to fit enrichment, pairwise, and ANOVA models, respectively.

### 4.10 Functional enrichment analyses

Lists of genes with adjusted *p*-values < 0.05 were compiled from pseudo-bulked spatial domains and age groups after DE analysis with `pseudoBulkDGE()`(**Table S3**, **Table S5**). Each list of genes were computed with Over Representation Analysis (ORA) ^110^ to determine whether known biological functions are over-represented in each spatial domain or age group, by large gene expression differences, for the follow aspects: cellular component (intracellular locations where gene products are active, CC), molecular function (documented molecular activities associated with gene products, MF), and biological processes (sets of pathways and broader biological functions made up of the activities of multiple gene products, BP, **Table S4**). ORA was computed with the `enrichGO()` function as an argument within the `compareCluster()` function from the `clusterProfiler` v.4.4.4 R Bioconductor package ^111^.

We compiled a list of differentially expressed genes for superfine cell class GC.3 versus all other superfine cell classes, and GC.4 versus all other superfine cell classes, using the pseudobulked DE results taken from Nelson et al., 2024 ^59^ with FDR < 0.05, along with their logFC values (**Figure S24**). Each list of ordered genes with their logFC were further subdivided into lists with positive logFC and lists with negative logFC, to represent upregulation and downregulation. They were then computed with gene set enrichment analysis (GSEA) for all ontology categories (CC, MF, and BP). GSEA was computed with the `gseGO()` function as an argument within the `compareCluster()` function from the `clusterProfiler` v.4.4.4 R Bioconductor package.

### 4.11 Single-molecule fluorescent in situ hybridization (smFISH), imaging, and analysis

Infant (*n*=1) and adult (*n*=1) hippocampal sections (10μm) were fixed in 10% neutral buffered formalin (catalog no. HT501128, Sigma-Aldrich) for 30 minutes at RT, followed by ethanol-based serial dehydration. Hybridization assays were performed according to manufacturer instructions using the RNAscope Multiplex Fluorescent Reagent Kit V2 (catalog no. 323100, Advanced Cell Diagnostics [ACD]) and the 4-plex ancillary kit V2 (catalog no. 323120, ACD). Briefly, sections were pretreated with hydrogen peroxide for 10 minutes and then permeabilized with Protease IV for 30 minutes at RT. Probes for *PROX1* (catalog no. 530241, ACD), *LAMP5* (catalog no. 487691-C2, ACD), *GAD1* (catalog no. 404031-C3, ACD), and *SLC17A7* (catalog no. 415611-C4, ACD) were applied to the slide and allowed to hybridize for 2 hours at 40 °C. Slides were washed briefly and stored in 4x saline sodium citrate (catalog no. 351–003-101, Quality Biological) overnight at 4 °C. The next day, probes were amplified and fluorescently labeled with Opal dyes as follows: *PROX1* was labeled with 1:500 Opal 520 (catalog no. FP1487001KT, Akoya Biosciences [AB]), *LAMP5* was labeled with 1:500 Opal 690 (catalog no. FP1497001KT, AB), *GAD1* was labeled with 1:500 Opal 570 (catalog no. FP1488001KT, AB) and *SLC17A7* was labeled with 1:500 Opal 620 (catalog no. FP1495001KT, AB). DAPI was applied to each slide for 20 seconds prior to mounting with Fluoromount-G (catalog no. 00-4958-02, ThermoFisher).

Slides were imaged on an AX Nikon Ti2-E confocal fluorescence microscope equipped with NIS-Elements (v5.42.02). The DG, from the ML to the CA4, was captured with a combination of tiles (30-60 tiles/image, 2048 x 2048 pixels per tile) and z-stacks (7 steps, 2μm/step, 12μm range) at 20x magnification (Nikon PLAN APO λ 20x/0.80) with a pinhole of 1.0 AU. Fluorescently-tagged probes were captured using a custom 6-channel, 3-track experiment setup that includes DAPI (405nm excitation laser, 420-500nm filter), Opal 520 (488nm excitation laser, 500-535nm filter), Opal 570 (561nm excitation laser, 580-600nm filter), Opal 620 (561nm excitation laser, 610-630nm filter), Opal 690 (640nm excitation laser, 675-700nm filter) and a lipofuscin channel (488nm excitation laser, 700-750nm filter). All images were captured using the same laser power (LP) and gain (G) settings as follows: DAPI: 28 LP/5 G, Opal 520: 28 LP/5 G, Opal 570: 6 LP/2 G, Opal 620: 6 LP/0.5 G, Opal 690: 9 LP/1 G, Lipo: 28 LP/15 G. After capture, individual tiles were stitched together prior to max-intensity projecting. Linear unmixing was performed using Opal dye spectral standards and a human post-mortem lipofuscin spectral standard. Opal dye standards were previously created using single-positive *Polr2a* stained, wild-type mouse, coronal sections, where each section was stained singularly with one Opal dye (520, 570, 620, 690). After images were unmixed, they were exported as single channel .tiff files for analysis.

### 4.12 Transfer learning of nmf patterns

We used the `RcppML` v.0.5.5 R package ^112^ for our nmf-based transfer learning as previously described ^59^. Briefly, nmf is a dimensionality reduction technique which for gene expression data generally takes as input a normalized log_2_ counts matrix (genes x observations). This is factored into two lower-rank, orthogonal matrices: one with dimensions genes x *k* (*h*) and one with dimensions *k* x observations (*w*). Here, *k* represents nmf rank, which corresponds with the number of patterns specified by the user. The *h* and *w* matrices correspond with the observation-level and gene-level loadings, respectively, for *k* nmf patterns. For predicting nmf patterns in other scRNA-seq, snRNA-seq, and SRT datasets, we used the `project()` function, which takes the gene-level loadings in *h*, for the shared genes between datasets, as input to impute/predict the *w* (*k* x observations) matrix and thus the cell-level, nuclei-level, or Visium spot-level loadings for all the nmf patterns. Following pattern transfer, nmf pattern weights were normalized to sum to 1 for each transcriptomic dataset. To extract marker genes from nmf patterns we used the `patternMarkers()` function from the `CoGAPS` v3.20.0 Bioconductor package ^113^, which computes a statistic that scores the association of the gene’s values in *h* with a single nmf pattern or linear combination of patterns.

### 4.13 Spot-level deconvolution of cell types

To perform spot-level deconvolution of cell types within each gene expression spot, we used publicly available snRNA-seq data ^31^. To complement the design of this study, which targeted the DG, we truncated the Franjic *et al.*, 2022 data to only the DG (*N*=102,753 nuclei) for use as a reference dataset (**Figure S26**) ^31^ and using `cell2location` v0.1.3 Python package ^65^ as previously described ^104^. Briefly, negative binomial regression was performed to estimate reference cell type signatures. `Cell2location` establishes Bayesian priors of cell abundances by using the Visium spatial and count data, and two manually entered hyperparameters: N_cells_per_location = 5 & detection_alpha = 20. A value of 3 for N_cells_per_location is typically recommended for cortex, but, due to the cell density within the GCL, we increased the estimation (**Figure S15**). To ensure unique gene expression profiles for optimized performance, we collapsed cell subtype clusters for granule cells (GC), CA1, somatostatin inhibitory neurons (InN_SST), parvalbumin inhibitory neurons (InNPV), vasointestinal peptide inhibitory neurons (InN_VIP), *NR2F2* inhibitory neurons (InN_NR2F2), *LAMP5* inhibitory neurons (InN_LAMP5), oligodendrocyte precursors, oligodendrocytes (Oligo), microglia, endothelial cells (Endo), and smooth muscle cells (SMC). Variational Bayesian Inference is employed to produce posterior distributions of estimated cell abundances (**Figure S27, Figure S28**). The mean of these distributions were then assigned to each spot. Cell proportions per spot were derived from the mean cell abundances per spot. To assess cell type heterogeneity of Visium spots, PCA was used to extract the top components of variation for cell type proportions for all spots (**Figure S33**), as previously done to assess cell type heterogeneity in blood ^71^.

### 4.14 Aging gene signature generation and score calculation

Computation of the common aging score (CAS) was performed similarly as previously described ^67^. Using each individual pseudo-bulked spatial domain-level data to control for differences in spatial domains, the `pseudoBulkDGE()` function, from the `edgeR` v.3.38.4 R Bioconductor package, was used following the limma-voom method with `eBayes` for differential modeling comparing each age group (teen, adult, elderly) to the infant age group. We computed Student’s *t*-test statistics, log_2_FC, and adjusted *p*-values. Membership in the aging signature gene set was determined by being a DEG in at least two of the age-based differential modeling with thresholds of adjusted *p*-values smaller than 0.05 and a log_2_ fold change of at least 1.5, and shared by at least 3 spatial domains. The constructed signed gene set assigned genes with a value of 1 if positively associated with aging and -1 if negatively associated with aging. The Visium QCed count data, with no log normalization, was UMI-scaled and, along with the signed gene set, used as input for the Visium function then further processed with the analyze function from the `VISION` package v.3.0.1. Briefly, signature aging scores for every spot is calculated with the analyze function by first log-transformation and removing global cell-specific distributional effects from the signature scores by *Z*-normalizing the expression data, then taking the sum of expression values for positive genes minus the sum of expression values for the negative genes divided by the total amount of genes in the signed gene set. This results in a score that summarizes the contrast between the positive and negative signed gene set (**Table S6**, **Figure S31**).

To construct CAS velocities, linear modeling of CAS from age and spatial domain was performed with the lm function in R. The `lstrends()` function from the `lsmeans` package v.2.30.0 with Tukey’s HSD test within all possible spatial domain-to-spatial domain comparisons was used to estimate and compare the slopes of fitted lines for each spatial domain to assess significant slope differences.

### Statistics and reproducibility

No statistical methods were used to predetermine sample sizes. **Table S1** contains the demographic information for the 17 donors in the study. All box plots display the median as the center, interquartile ranges (IQR) (25th & 75th percentiles) as the box edges, and 1.5× the IQR for the whiskers. All reported *p* values were two sized and were adjusted for multiple testing with Benjamini–Hochberg correction unless otherwise stated. Distributions of the residuals of the linear modeling were assumed to be normally distributed across all genes and models, but this was not formally tested. Wilcoxon signed-rank test used for comparing mean cell abundances between two age groups. Tukey’s HSD test was used across all possible spatial domain-to-spatial domain comparisons of CAS slopes. Spots that were outside of tissue or did not pass QC checks were omitted from all analyses. We used the brain donors as a blocking factor in our analyses, as they were also unique for each Visium capture area. Data collection and analysis were not performed blind to the conditions of the experiments. Plots in R were created either in base R or with the `ggplot2` R package ^114^.

### 4.15 Data Availability

The raw and processed data are publicly available through Zenodo listed at https://doi.org/10.5281/zenodo.10126688 ^115^. The raw data provided through Zenodo include all the FASTQ files and raw image files. The processed data include two *spe* objects: (1) with the untransformed feature counts, and (2) with normalized log_2_ transformed feature counts, batch correction, unsupervised spatial clustering, and cell-type deconvolution.

### Additional resources

To visualize the spot-level Visium data and as a resource to the neuroscience community, we created a shiny ^116^ interactive browser available at https://libd.shinyapps.io/Lifespan_DG/ which is powered by the spatialLIBD v.1.15.4 R Bioconductor package ^107^.

### 4.16 Code Availability

The code for this project is publicly available through GitHub at https://github.com/LieberInstitute/spatial_DG_lifespan ^117^ and is described in the associated README.md file. Analyses were performed using R v.4.2.1 with Bioconductor v.3.15.2. `cell2location` v0.1.3 was employed via the `reticulate` R package v.1.28.

### 4.17 Acknowledgements, Funding, Authorship Contributions

Portions of some figures were generated with BioRender.com. We thank the LIBD neuropathology team, particularly James Tooke and Amy Deep-Soboslay, for curation of the brain samples and assistance with tissue dissections. We thank the staff and physicians at the brain donation sites, and the generosity of the brain donors and their families, without whom this work would not be possible. We thank Andrew E Jaffe, Daniel R Weinberger, and members of the LIBD spatial team for helpful feedback to the manuscript. Finally, we thank the families of Connie and Steve Lieber and Milton and Tamar Maltz for their generous support.

Funding for this project was provided by U01MH122849 (KM), R21AG083328 (SCP and SCH), and the Lieber Institute for Brain Development.

## Authorship Contributions

Conceptualization: ADR, KM, SCP

Data Curation: ADR, HRD, SCP

Formal Analysis: ADR, MT, HRD, SCH, EDN

Investigation: ADR, HRD, ARP, EAP, SCP

Methodology: ADR, SCH, EDN

Validation: ADR, ARP

Resources: JEK, TRH

Software: ADR, HRD, RAM

Visualization: ADR, MT, ARP

Project Administration: KRM, LC-T, KM, SCH, SCP

Supervision: KRM, KM, SCH, SCP

Funding acquisition: KM, SCH, SCP

Writing – original draft: ADR

Writing – review & editing: ADR, KM, SCH, SCP

## 5 Supplementary Materials

### 5.1 Supplementary Tables

**Supplementary Table 1**

Demographic information on brain donors, including information on brain id # (BrNum), age at time of death, slide and capture area for each Visium slide, sex, ancestry, RIN for PFC tissue (Screening RIN PFC, performed at time of collection), post mortem interval (PMI), and psychiatric diagnoses (PrimaryDx).

**Supplementary Table 2**

Manual annotations across all Visium spots performed by a scientist blinded to the results of semisupervised clustering. Contains information on donor tissue sample (sample_id), spatial barcode (spot_name), and manual annotation.

**Supplementary Table 3**

DEGs across aggregated DG spatial domains for each age group vs. all others. Contains information on ENSEMBL gene id (gene_id), gene name (gene_name), protein coding or not (gene_type), p-value, adjusted p-value, and log_2_ fold-change.

**Supplementary Table 4**

All significant upregulated/downregulated GO terms from DG age groups vs. all others. Contains information on age group (Cluster), upregulated or downregulated, accession ID (ID), GO term description, ratio of genes from gene set in GO term (GeneRatio), ratio of genes from all genes (BgRatio), p-value, adjusted p-value, q-values, list of genes in GO term (geneID), and number of genes from gene set in GO term (Count).

**Supplementary Table 5**

DEGs from age groups vs. all others for each separate DG spatial domain and the SLM. Contains information on ENSEMBL gene id (gene_id), gene name (gene_name), protein coding or not (gene_type), p-value, adjusted p-value, and log_2_ fold-change.

**Supplementary Table 6**

Signed gene set that makes up the common HPC aging signature. Contains information on ENSEMBL gene id (gene_id), gene name (gene_name), protein coding or not (gene_type), and sign for positively associated aging genes or negatively associated aging genes.

