## Supplementary Figures S1-S34 for "Spatiotemporal analysis of gene expression in the human dentate gyrus reveals age-associated changes in cellular maturation and neuroinflammation"

### 5.2 | Supplementary Figures

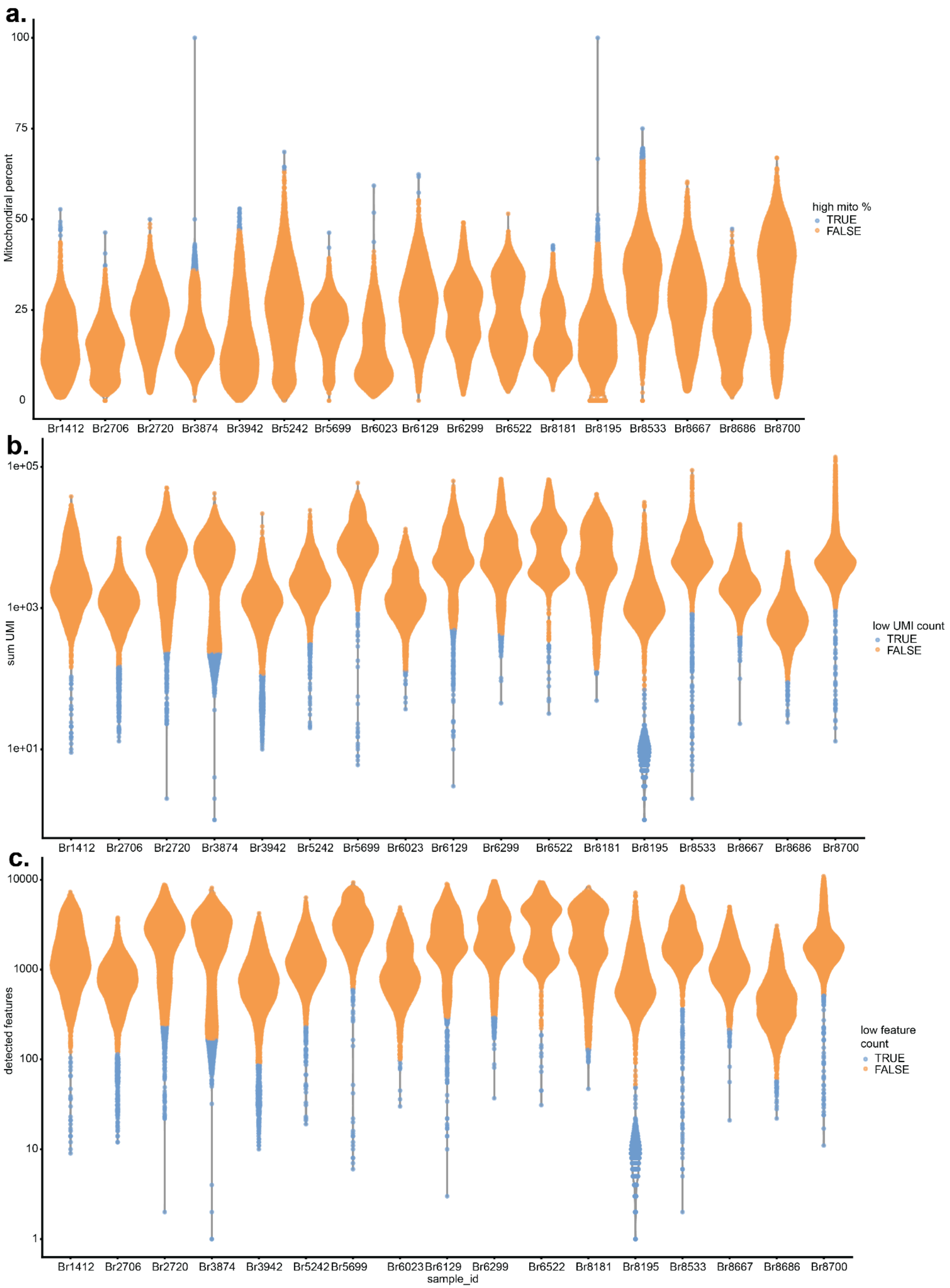

**Figure S1. Quality control (QC) metrics of spot-level Visium data per capture area.** Violin plots for all Visium spots for all samples colored by (**a**) low total UMI count per capture area (library size), (**b**) low number of detected genes or features per capture area, and (**c**) high percent of reads mapping to mitochondrial (mito) genes. The percentage of mitochondrial genes labeled as high for each sample is defined as the percent of the Visium capture area occupied by regions of the HPC; neuropil-rich regions, such as the stratum radiatum (SR) and the molecular layer (ML) have high mitochondrial expression.

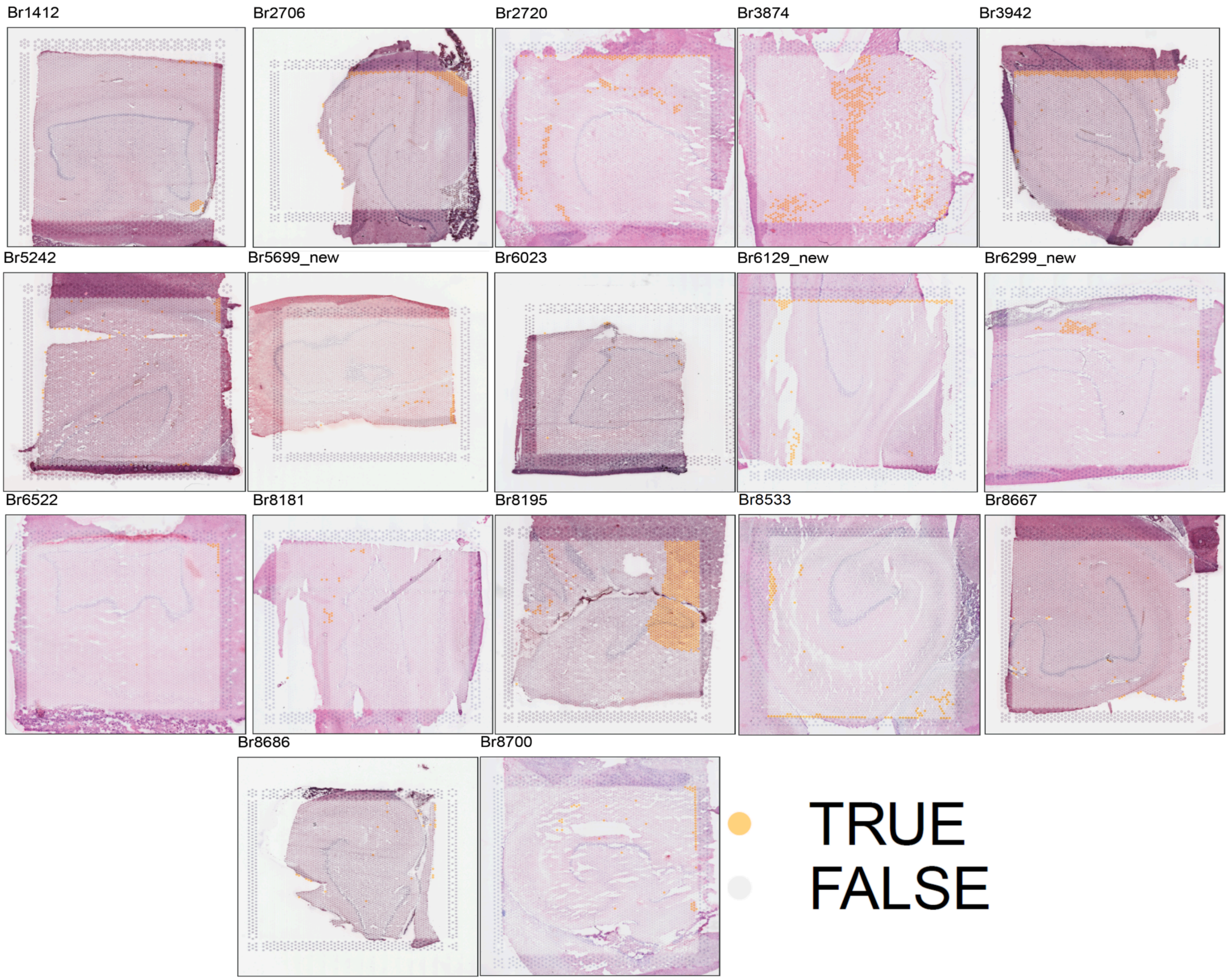


**Figure S2. Visualization of spots that were dropped based on quality control metrics.** Spot plots for all Visium samples showing which spots were discarded (labeled TRUE). Due to low library size, low number of detected features, or high percent of mitochondrial genes. Qualitative assessment of the H&E image for Br3874 revealed that this sample did not contain GCL, and it was thus excluded from downstream analyses.

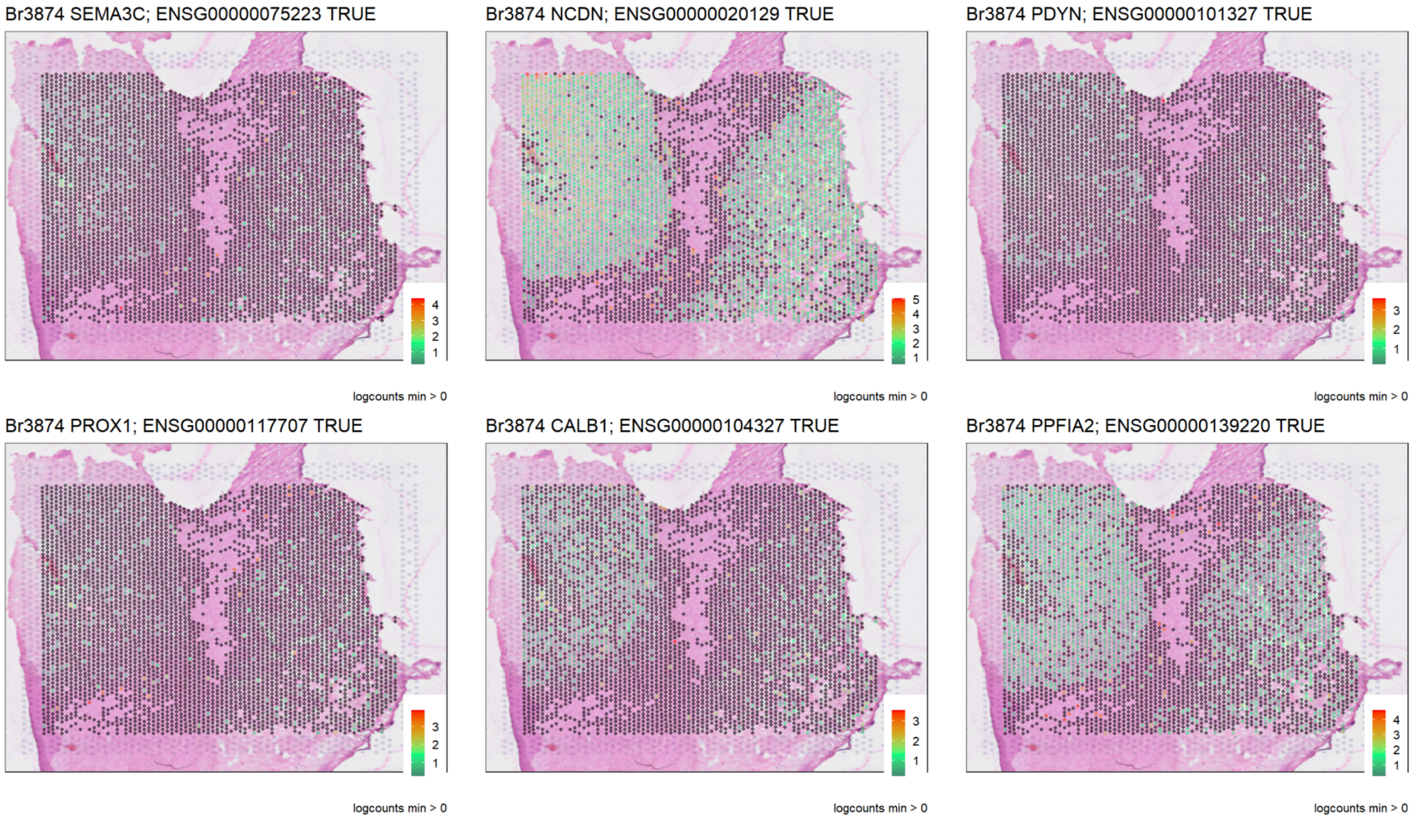


**Figure S3. Gene markers for GCL do not colocalize in Br3874.** Data visualization of the Visium spots of logcounts for canonical GCL markers *SEMA3C, NCDN, PDYN, PROX1, CALB1, and PPFIA2* overlaid onto H&E images. The markers are lowly expressed, do not colocalize consistently, and do not recapitulate known gross anatomy of the GCL.

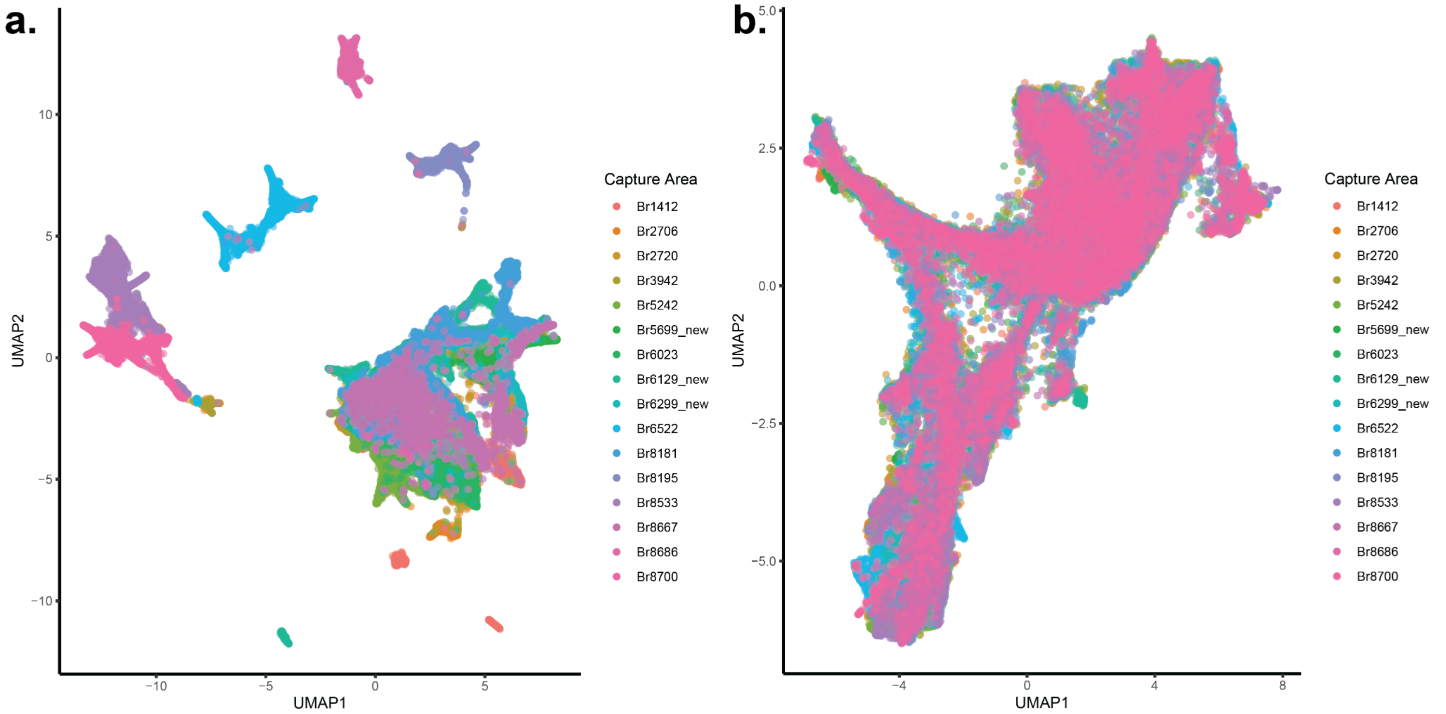


**Figure S4. Spot-level data integration and batch correction with `Harmony`.** (**a**) UMAP plot of all spots colored by donor. (**b**) UMAP plot of all spots colored by donor after data integration and batch correction with `Harmony` [46].

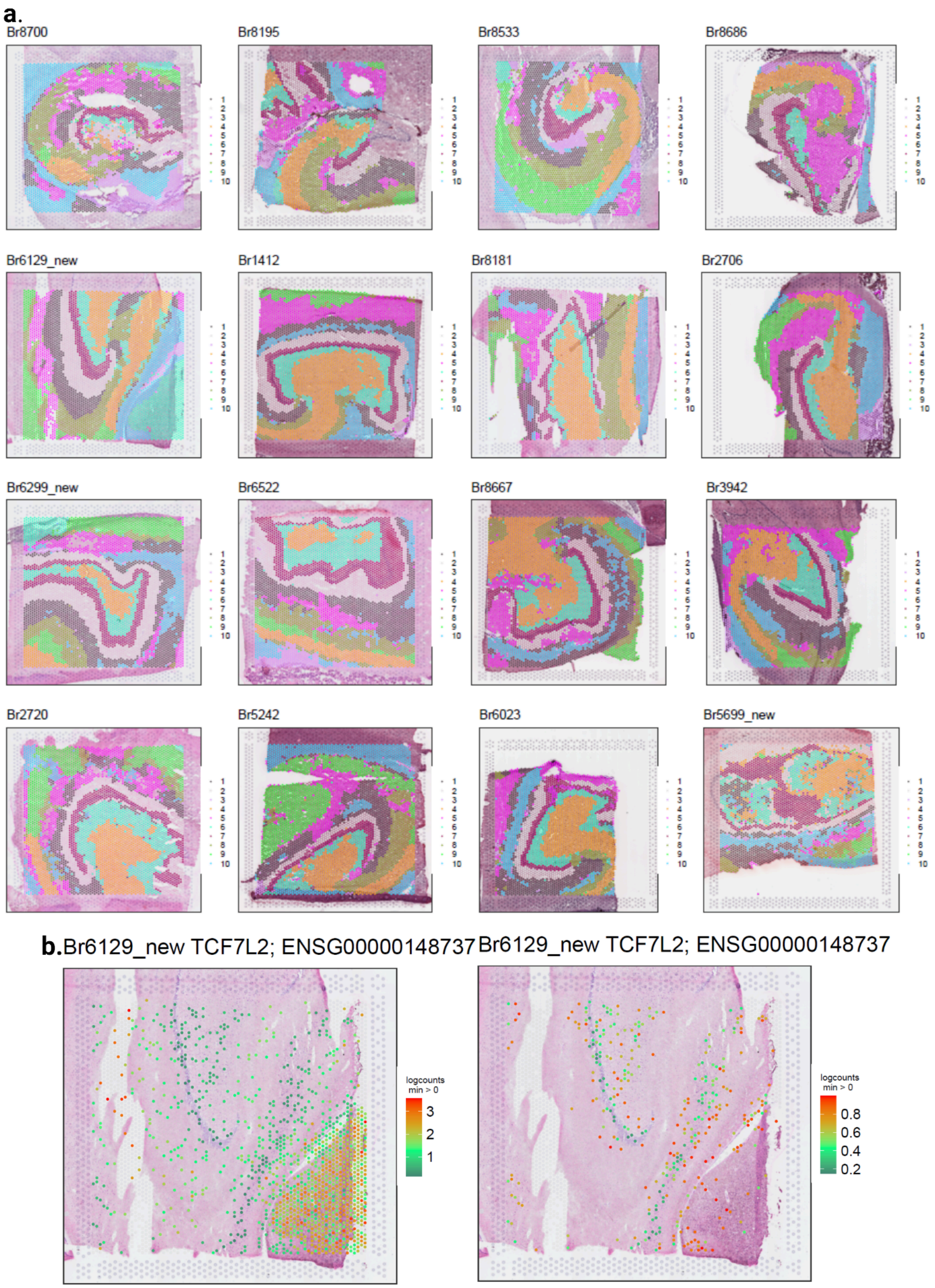


**Figure S5. Data-driven spatial domain clustering using `BayesSpace`.** (**a**) Data visualization for all samples after unsupervised clustering with `BayesSpace` at *k*=10 with 50,000 iterations. Domain numbers and colors are consistent in all capture areas. (**b**) One sample (Br6192_new) included thalamic tissue; a threshold of < 1 logcount for the pan-thalamic gene marker *TCF7L2* was used to remove these spots prior to downstream differential expression-based analyses.

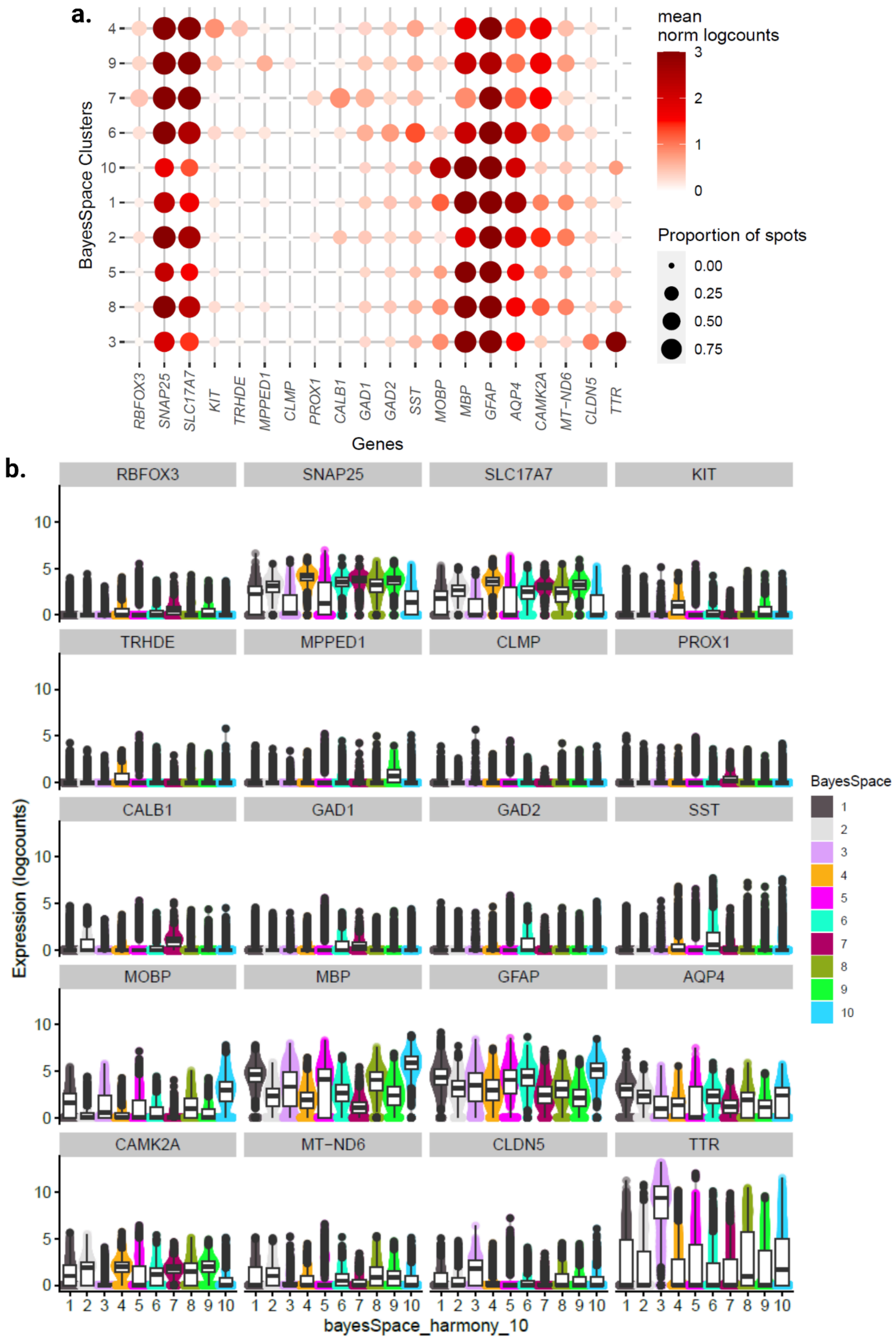


**Figure S6. Canonical gene markers for `BayesSpace` domains.** (**a**) Dot plot colored by mean $\log_2$ normalized counts for canonical domain-specific and neuropil-enriched gene markers averaged across spots (columns) for 10 domains (rows). Circles are sized by the proportion of spots with nonzero expression and colored by mean $\log_2$ normalized counts. (**b**) Violin plots superimposed with box plots for the gene markers from (a). *Y*-axis is $\log_2$ normalized counts and *x*-axis indicates spatial domains.

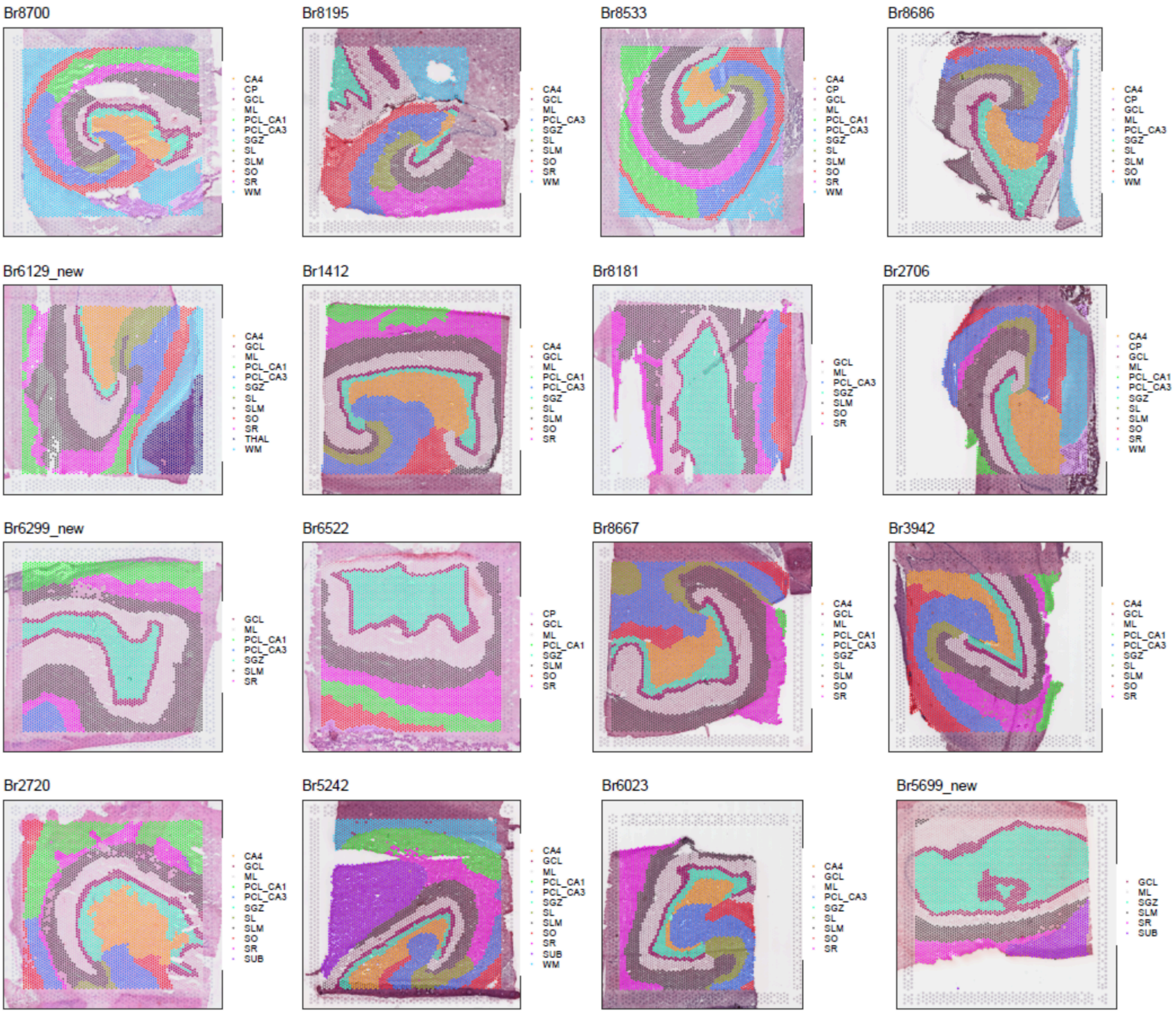


**Figure S7. Manual annotations of spatial domains guided by histology and known marker genes.** Data visualization of Visium spots for all capture areas after performing histologically- and marker gene-guided manual annotations. Spots are colored by manually annotated spatial domains, and are consistent across all capture areas. Annotated spatial domain abbreviations are as follows: cornu ammonis 4 (CA4), choroid plexus (CP), granule cell layer (GCL), molecular layer (ML), principal cell layer cornu ammonis 1 (PCL_CA1), principal cell layer cornu ammonis 3 (PCL_CA3), subgranular zone (SGZ), stratum lucidum (SL), stratum lacunosum moleculare (SLM), stratum oriens (SO), stratum radiatum (SR), subiculum (SUB), and white matter (WM).

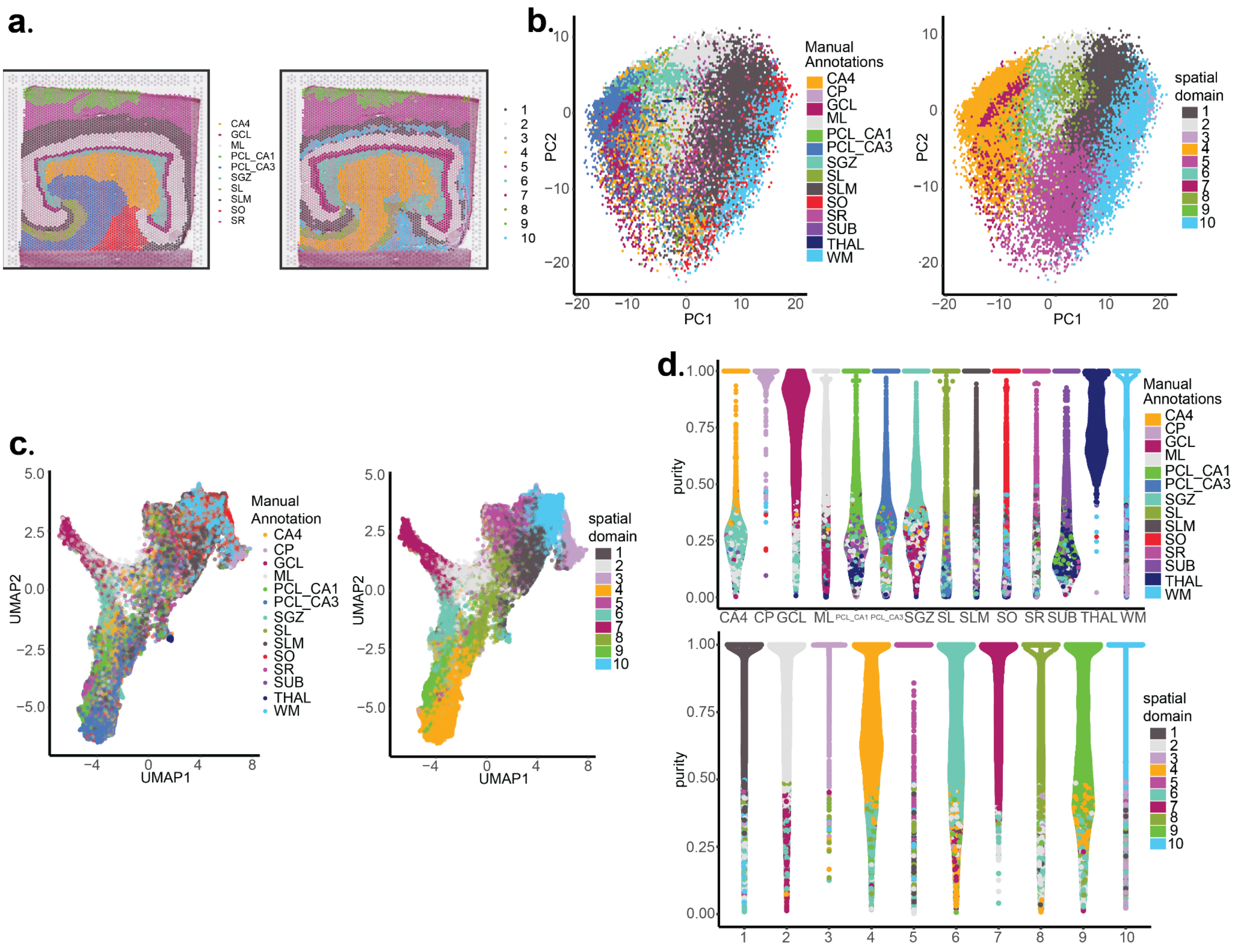


**Figure S8. Comparing `BayesSpace` domains to manual annotated domains.** (**a**) Representative spot plot for manual annotations (left) and unsupervised clustering with `BayesSpace` at *k*=10 (right) of the same tissue sample (Br1412). (**b**) Hexagonal cell representation of the first two principal components (PC1, PC2), with range binned to 100 hexagons (to prevent overplotting) representing the majority label of spots colored by manual annotation (left) or `BayesSpace` domains (right). (**c**) UMAP plots for all spots across all samples qualitatively show better separation of points visually in transcriptomic space with predicted domains (right) compared to manual annotations (left). (**d**) Violin plots of cluster purity for all spots are overall lower for manual annotations (top) compared to *k*=10 `BayesSpace` domains (bottom).

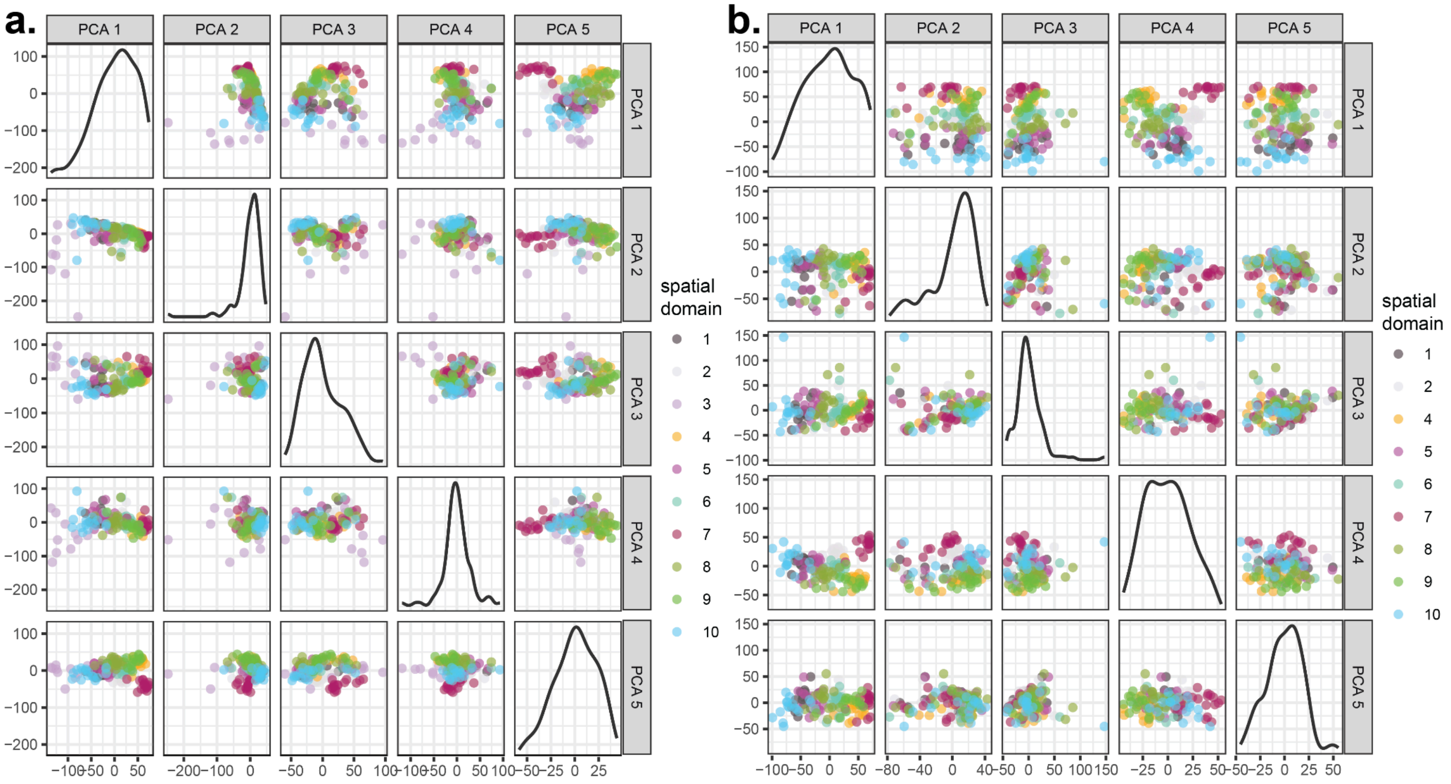


**Figure S9. Choroid plexus compared to non-choroid plexus tissue explains the most variation using principal components analysis.** (**a**) Principal component (PC) plots for the top 6 PCs after pseudo-bulking Visium spots across all `BayesSpace` k=10 domains and *N*=16 donors (total of 160 pseudo-bulked tissue samples); data points are colored by predicted spatial domains 1 to 10. (**b**) Same as (a), but excluding the pseudo-bulked samples containing mostly choroid plexus (predicted spatial domain 3).

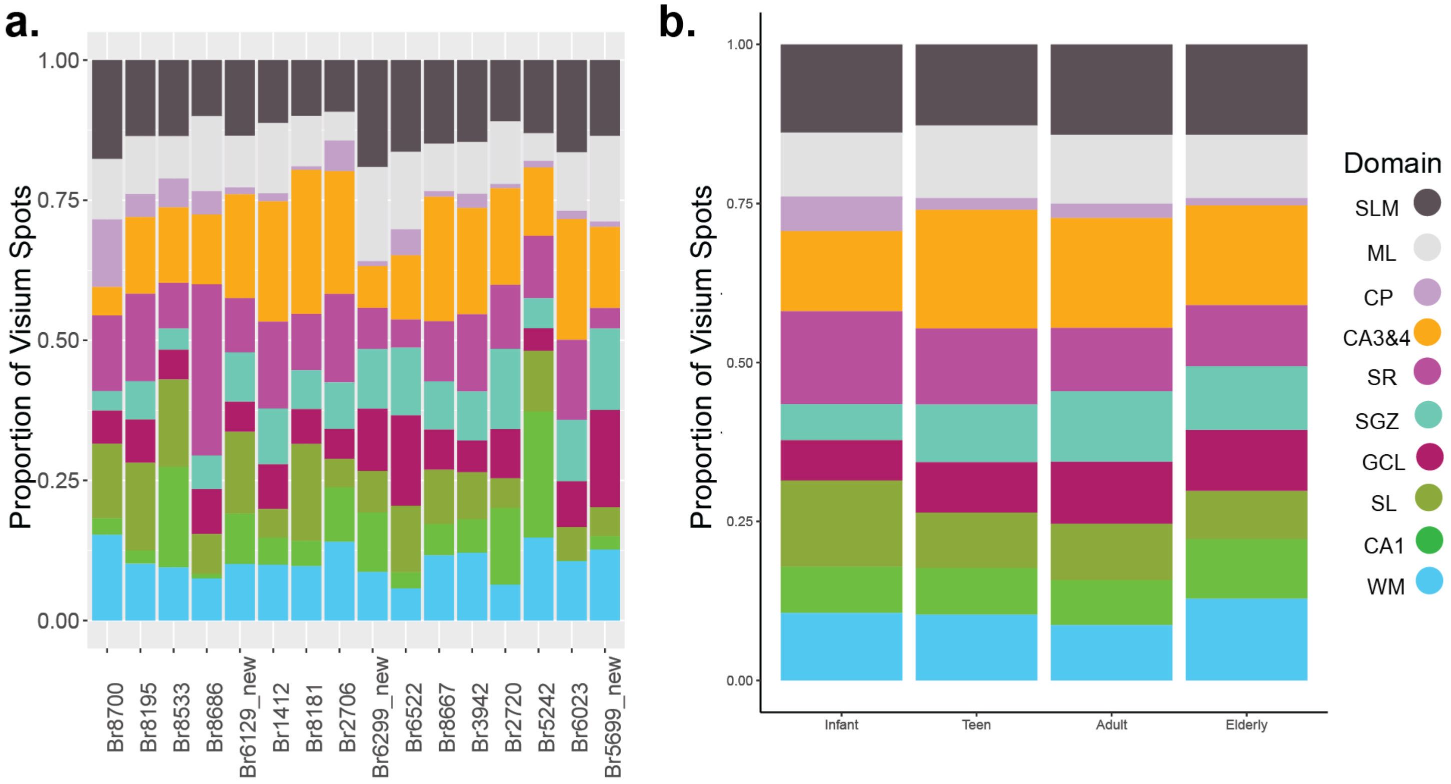


**Figure S10. Proportion of spatial domains.** (**a**) Stacked bar plot illustrating the proportion of spots (*y*-axis) for each `BayesSpace`-predicted spatial domain per donor (*x*-axis). (**b**) Stacked bar plot illustrating the proportion of spots for each `BayesSpace`-predicted spatial domain per age group.

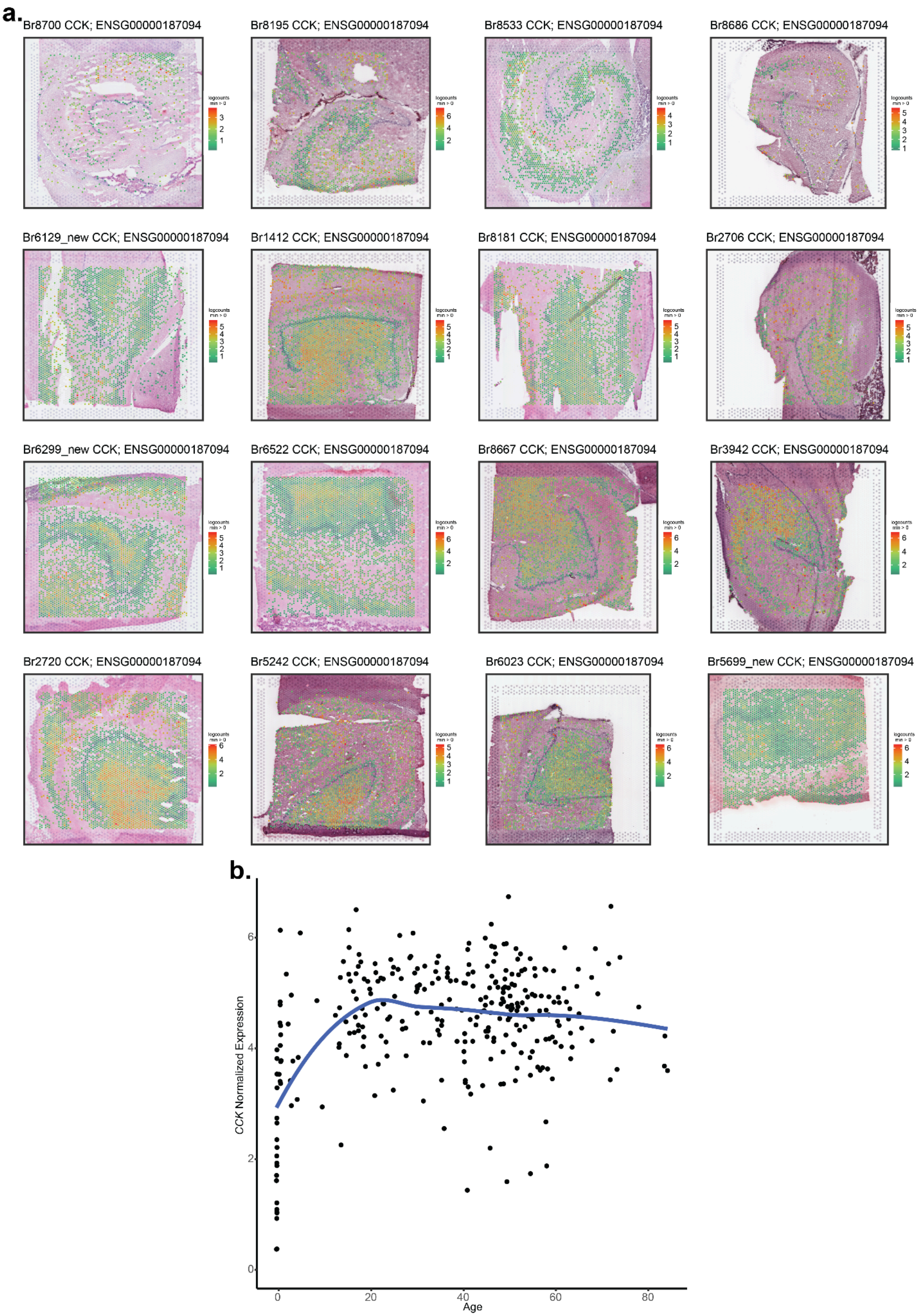
a.
Br8700 CCK; ENSG00000187094
Br8195 CCK; ENSG00000187094
Br8533 CCK; ENSG00000187094
Br8686 CCK; ENSG00000187094
Br6129_new CCK; ENSG00000187094
Br1412 CCK; ENSG00000187094
Br8181 CCK; ENSG00000187094
Br2706 CCK; ENSG00000187094
Br6299_new CCK; ENSG00000187094
Br6522 CCK; ENSG00000187094
Br8667 CCK; ENSG00000187094
Br3942 CCK; ENSG00000187094
Br2720 CCK; ENSG00000187094
Br5242 CCK; ENSG00000187094
Br6023 CCK; ENSG00000187094
Br5699_new CCK; ENSG00000187094
b.
CCK Normalized Expression
Age
0
20
40
60
80
0
2
4
6

**Figure S11. The spatial gene expression of *CCK* across age.** (**a**) Data visualization of the Visium spots with the log normalized counts for *CCK* overlaid onto H&E images. The samples are ordered by age from left to right and top to bottom. (**b**) Scatter plot of $\log_2$ normalized expression (*y*-axis) and age in years (*x*-axis) with local regression line from bulk RNA-seq human HPC data [51]. Each point represents one neurotypical donor.

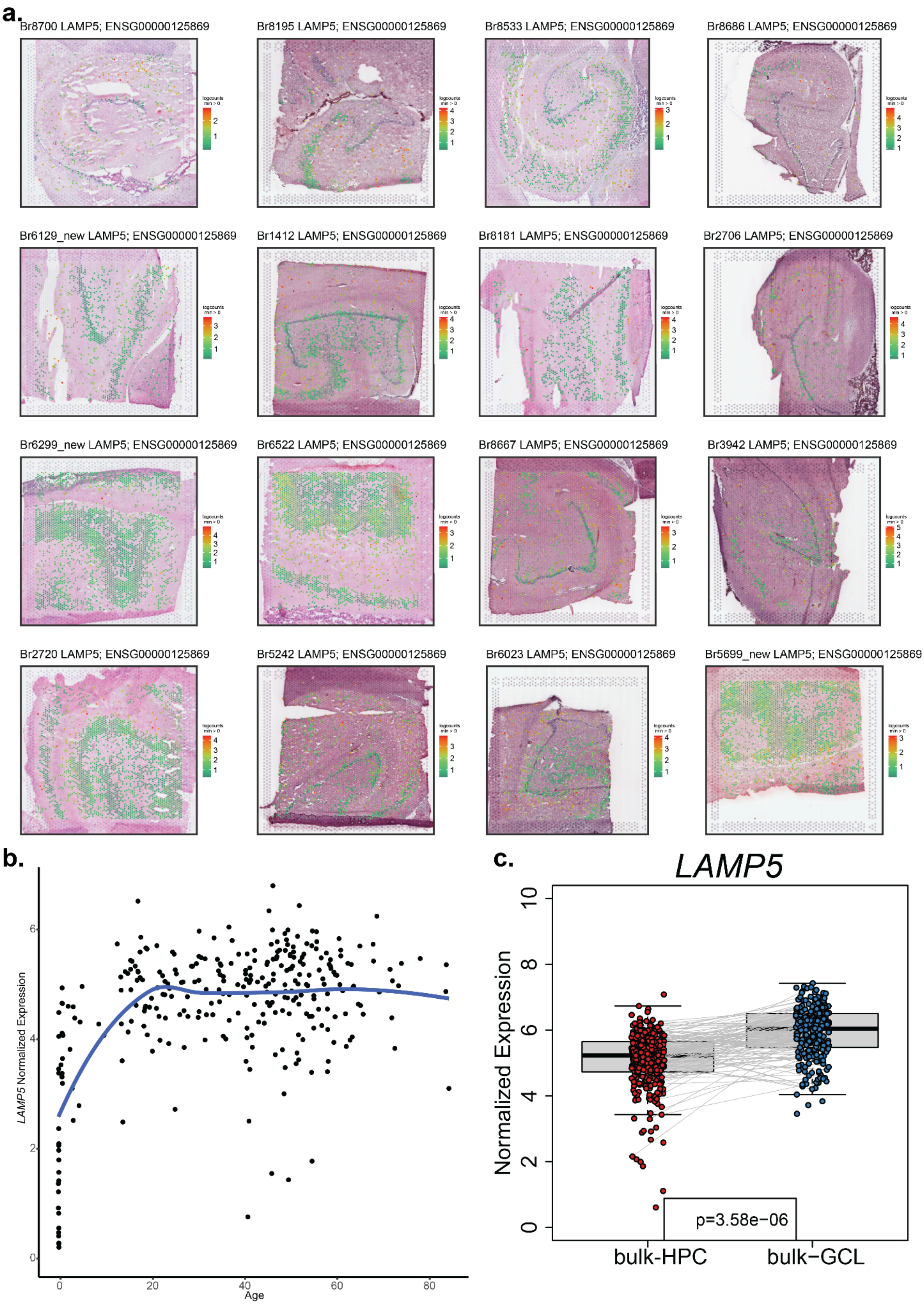
a.
Br8700 LAMP5; ENSG00000125869
Br8195 LAMP5; ENSG00000125869
Br8533 LAMP5; ENSG00000125869
Br8686 LAMP5; ENSG00000125869
Br6129_new LAMP5; ENSG00000125869
Br1412 LAMP5; ENSG00000125869
Br8181 LAMP5; ENSG00000125869
Br2706 LAMP5; ENSG00000125869
Br6299_new LAMP5; ENSG00000125869
Br6522 LAMP5; ENSG00000125869
Br8667 LAMP5; ENSG00000125869
Br3942 LAMP5; ENSG00000125869
Br2720 LAMP5; ENSG00000125869
Br5242 LAMP5; ENSG00000125869
Br6023 LAMP5; ENSG00000125869
Br5699_new LAMP5; ENSG00000125869
b.
LAMP5 Normalized Expression
Age
0
20
40
60
80
0
2
4
6
c.
LAMP5
Normalized Expression
0
2
4
6
8
10
p=3.58e−06
bulk-HPC
bulk−GCL

**Figure S12. The spatial gene expression of *LAMP5* across age.** (**a**) Data visualization of the Visium spots with log normalized counts for *LAMP5* overlaid onto H&E images. Samples are ordered by age from left to right and top to bottom. (**b**) Scatter plot of $log_2$ normalized expression (*y*-axis) and age in years (*x*-axis) with local regression line from bulk RNA-seq human HPC data [51]. Each point represents one neurotypical donor. (**c**) Boxplot of $log_2$ normalized expression comparing reference bulk RNA-seq of HPC and laser dissected GCL data [51,52]. Paired points represent donors that are shared between the datasets.

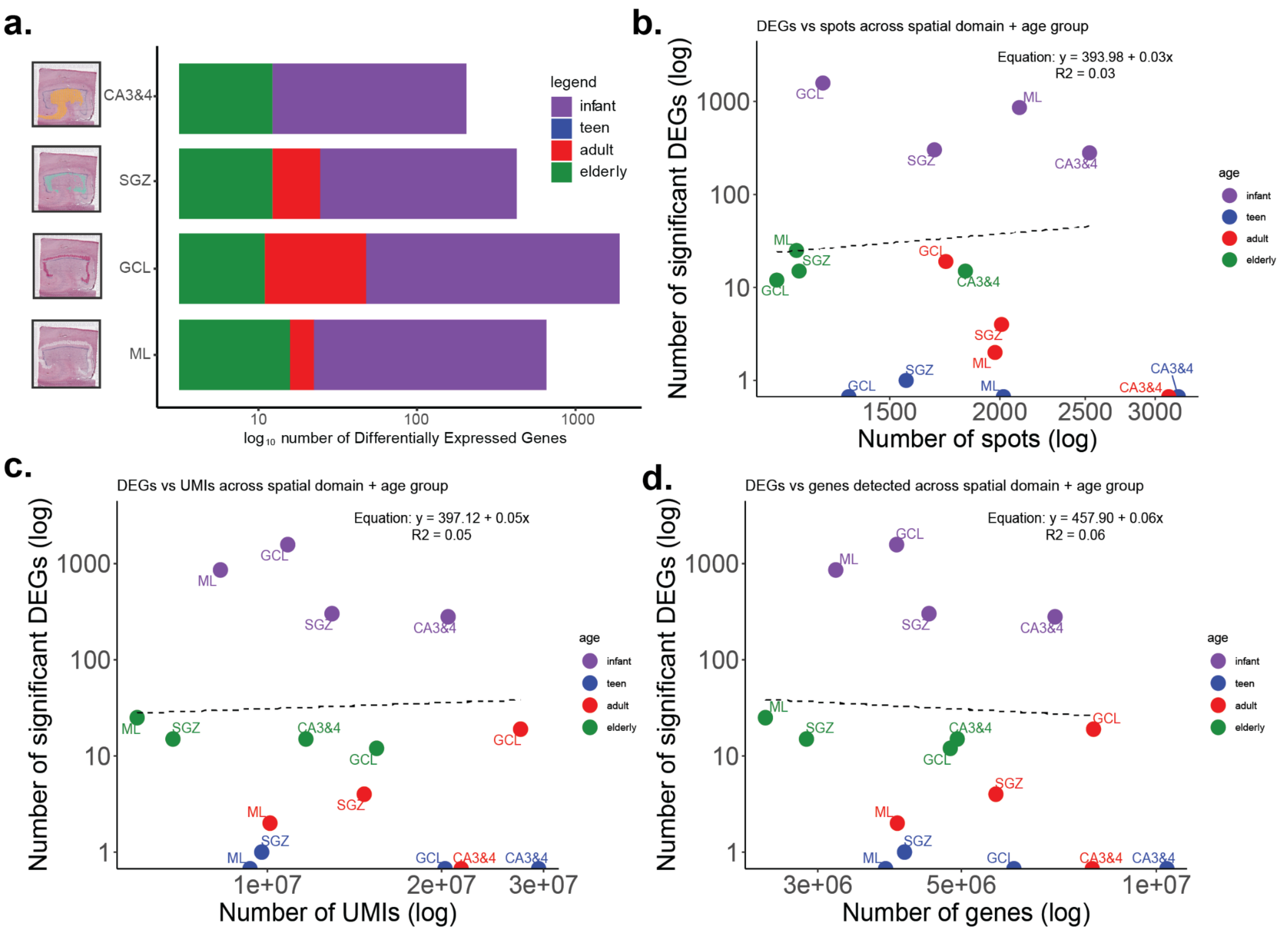


**Figure S13. Number of spots, UMI, and number of genes do not predict the number of DEGs per age group.** (**a**). Stacked barplot of the number of differentially expressed genes (DEGs) for each spatial domain of the dentate gyrus (DG), stacked by age group. *X*-axis on log scale. In the spatial domain-specific DE analyses, these are scatterplots (on log scale for *x*- and *y*-axis) of the number of significant DEGs found (*y*-axis) versus (**b**) spots, (**c**) UMI, and (**d**) detected number of genes (*x*-axis).

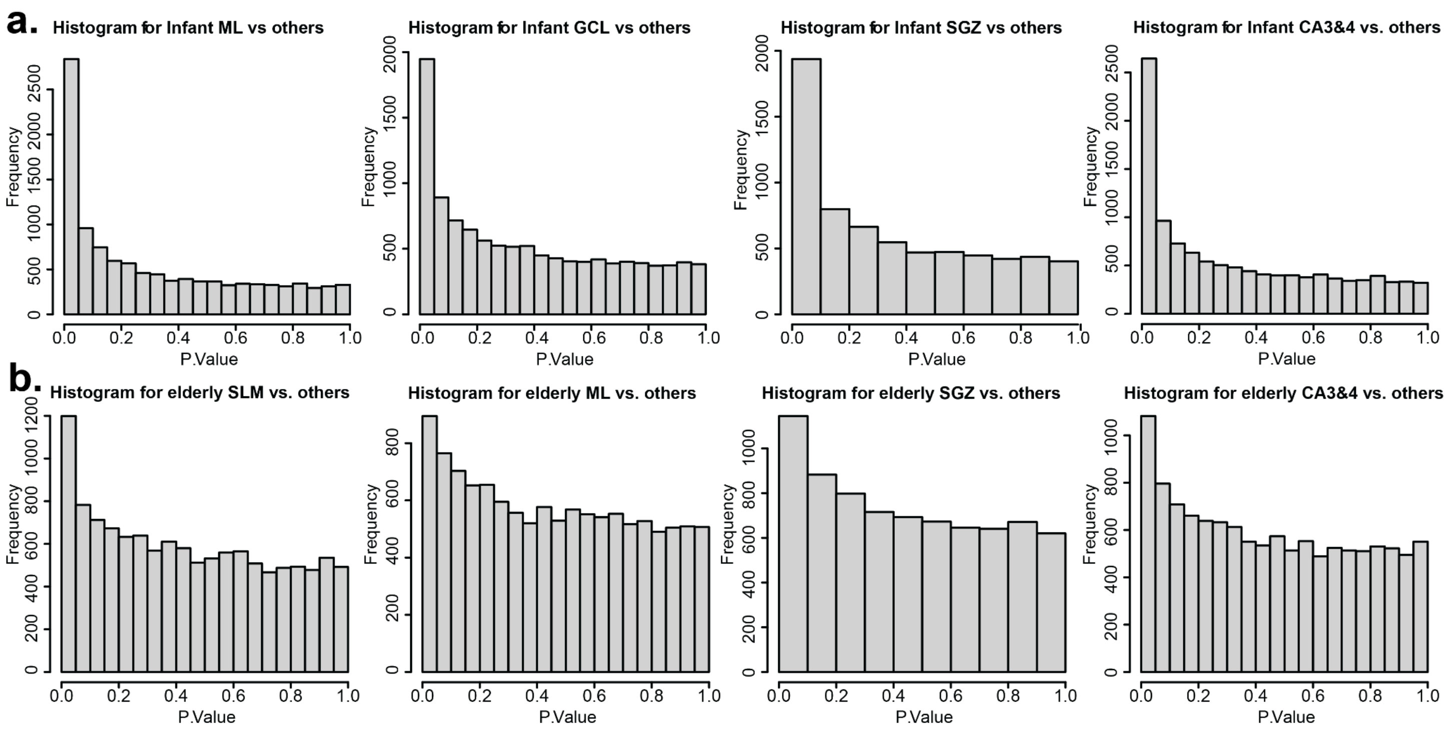


**Figure S14. Anti-conservative distribution of p-values in age comparisons within DG sub-domains for infant and elderly DEGs.** Histograms for the frequency of p-values resulting from pseudo-bulked DE tests comparing one age group to all others. (**a-d)** Infant versus all other age groups within ML, GCL, SGZ, and CA3&4. (**e-h**) Elderly versus all other age groups within SLM, ML, SGZ, and CA3&4.

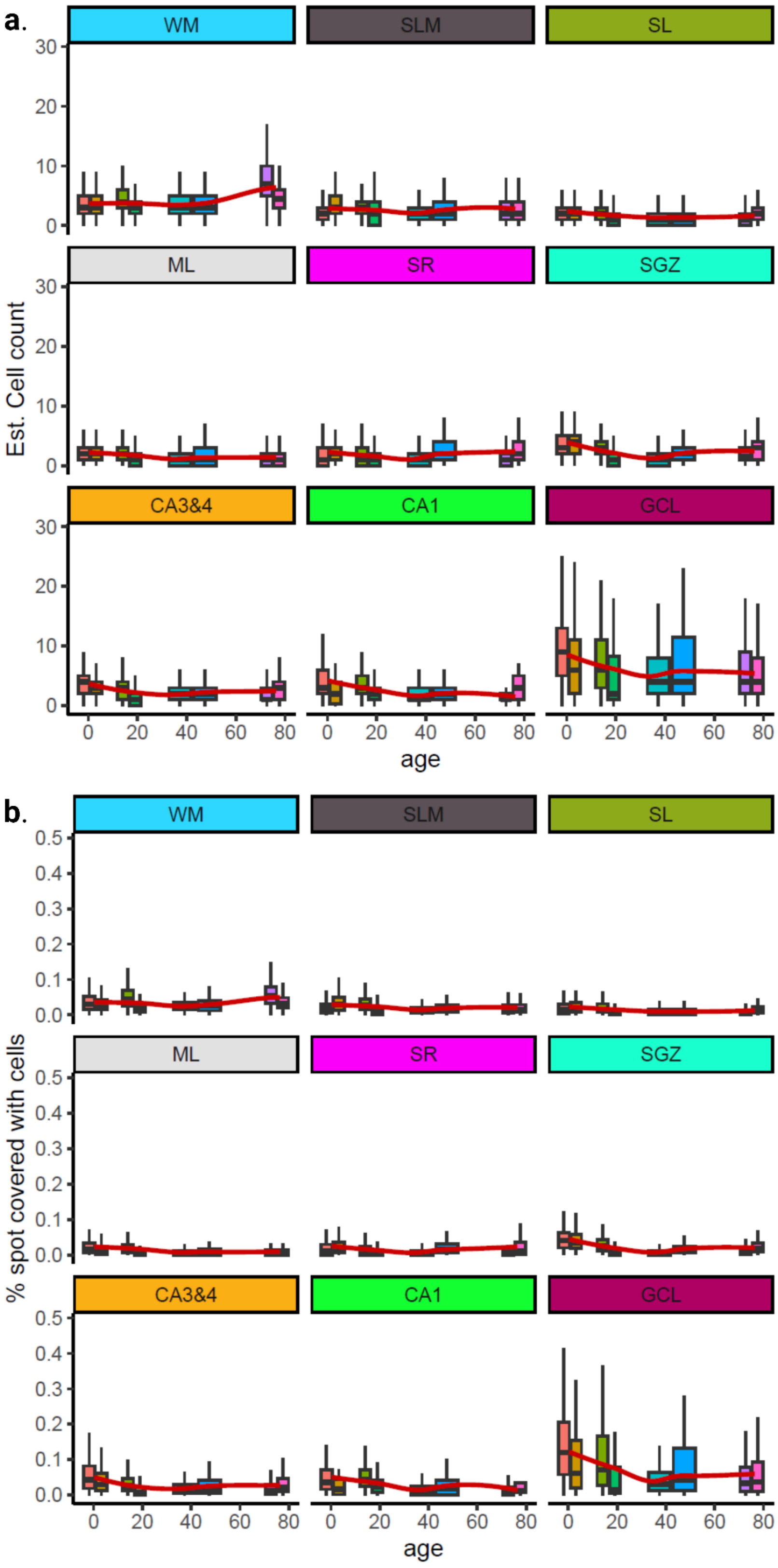


**Figure S15. Spot level nuclei density estimates.** (**a**) Boxplots of the estimated cell count per spot versus age, faceted by HPC spatial domain and fitted with local regression line. (**b**) Boxplots of the percentage of spots covered by nuclei versus age, faceted by spatial domain and fitted with local regression line. Note these data were only calculated from 2 donors per age group (*n*=8 total).

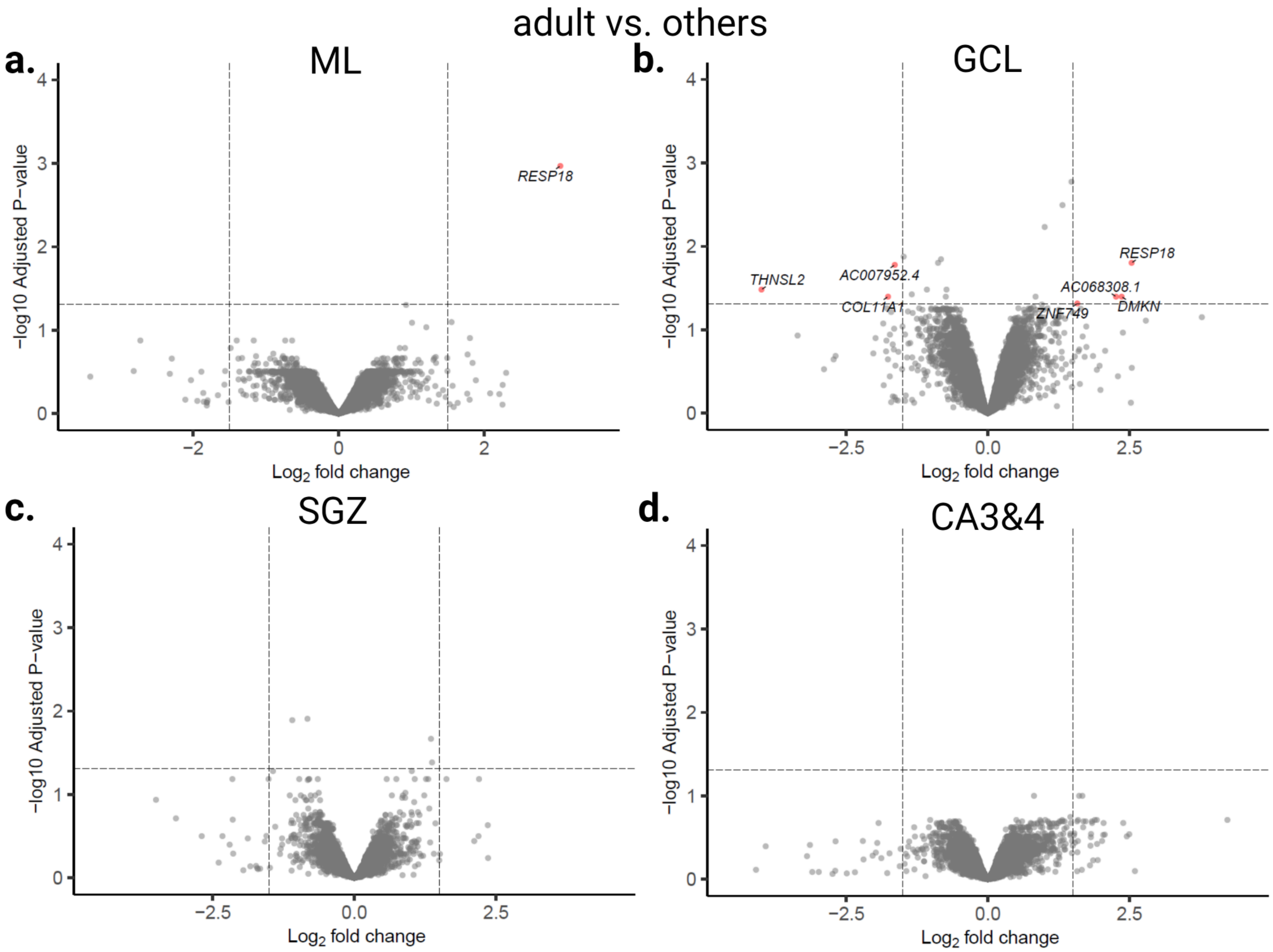


**Figure S16. Number of DEGs in adult DG sub-domains and elderly GCL.** (**a-d**) Volcano plots of adult versus non-adult for each individual pseudo-bulked spatial domain. The *x*-axis is the $\log_2$ fold change in expression and the *y*-axis is the negative $\log_{10}$ adjusted *p*-values.

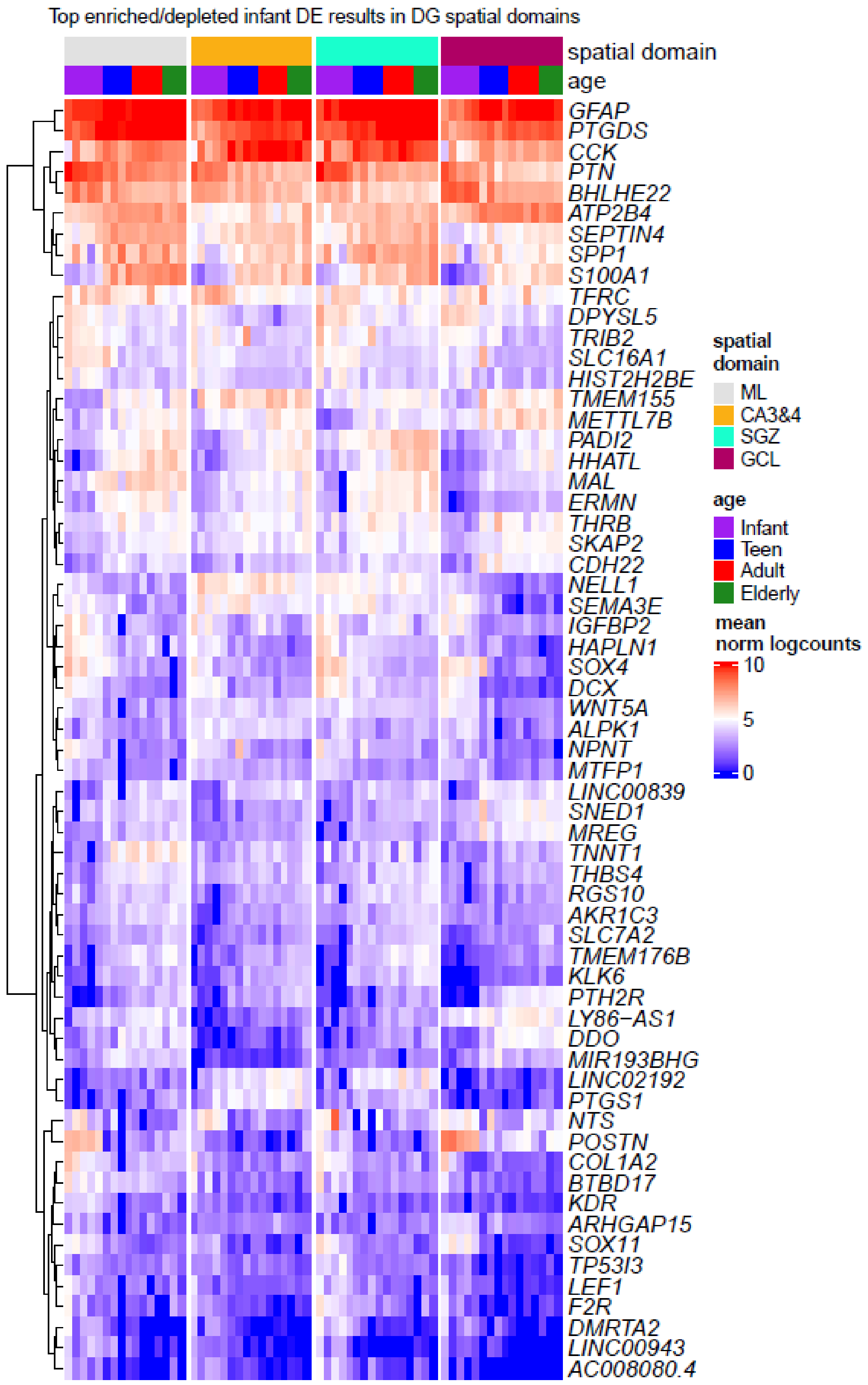


**Figure S17. Top 10 enriched and depleted genes for each spatial domain in infant.** Heatmap of the mean log2 normalized counts for top 10 enriched and depleted genes with ≥1.5 logFC or ≤-1.5 logFC from each spatial domain. Pseudo-bulked data limited to DG sub-domains. Hierarchical clustering was performed across rows. Columns are organized by spatial domains corresponding to DG regions.

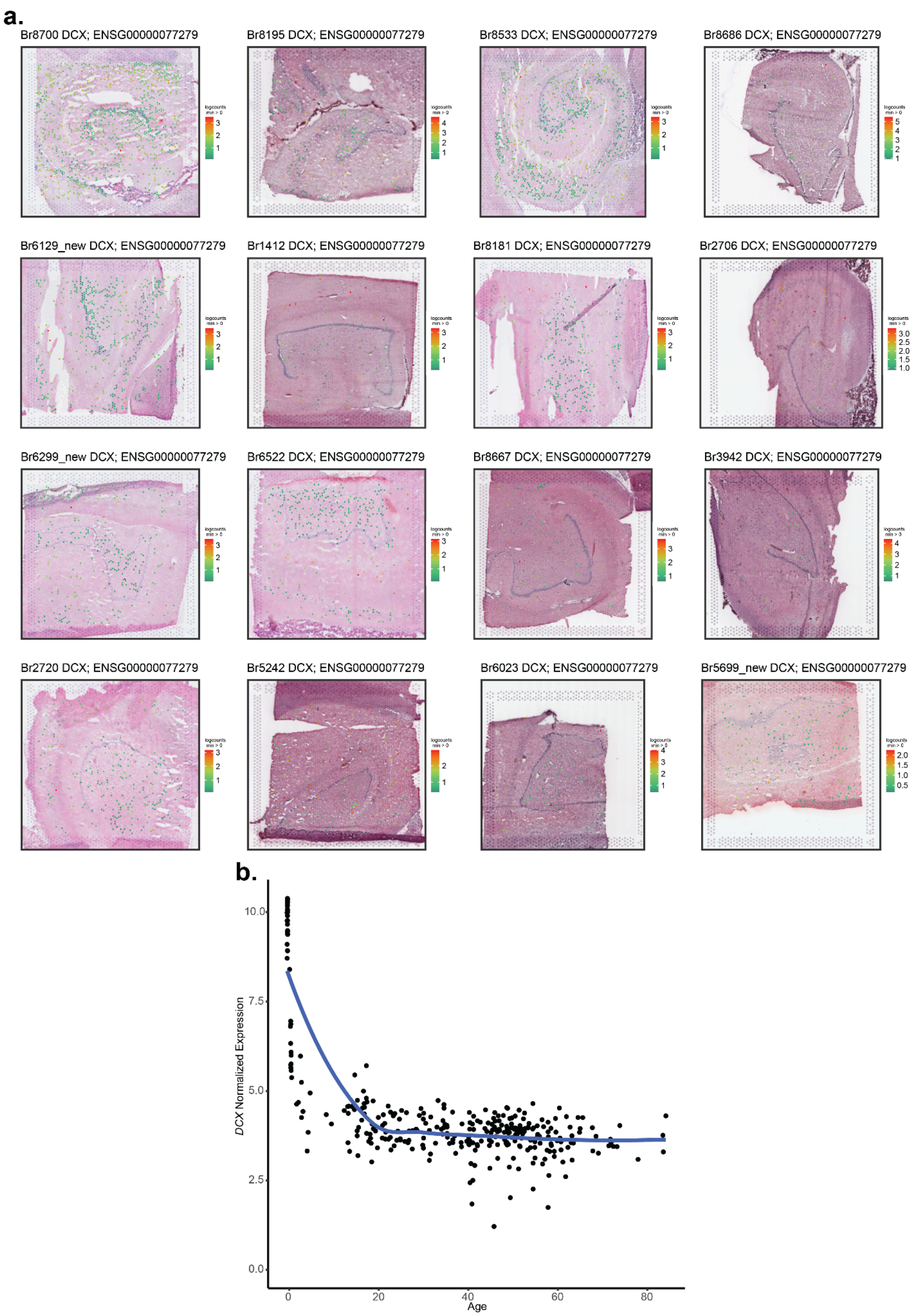
a.
Br8700 DCX; ENSG00000077279
Br8195 DCX; ENSG00000077279
Br8533 DCX; ENSG00000077279
Br8686 DCX; ENSG00000077279
Br6129_new DCX; ENSG00000077279
Br1412 DCX; ENSG00000077279
Br8181 DCX; ENSG00000077279
Br2706 DCX; ENSG00000077279
Br6299_new DCX; ENSG00000077279
Br6522 DCX; ENSG00000077279
Br8667 DCX; ENSG00000077279
Br3942 DCX; ENSG00000077279
Br2720 DCX; ENSG00000077279
Br5242 DCX; ENSG00000077279
Br6023 DCX; ENSG00000077279
Br5699_new DCX; ENSG00000077279
b.
DCX Normalized Expression
Age
0.0
2.5
5.0
7.5
10.0
0
20
40
60
80

**Figure S18. Spatial gene expression of *DCX* across age.** (**a**) Data visualization of Visium spots with the log normalized counts for *DCX* overlaid onto H&E images. Samples are ordered by age from left to right and top to bottom. (**b**) Scatter plot of $\log_2$ normalized expression (*y*-axis) and age (*x*-axis) with local regression line from bulk RNA-seq human HPC data [51]. Each point represents one neurotypical donor.

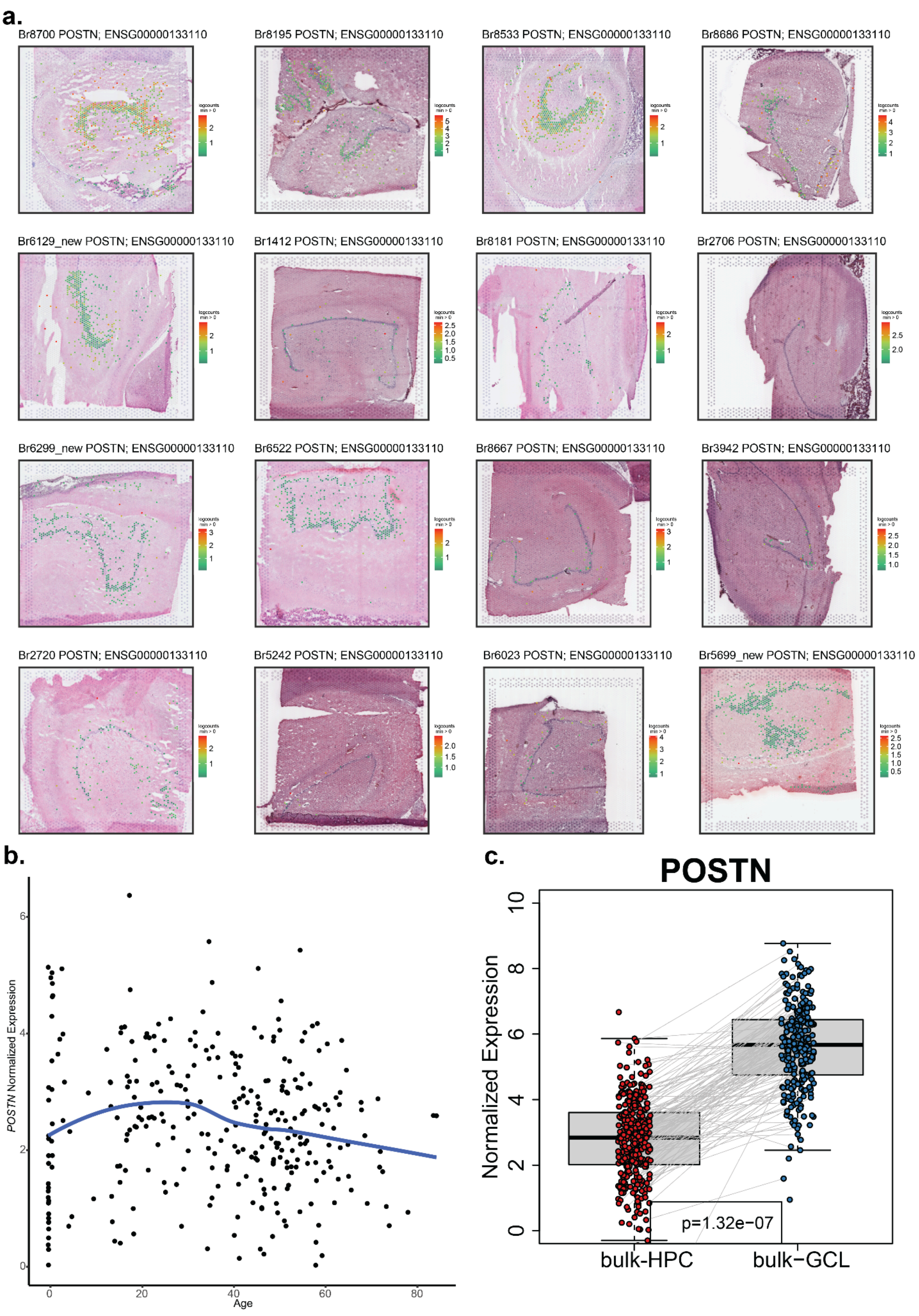
a.
Br8700 POSTN; ENSG00000133110
Br8195 POSTN; ENSG00000133110
Br8533 POSTN; ENSG00000133110
Br8686 POSTN; ENSG00000133110
Br6129_new POSTN; ENSG00000133110
Br1412 POSTN; ENSG00000133110
Br8181 POSTN; ENSG00000133110
Br2706 POSTN; ENSG00000133110
Br6299_new POSTN; ENSG00000133110
Br6522 POSTN; ENSG00000133110
Br8667 POSTN; ENSG00000133110
Br3942 POSTN; ENSG00000133110
Br2720 POSTN; ENSG00000133110
Br5242 POSTN; ENSG00000133110
Br6023 POSTN; ENSG00000133110
Br5699_new POSTN; ENSG00000133110
b.
POSTN Normalized Expression
Age
0
20
40
60
80
c.
POSTN
Normalized Expression
p=1.32e−07
bulk-HPC
bulk−GCL

**Figure S19. Spatial gene expression of *POSTN* across age.** (**a**) Data visualization of the Visium spots with the log normalized counts for *POSTN* overlaid onto H&E images. The samples are ordered by age from left to right and top to bottom. (**b**) Scatter plot of $\log_2$ normalized expression (*y*-axis) and age (*x*-axis) with local regression line of reference bulk RNA-seq HPC data [51]. Each point represents one neurotypical donor. (**c**) Boxplot of $\log_2$ normalized expression comparing reference bulk RNA-seq of HPC and laser dissected GCL data [51,52]. Paired points represent donors that are shared between the datasets.

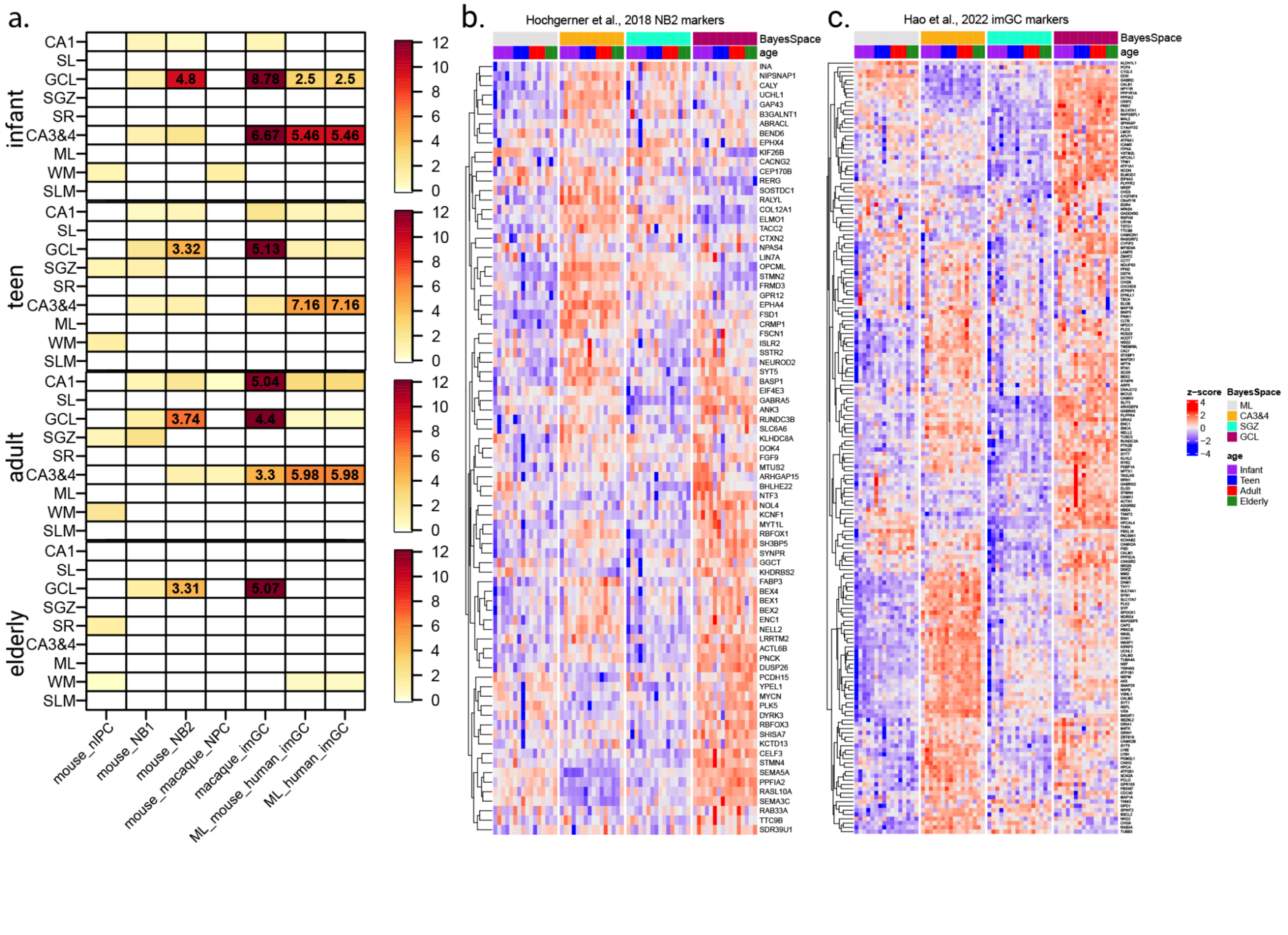


**Figure S20. Gene set enrichment analysis identifies the presence of neurodevelopmentally related**

**gene signatures in the GCL across the lifespan.** (**a**) Enrichment analyses using Fisher's exact tests for predefined gene sets for neural precursor cells (nIPC and NPC), neuroblasts (NB1 and NB2), and immature granule cells (imGC). Nomenclature was adopted from each publication [20,21,56]. Color indicates negative $\log_{10}$ *p*-values while numbers within each significant heatmap cell indicates the odds ratios for the enrichment. (**b**) Heatmap of the mean $\log_2$ normalized counts (centered and scaled) for the mouse NB2 gene set [20] on pseudo-bulked data limited to the DG sub-domains. Hierarchical clustering was performed across rows. Columns are organized by spatial domain. (**c**) Heatmap of the mean $\log_2$ normalized counts (centered and scaled) for the macaque imGC gene set [21] on pseudobulked data limited to the DG sub-domains. Hierarchical clustering was performed across rows, and columns are organized by spatial domain.

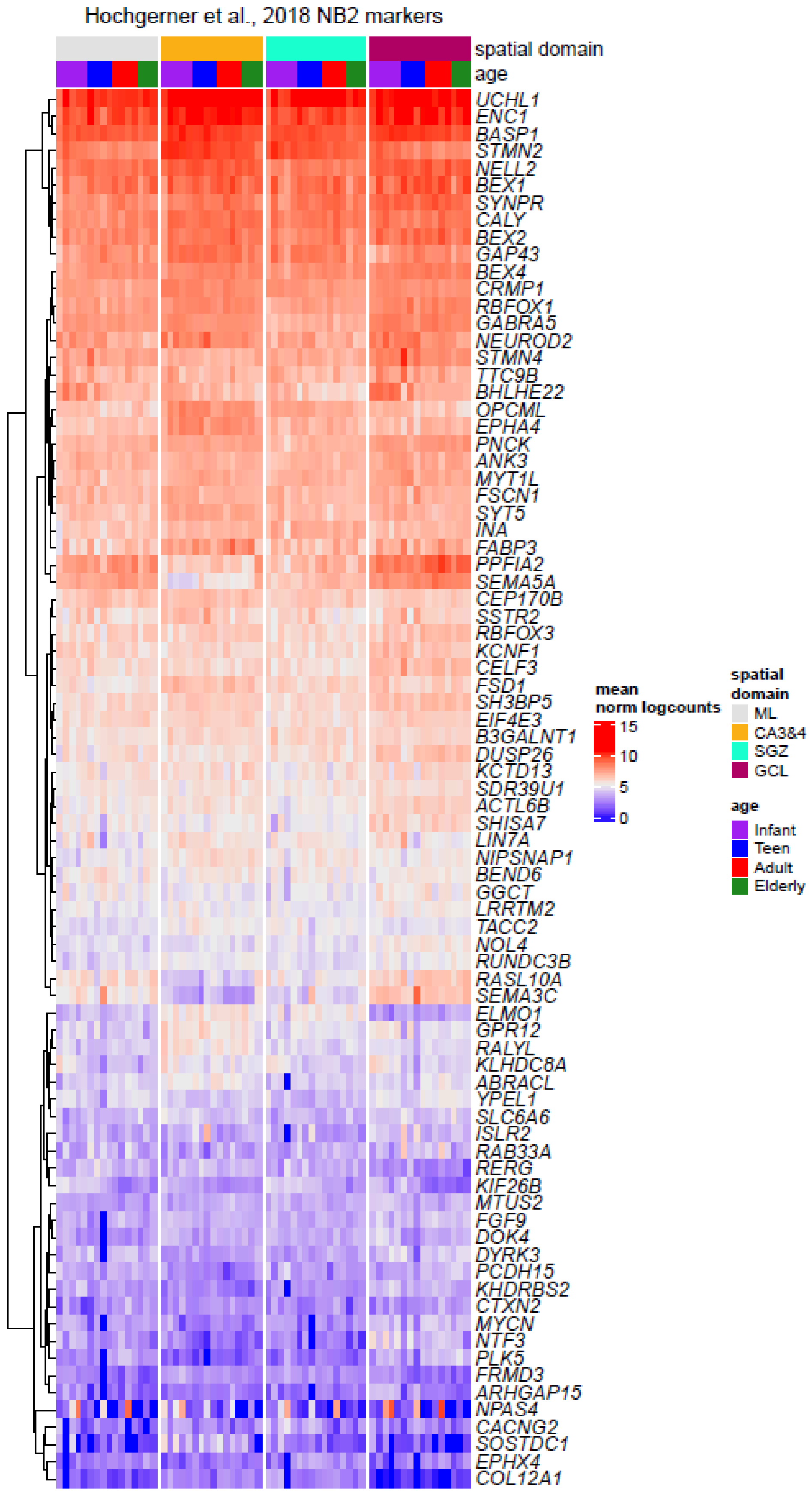


**Figure S21. NB2 gene set expression across pseudo-bulked DG sub-domains.** Heatmap of the mean $\log_2$ normalized counts for the NB2 gene set [20] on pseudo-bulked data limited to DG sub-domains. Hierarchical clustering was performed across rows. Columns are organized by spatial domains.

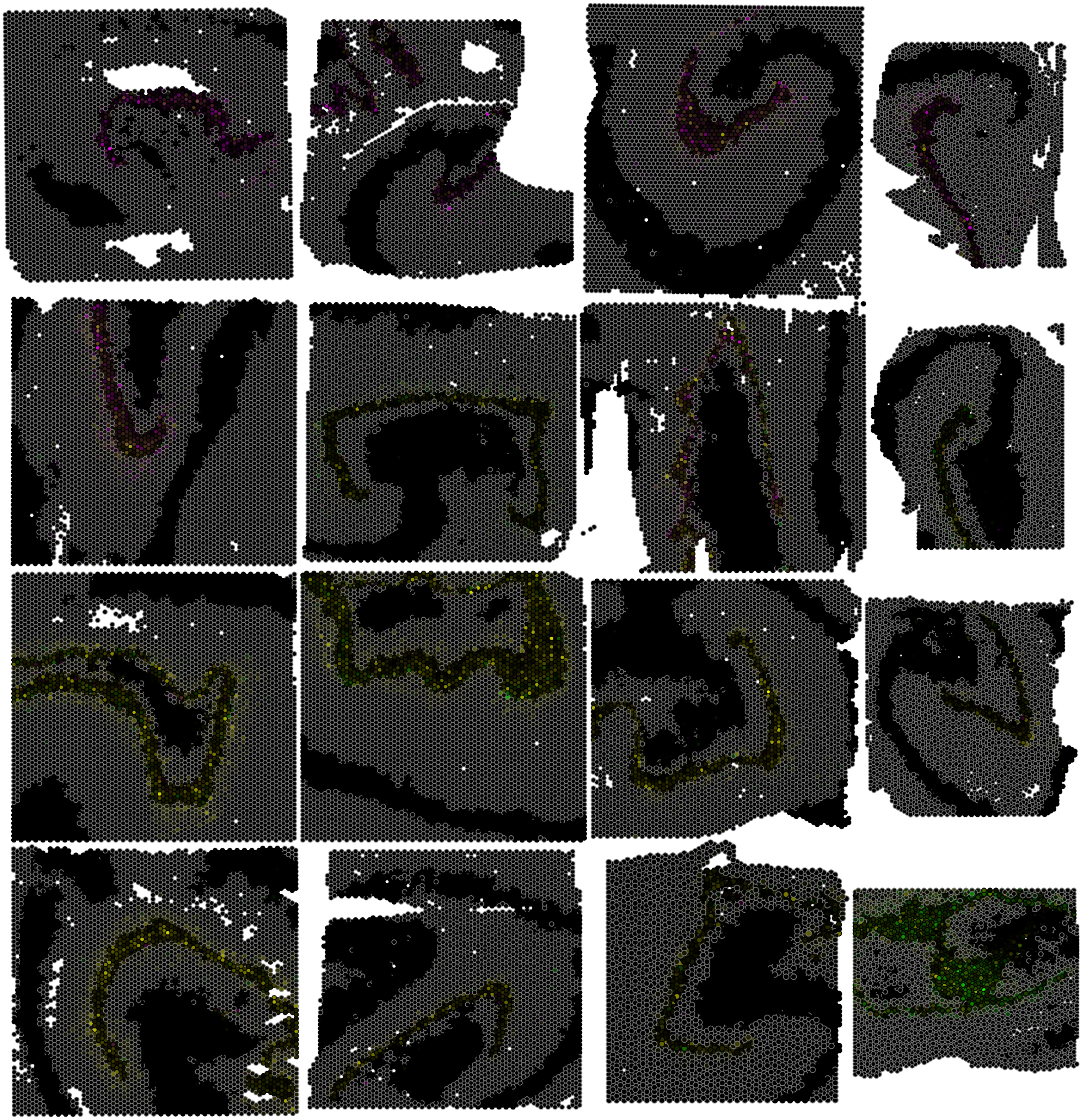

**Figure S22. Spatial gene expression of nmf26, nmf5, nmf14 across age.** (**a**) Data visualization of spot-level weights of nmf26, nmf5, nmf14 across age [59]. Spots are colored by weight of nmf26, nmf5, nmf14. GCL, CA3&4, CA1 spots outlined in black, others outlined in gray. The samples are ordered by age from left to right and top to bottom.

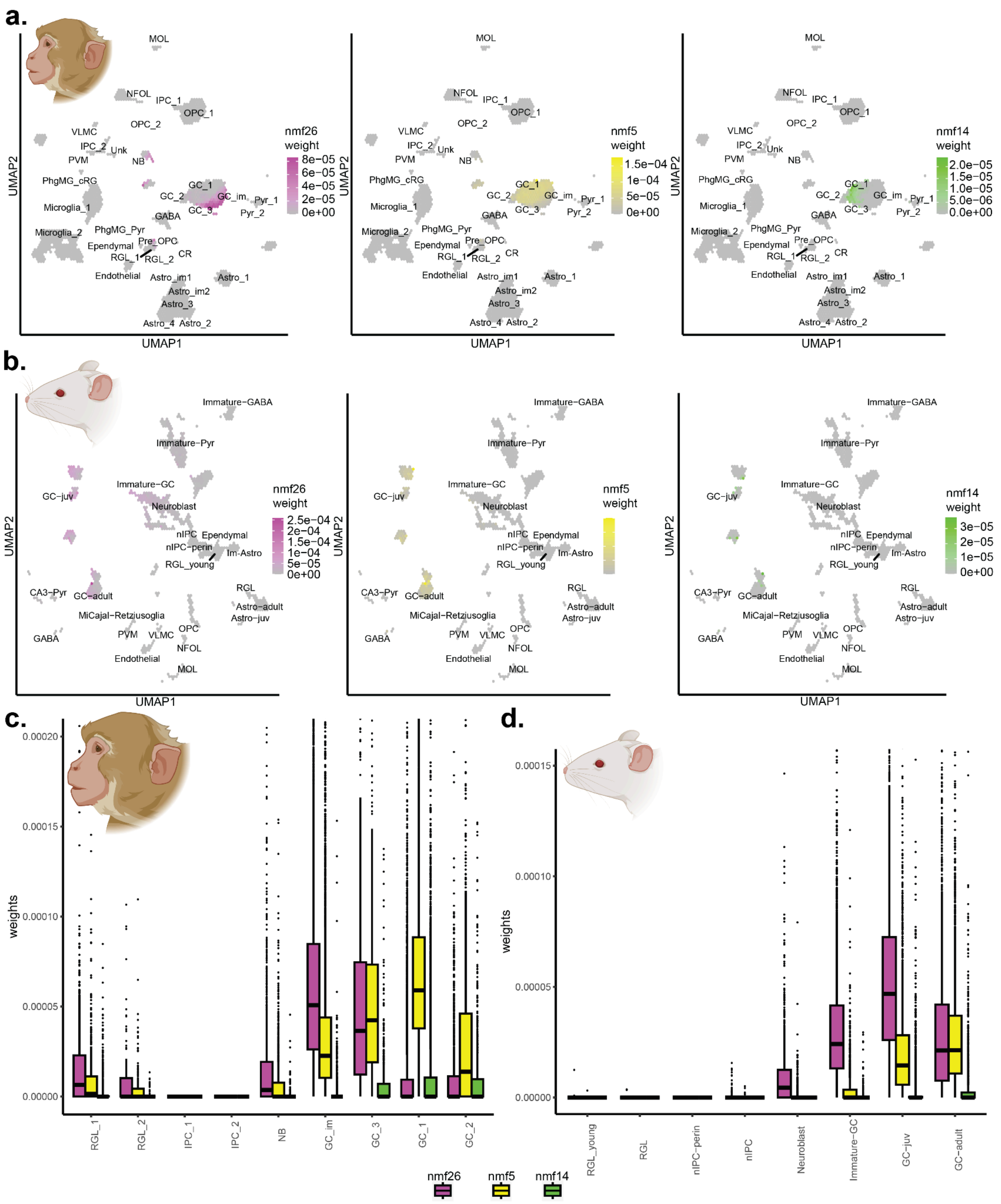


**Figure S23: Nmf-based transfer learning predicts a trajectory of maturation for macaque and mouse granule cells.** (**a**) UMAP of scRNA-seq dataset from adult macaque DG with cell type annotations as published [21]. Every 100 cells were binned into hexagons to prevent overplotting. Color scale indicates the median value of each given nmf pattern weight [59] for every hexagon. (**b**) UMAP of scRNA-seq dataset from embryonic and postnatal adult mouse DG with cell type annotations as published [20]. Every 100 cells were binned into hexagons to prevent overplotting. Color scale indicates the median value of each given nmf pattern weight for every hexagon. (**c**) Boxplots of nmf weight for cells versus adult macaque DG cell type annotation as published [21]. Each box is colored by their respective nmf pattern. (**d**) Boxplots of nmf weight for cells versus adult macaque DG cell type annotation as published. Each box is colored by their respective nmf pattern.

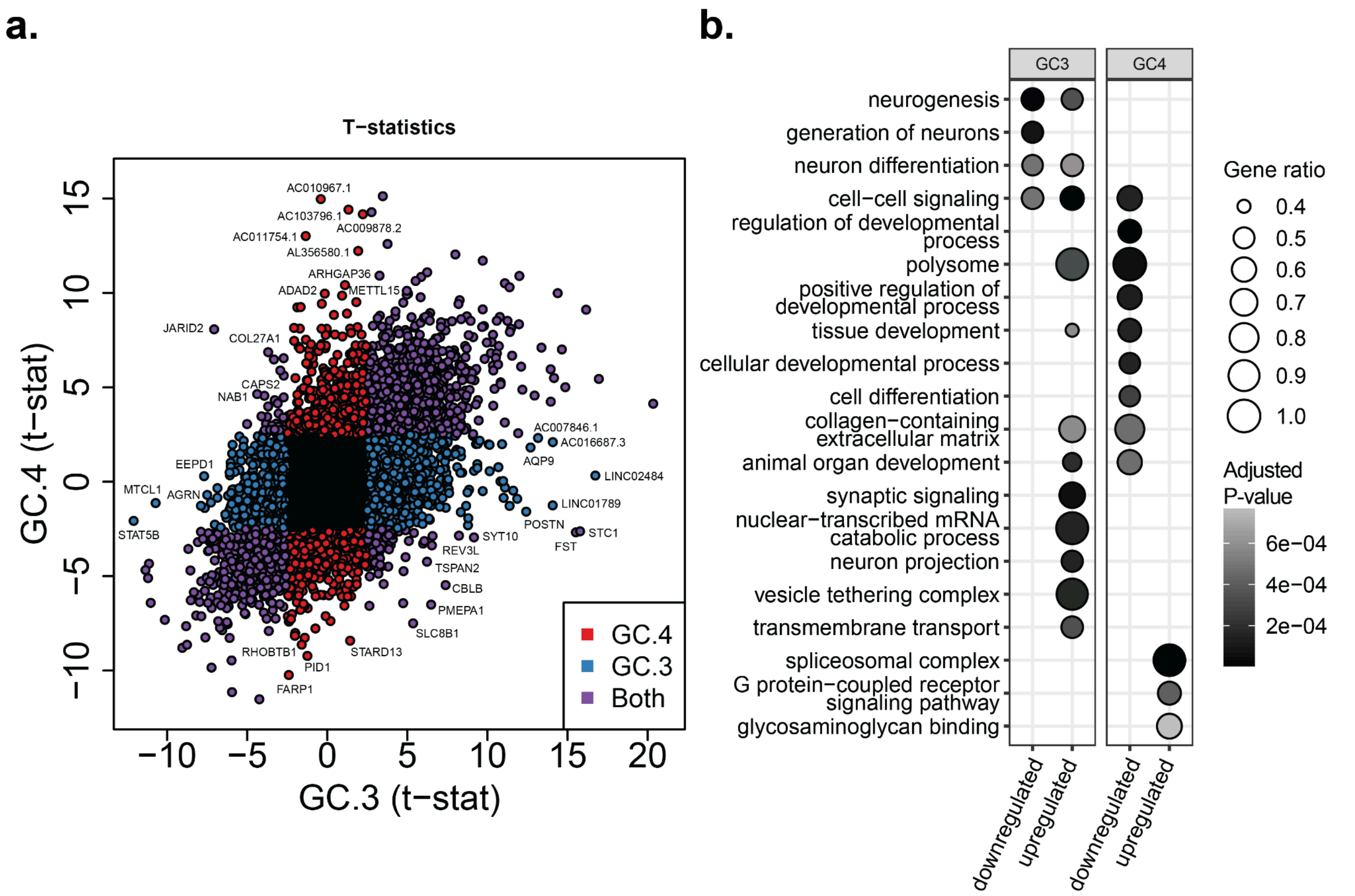


**Figure S24: Differentially enriched/depleted genes of immature and mature granule cells reveal functional differences.** (**a**) Scatterplot of t-statistics for GC.4 versus GC.3 (terminology from original publication) [59]. Each gene is colored by FDR < 0.05 for GC.4 (red), GC.3 (blue), or both (purple). (**b**) Dot plots for major gene ontology terms. Dot plots are faceted by significance and directionality for GC.3 or GC.4, respectively. Dot size represents the fraction of gene set in GO term that was differentially expressed (Gene ratio). Greyscale gradient represents adjusted *p*-value.

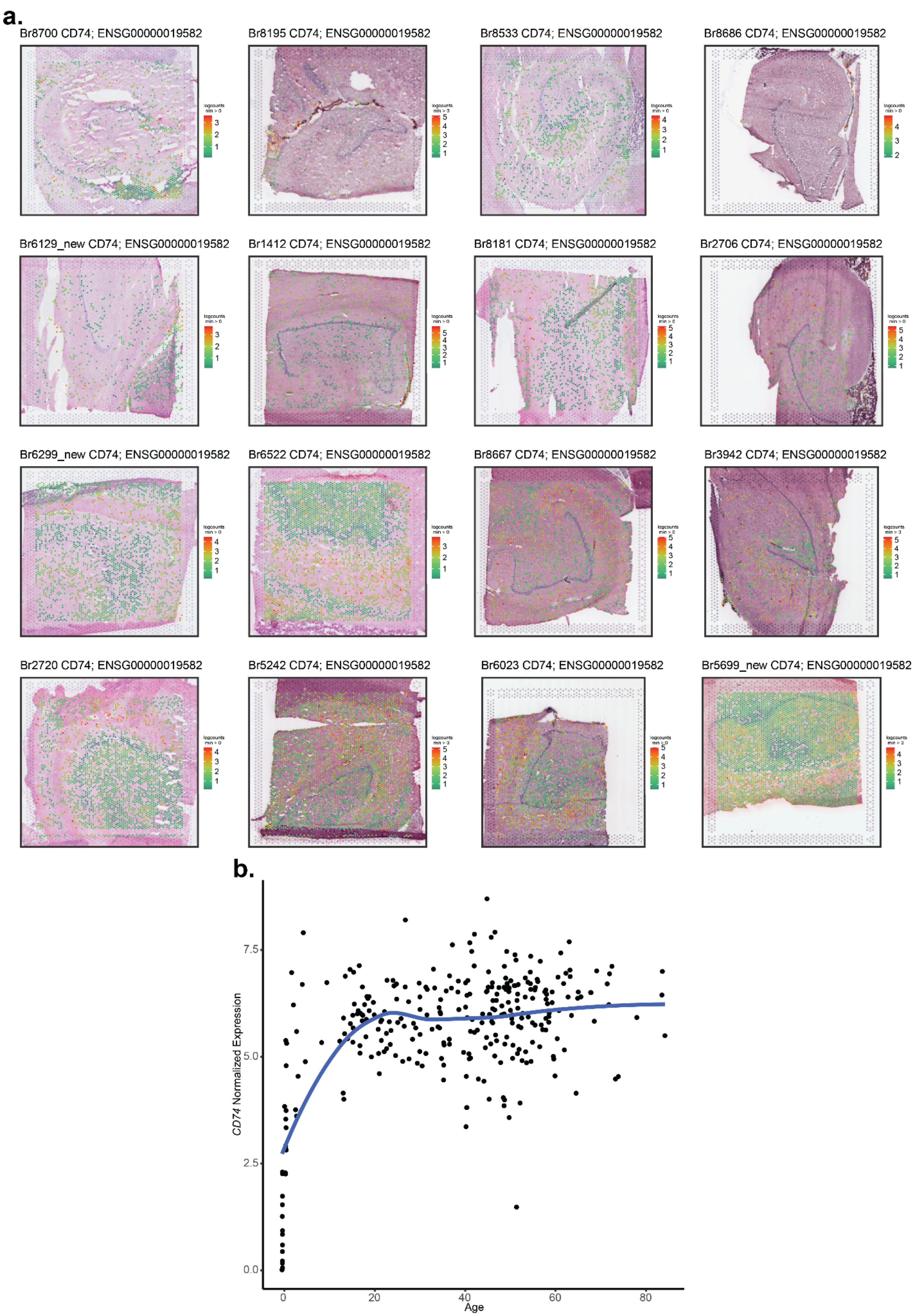
a.
Br8700 CD74; ENSG00000019582
Br8195 CD74; ENSG00000019582
Br8533 CD74; ENSG00000019582
Br8686 CD74; ENSG00000019582
Br6129_new CD74; ENSG00000019582
Br1412 CD74; ENSG00000019582
Br8181 CD74; ENSG00000019582
Br2706 CD74; ENSG00000019582
Br6299_new CD74; ENSG00000019582
Br6522 CD74; ENSG00000019582
Br8667 CD74; ENSG00000019582
Br3942 CD74; ENSG00000019582
Br2720 CD74; ENSG00000019582
Br5242 CD74; ENSG00000019582
Br6023 CD74; ENSG00000019582
Br5699_new CD74; ENSG00000019582
b.
CD74 Normalized Expression
0.0
2.5
5.0
7.5
0
20
40
60
80
Age

**Figure S25. *CD74* spatial expression across age.** (**a**) Data visualization of the Visium spots with the log normalized counts for *CD74* overlaid onto H&E images. The samples are ordered by age from left to right and top to bottom. (**b**) Scatter plot of $\log_2$ normalized expression (*y*-axis) and age (*x*-axis) with local regression line of reference bulk RNA-seq HPC data [51]. Each point is one donor sample from the neurotypical group of the reference dataset.

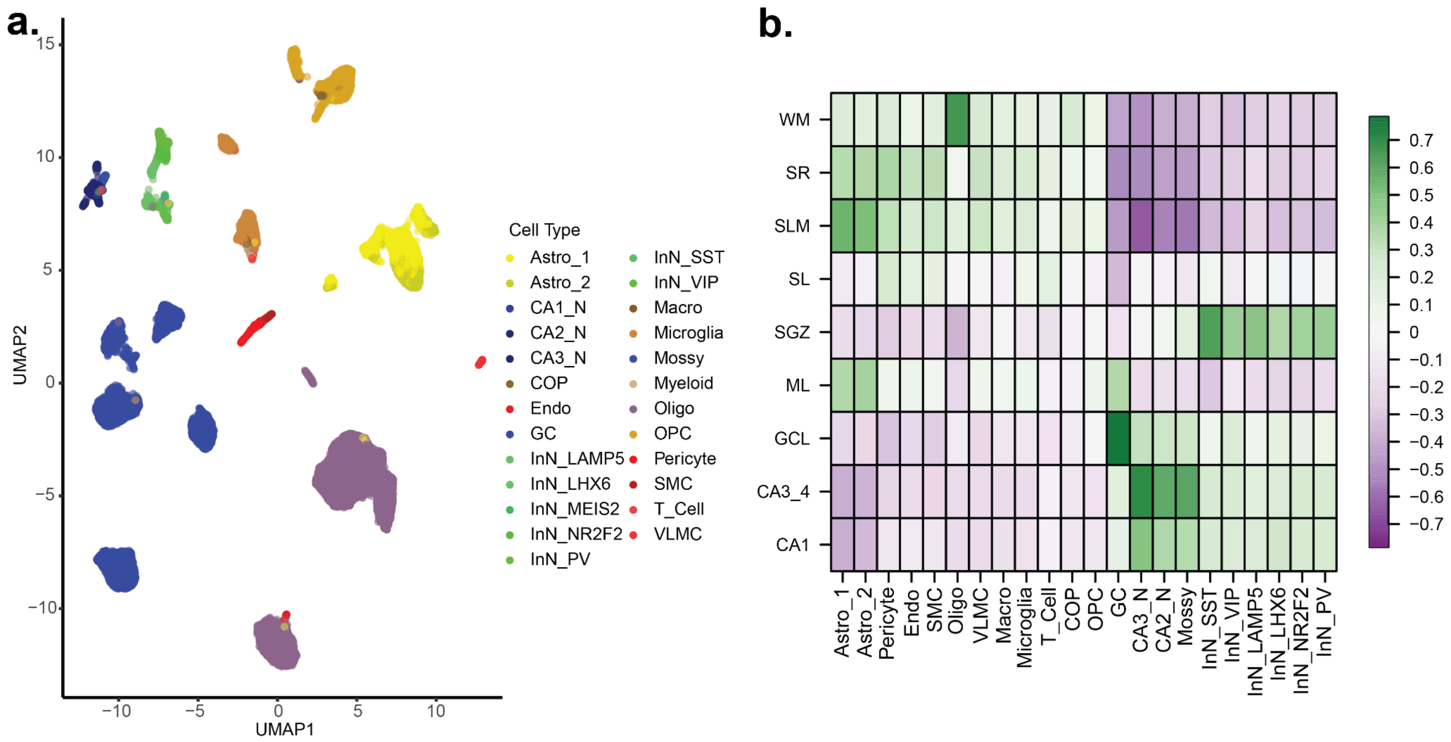


**Figure S26. Isolated dentate gyrus snRNA-seq nuclei from Franjic *et al.* 2022.** (**a**) Uniform Manifold Approximation and Projection (UMAP) representation of the single-nucleus RNA sequencing (snRNA-seq) dataset from Franjic *et al.* 2022 [31]. This dataset was truncated to include only dentate gyrus nuclei annotated by the original authors. Many cell subtypes were collapsed to more general types to improve the performance of cell type deconvolution, which is known to struggle with rare cell types. Points are colored by cell type labels. (**b**) Heatmap of Pearson correlation for enrichment *t*-statistics of pseudo-bulked cell types from Franjic, *et. al.*, 2022 (*x*-axis) and HPC spatial domains from this study (*y*-axis).

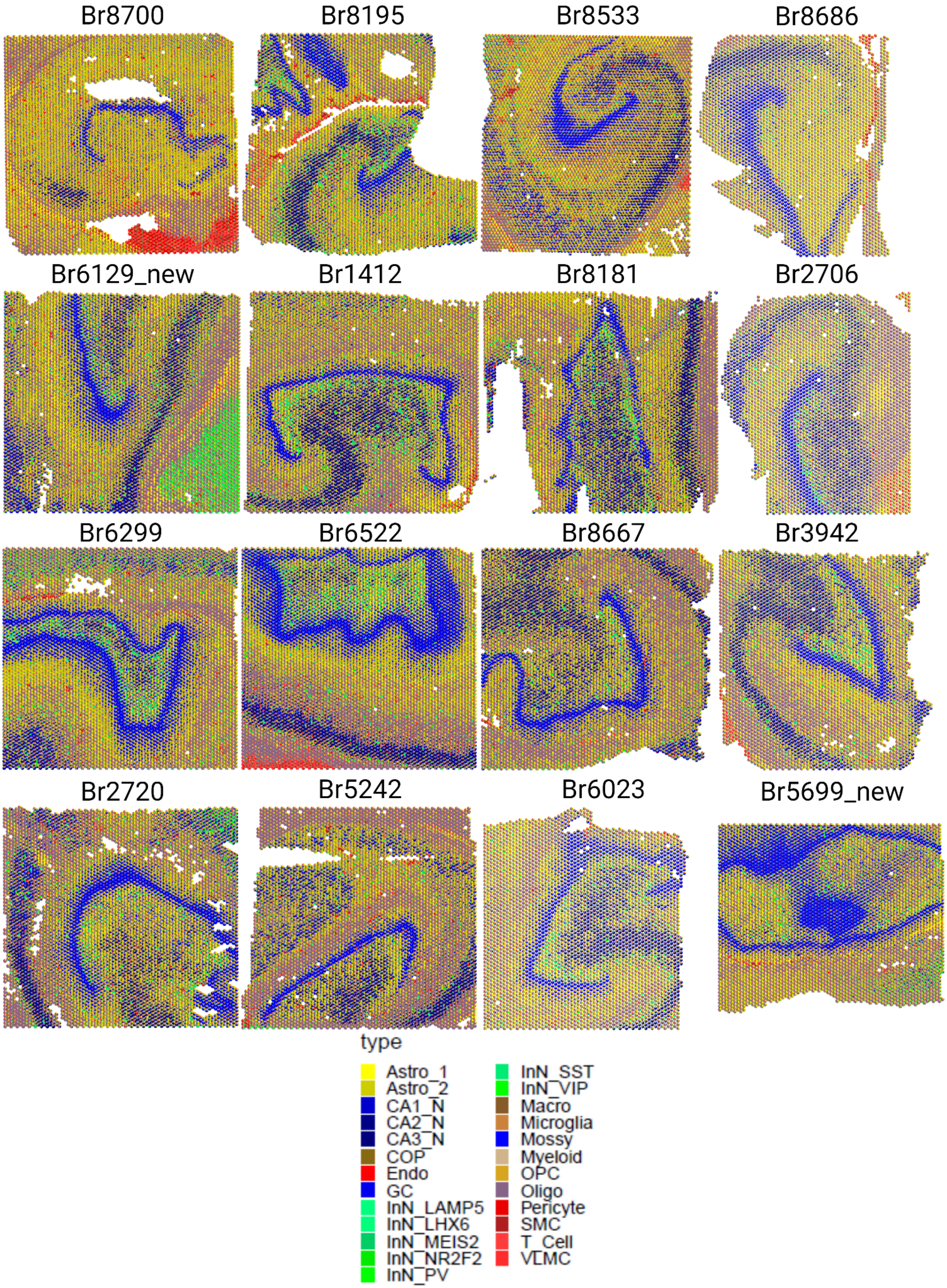

Br8700
Br8195
Br8533
Br8686
Br6129_new
Br1412
Br8181
Br2706
Br6299
Br6522
Br8667
Br3942
Br2720
Br5242
Br6023
Br5699_new
type
Astro_1
Astro_2
CA1_N
CA2_N
CA3_N
COP
Endo
GC
InN_LAMP5
InN_LHX6
InN_MEIS2
InN_NR2F2
InN_PV
InN_SST
InN_VIP
Macro
Microglia
Mossy
Myeloid
OPC
Oligo
Pericyte
SMC
T_Cell
VLMC

**Figure S27. Scatterpie plots of cell proportions derived from `cell2location` cell type deconvolution.** Spot-level data visualization for all *N*=16 capture areas where each Visium spot is a pie chart representing the relative abundance or proportion of cell types within each spot. Colors represent the proportions of cell type derived from mean cell abundances estimated by `cell2location` after performing spot-level deconvolution with Franjic *et al.* 2022 snRNA-seq data [31].

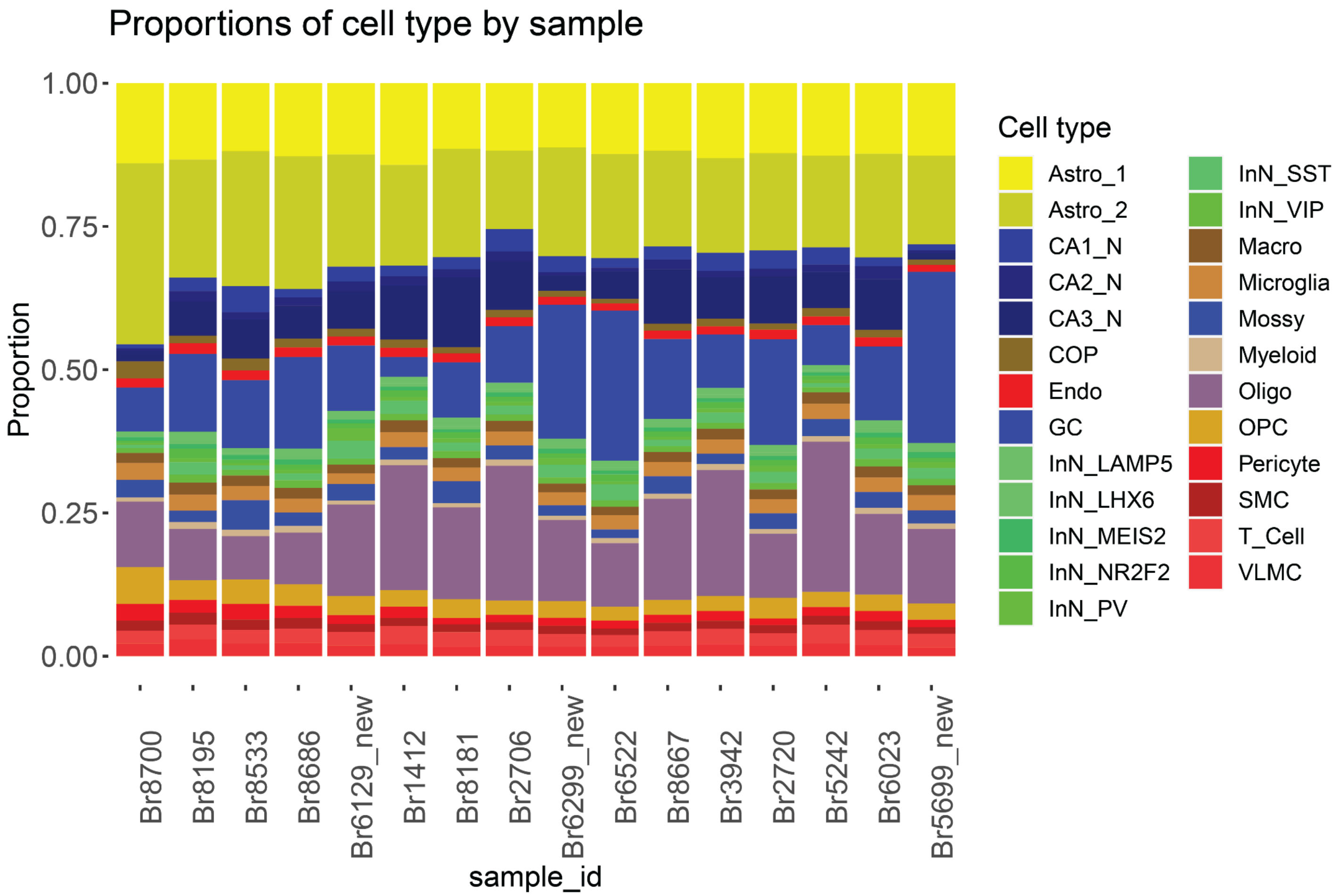


**Figure S28**. **Relative abundance or proportion of cell types derived from `cell2location`.** Barplot of cell type proportions (ranging from 0 to 1) (*y*-axis) derived from mean cell abundances estimated by `cell2location` across the entire capture area for all samples ordered from left to right by age (*x*-axis).

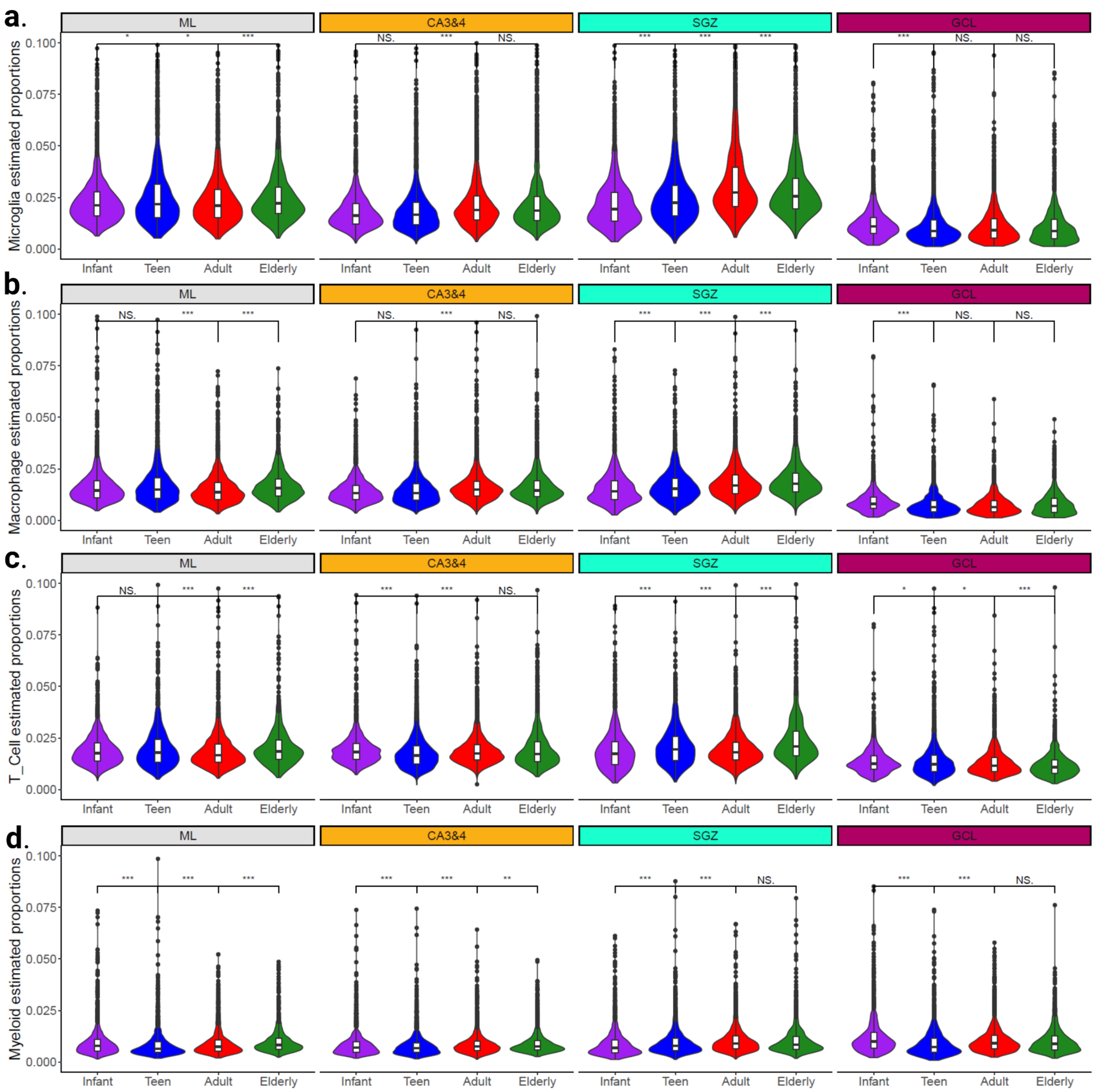


**Figure S29. Estimated proportions of select immune cell types.** (**a-d**) Violin plots superimposed with box plots for estimated cell proportions per Visium spot predicted by cell type deconvolution from `cell2location`, faceted by spatial domains corresponding to the DG layers ML, GCL, SGZ, and CA3&4. The *x*-axis is age groups, the *y*-axis is the estimated proportions. A two-sided Wilcoxon rank sum test was used to assess the statistical significance between pairs of age groups.

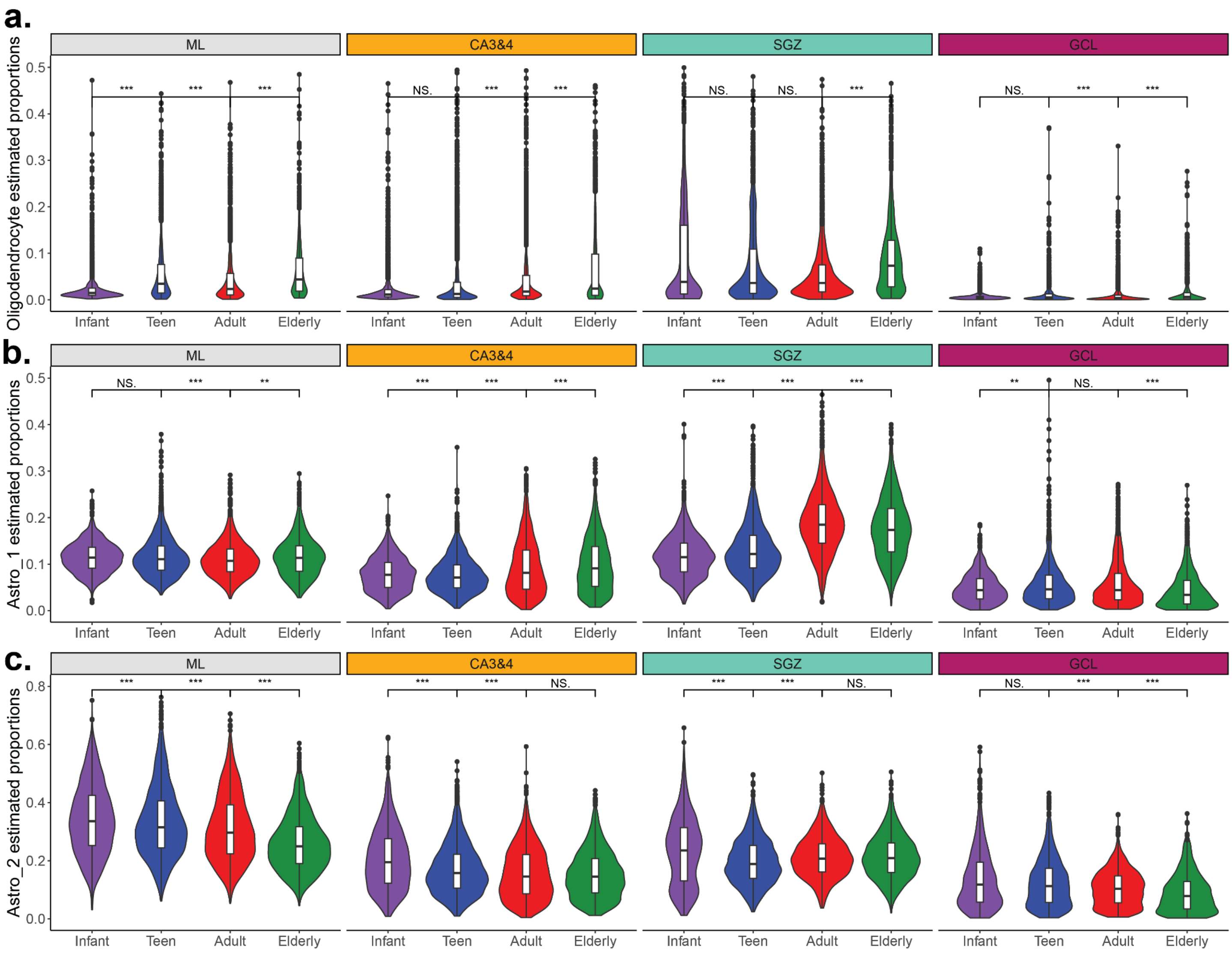


**Figure S30. Estimated proportions of oligodendrocytes and astrocyte subtypes.** (**a-d**) Violin plots superimposed with box plots for estimated cell proportions per Visium spot predicted by cell type deconvolution from `cell2location`, faceted by spatial domains corresponding to the DG layers ML, GCL, SGZ, and CA3&4. The *x*-axis is the age groups, the *y*-axis is the estimated proportions. A two-sided Wilcoxon rank sum test was used to assess the statistical significance between pairs of age groups. The rows represent four different cell types: (**a**) oligodendrocytes, (**b**) astrocyte subtype 1 (Astro_1), and (**c**) astrocyte subtype 2 (Astro_2).

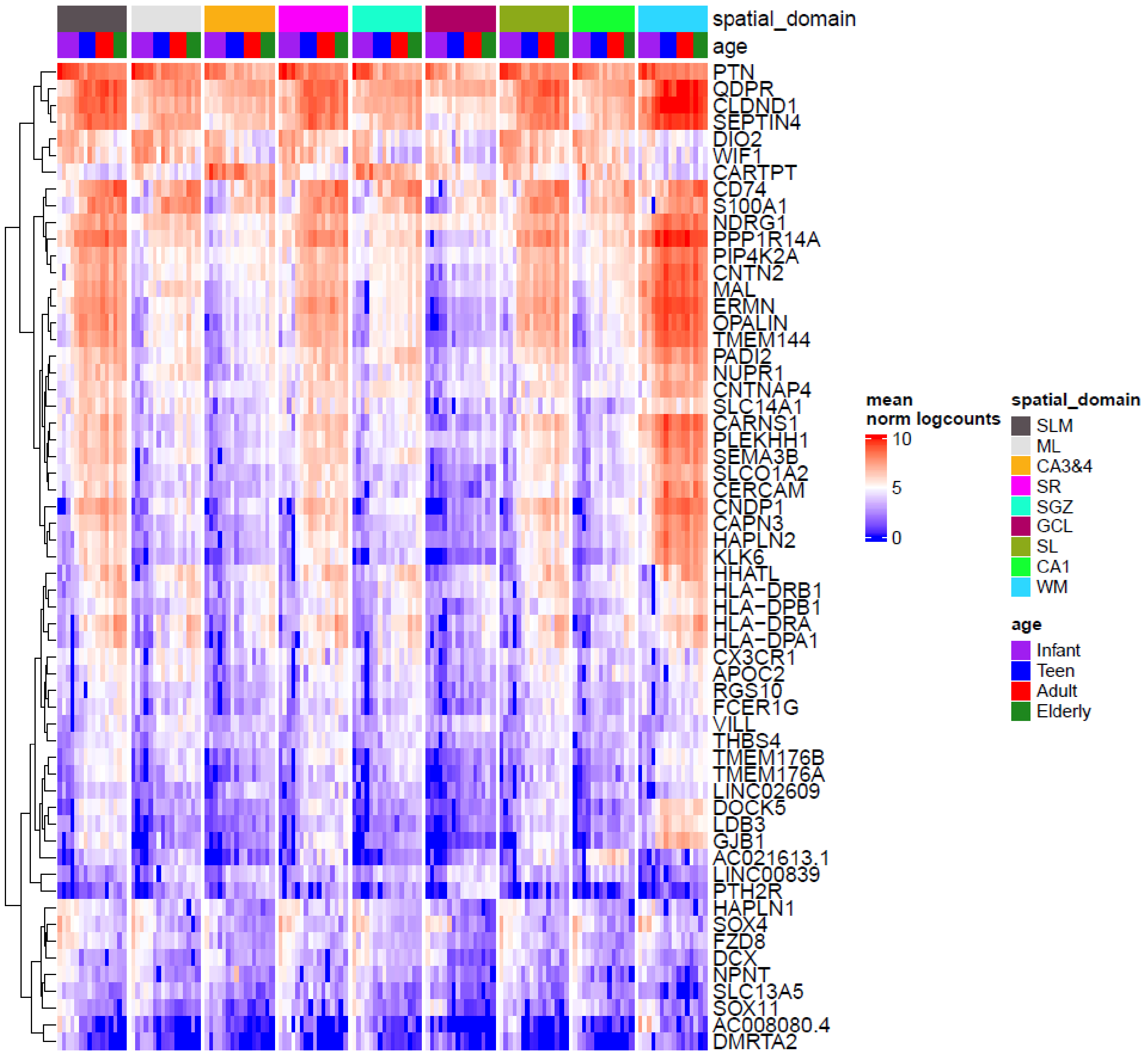


**Figure S31. The wide-spread aging signature throughout all pseudo-bulked spatial domains.** Heatmap of the mean log2 normalized counts for the wide-spread aging signature gene set. Hierarchical clustering was performed across rows. Columns are organized by spatial domains corresponding to HPC spatial domains. Labeled genes are those that show strong increasing age gradients across many spatial domains.

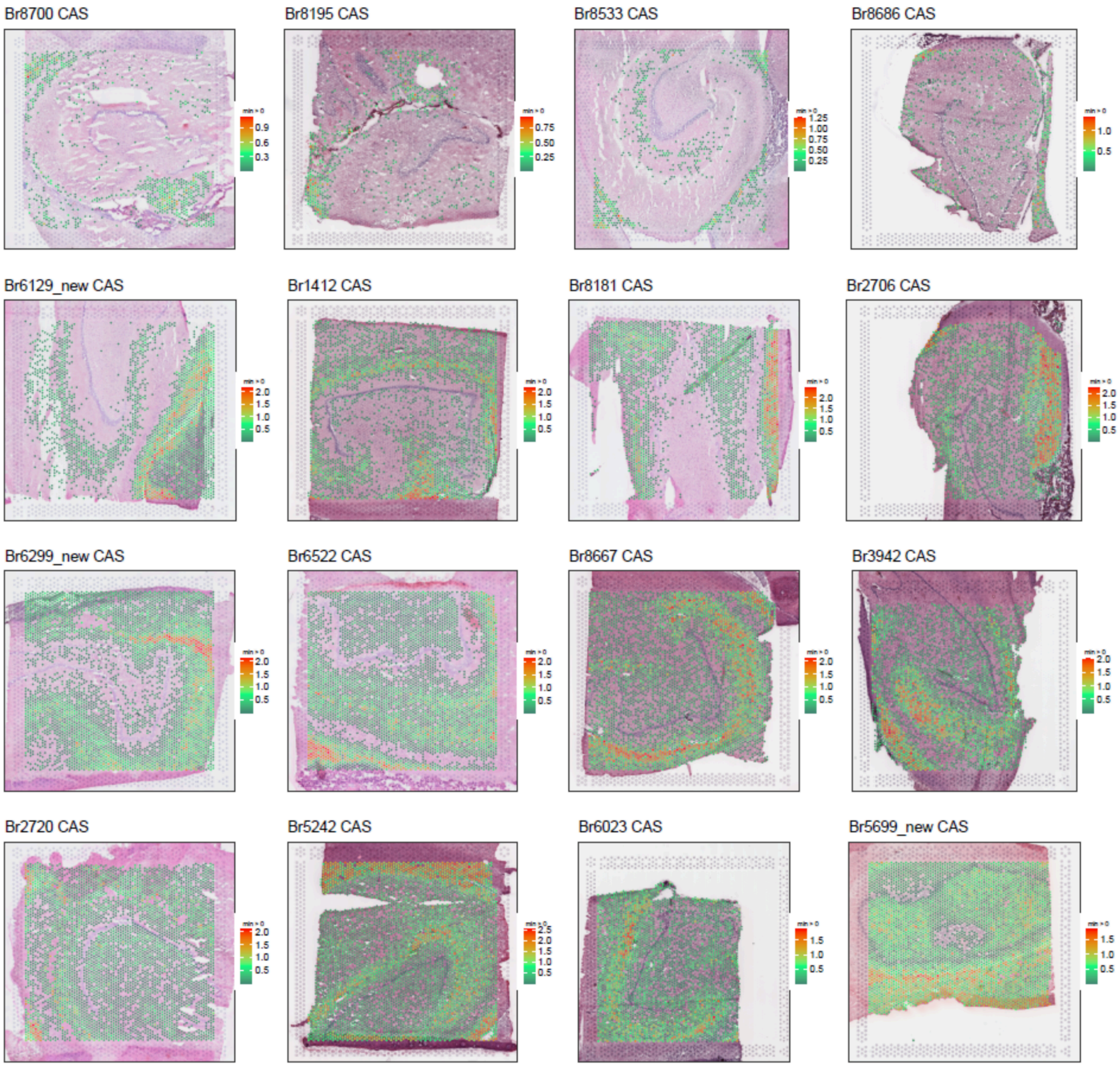


**Figure S32. Common aging score across age.** Data visualization of the Visium spots with common aging score (CAS) in arbitrary units overlaid onto H&E image. The samples are ordered by age from left to right and top to bottom.

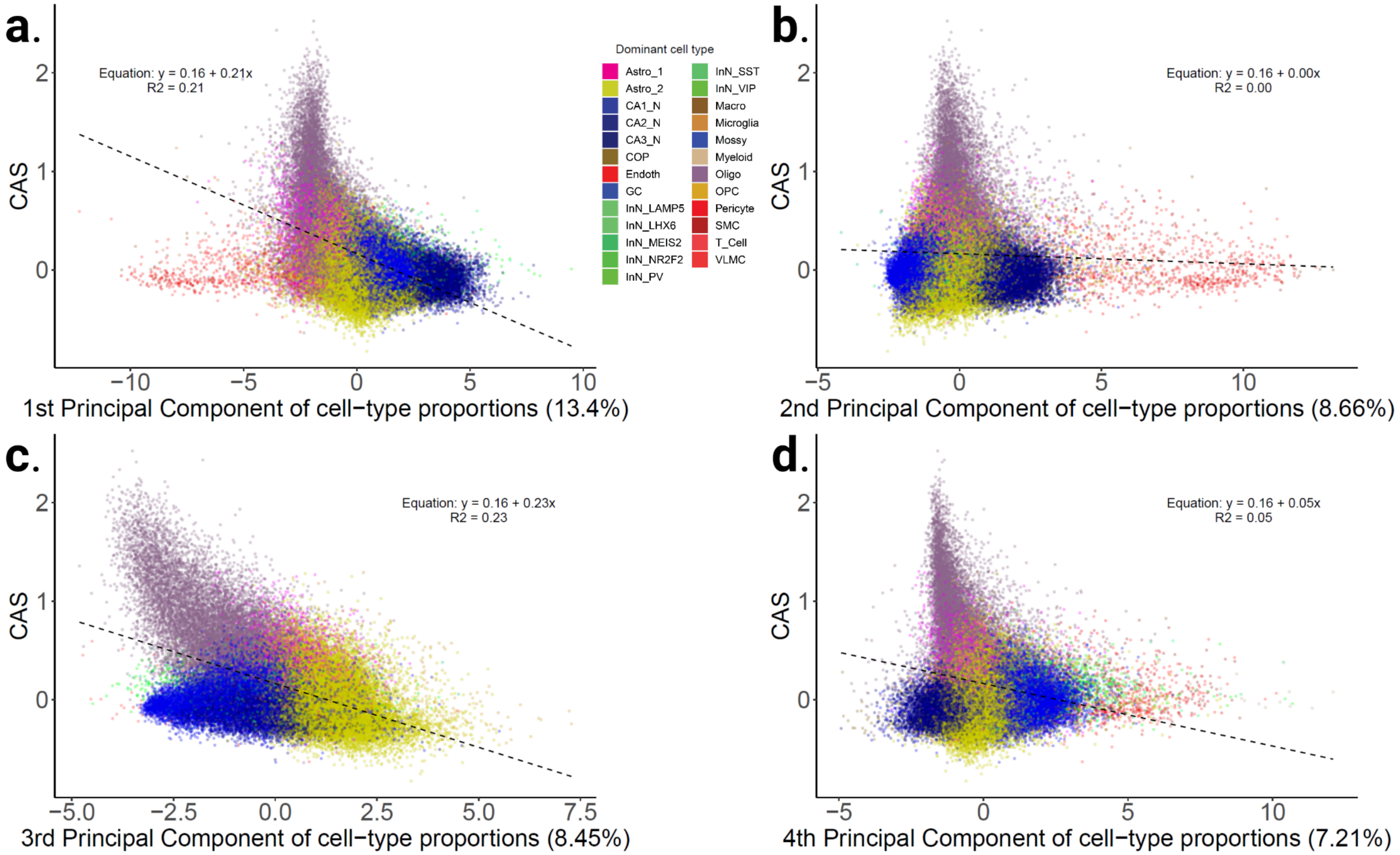


**Figure S33. Variance of cell type proportions alone does not predict CAS.** Using the estimated cell type proportions derived from `cell2location` for each spot, we performed PCA to obtain the first four PCs. (**a-d**) Scatter plots for CAS versus PCs 1-4 of cell type proportions derived from `cell2location`. Each point represents one spot and points are colored by the cell type with the highest estimated proportion for each spot.

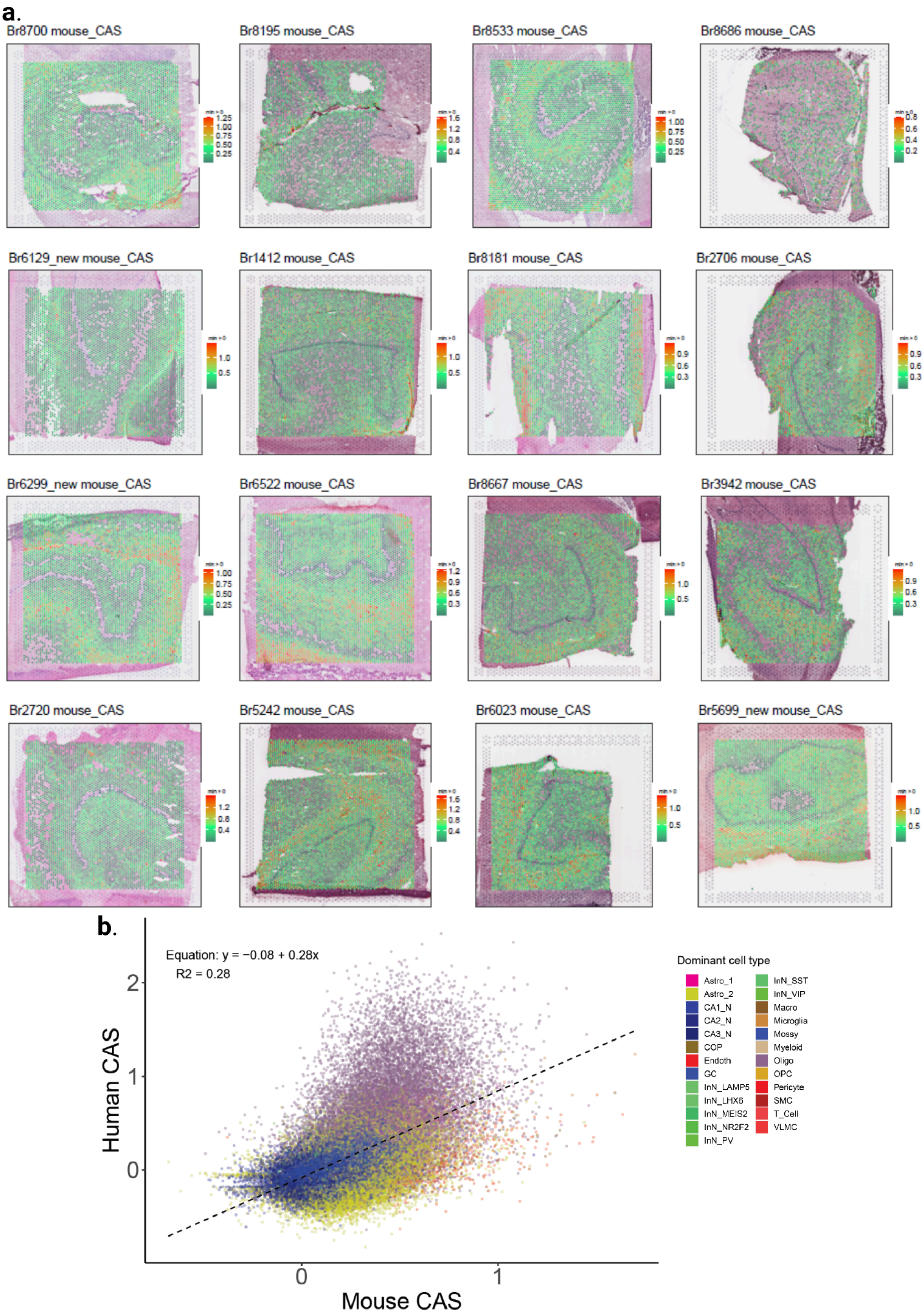
a.
Br8700 mouse_CAS
Br8195 mouse_CAS
Br8533 mouse_CAS
Br8686 mouse_CAS
Br6129_new mouse_CAS
Br1412 mouse_CAS
Br8181 mouse_CAS
Br2706 mouse_CAS
Br6299_new mouse_CAS
Br6522 mouse_CAS
Br8667 mouse_CAS
Br3942 mouse_CAS
Br2720 mouse_CAS
Br5242 mouse_CAS
Br6023 mouse_CAS
Br5699_new mouse_CAS
b.
Equation: y = −0.08 + 0.28x
R2 = 0.28
Human CAS
Mouse CAS
Dominant cell type
Astro_1
Astro_2
CA1_N
CA2_N
CA3_N
COP
Endoth
GC
InN_LAMP5
InN_LHX6
InN_MEIS2
InN_NR2F2
InN_PV
InN_SST
InN_VIP
Macro
Microglia
Mossy
Myeloid
Oligo
OPC
Pericyte
SMC
T_Cell
VLMC

**Figure S34. Visualization mouse of common aging score across age in human.** (**a**) Data visualization of the common aging score (CAS) derived from mouse whole-brain in arbitrary units overlaid on H&E image. The samples are ordered by age from left to right and top to bottom. (**b**) Scatter plot for human HPC CAS (**Figure 6**) versus mouse whole-brain CAS. Each point represents one spot and is colored by the cell type with the highest estimated proportion estimated by `cell2location`.
